# The evolution of the metazoan Toll receptor family and its expression during protostome development

**DOI:** 10.1101/2021.02.01.429095

**Authors:** Andrea Orús-Alcalde, Tsai-Ming Lu, Andreas Hejnol

## Abstract

**Background:** Toll-like receptors (TLRs) play a crucial role in immunity and development. They contain leucine-rich repeat domains, one transmembrane domain, and one Toll/IL-1 receptor domain. TLRs have been classified into V-type/scc and P-type/mcc TLRs, based on differences in the leucine-rich repeat domain region. Although TLRs are widespread in animals, detailed phylogenetic studies of this gene family are lacking. Here we aim to uncover TLR evolution by conducting a survey and a phylogenetic analysis in species across Bilateria. To discriminate between their role in development and immunity we furthermore analyzed stage-specific transcriptomes of the ecdysozoans *Priapulus caudatus* and *Hypsibius exemplaris*, and the spiralians *Crassostrea gigas* and *Terebratalia transversa*.

**Results:** We detected a low number of TLRs in ecdysozoan species, and multiple independent radiations within the Spiralia. V-type/scc and P-type/mcc type-receptors are present in cnidarians, protostomes and deuterostomes, and therefore they emerged early in TLR evolution, followed by a loss in xenacoelomorphs. Our phylogenetic analysis shows that TLRs cluster into three major clades: clade α is present in cnidarians, ecdysozoans, and spiralians; clade β in deuterostomes, ecdysozoans, and spiralians; and clade γ is only found in spiralians. Our stage-specific transcriptome and *in situ* hybridization analyses show that TLRs are expressed during development in all species analyzed, which indicates a broad role of TLRs during animal development.

**Conclusions:** Our findings suggest that the bilaterian TLRs likely emerged by duplication from a single TLR encoding gene (*proto*-TLR) present in the last common cnidarian-bilaterian ancestor. This *proto*-TLR gene duplicated before the split of protostomes and deuterostomes; a second duplication occurred in the lineage to the Trochozoa. While all three clades further radiated in several spiralian lineages, specific TLRs clades have been presumably lost in others. Furthermore, the expression of the majority of these genes during protostome ontogeny suggests their involvement in immunity and development.

## Background

Toll-like receptors (TLRs) are involved in immunity and development in metazoans [1–7]. The first described *Tlr* was the *Drosophila* gene *Toll*, which plays a role during early embryonic development [8, 9] and in immunity [10]. The human toll receptor TLR4 was the first TLR discovered in mammals [11]. Since then, TLRs have been found in most planulozoans (Cnidaria + Bilateria) [12–14]. Both in vertebrates and invertebrates, these receptors recognize pathogens and activate the Toll pathway, which induces the expression of downstream immune genes [15–17]. In *Drosophila*, TLRs are mainly activated by gram-positive bacteria, fungi, and viruses, promoting the synthesis of antimicrobial peptides (AMPs) [4, 10, 17–21]. In vertebrates, TLRs are involved in innate immunity and in the activation and regulation of adaptive immunity [11, 22–26]. TLRs are also involved in the immunity of other animals such as cnidarians [27], mollusks [28–31], annelids [32, 33], crustaceans [34] and echinoderms [35]. The developmental roles of TLRs in *Drosophila* [reviewed in 2] comprise the establishment of the dorso-ventral axis [8, 9], segmentation [36], muscle and neuronal development [37, 38], wing formation [39, 40] and heart formation [41]. TLRs also play a role in cnidarian development [27]. Moreover, in spiralians, TLRs are expressed during the development of mollusks [31] and annelids [32], but no further analyses have been conducted. TLRs are also involved in nervous system development in mice [42– 45], although the ligands that activate them during this process remain unknown [2].

TLRs are proteins characterized by an extracellular region containing one or more leucine-rich repeat (LRR) domains, one type-I transmembrane domain and one intracellular Toll/IL-1 receptor (TIR) domain (Figure 1) [46, 47]. The extracellular LRR domains are the regions that recognize the ligand [48, 49]. Each LRR domain is constituted by 22-26 amino acids, in which multiple leucine residues are present [46]. Some LRR domains contain cysteine residues in the N-terminal (LRRNT) or the C-terminal (LRRCT) part of the LRR domain [6, 47, 50]. However, LRR domains are also found in a large number of other proteins [51], for example in the immune NOD receptors [52] and in proteins involved in developmental processes (e.g. Slit, Capricious, Tartan) [53, 54]. The TIR domain is involved in signal transduction [47] and is also present in other proteins, e.g. in immune proteins in plants [55, 56], in members of the interleukin-I receptor family (IL-1) [47, 57] and in adaptors of the Toll pathway (e.g. MyD88) [58–60]. Although the TIR domain is the most characteristic domain of the TLRs, at least one LRR domain must be present to categorize a receptor as TLR (Figure 1) [13].

**Figure 1.**
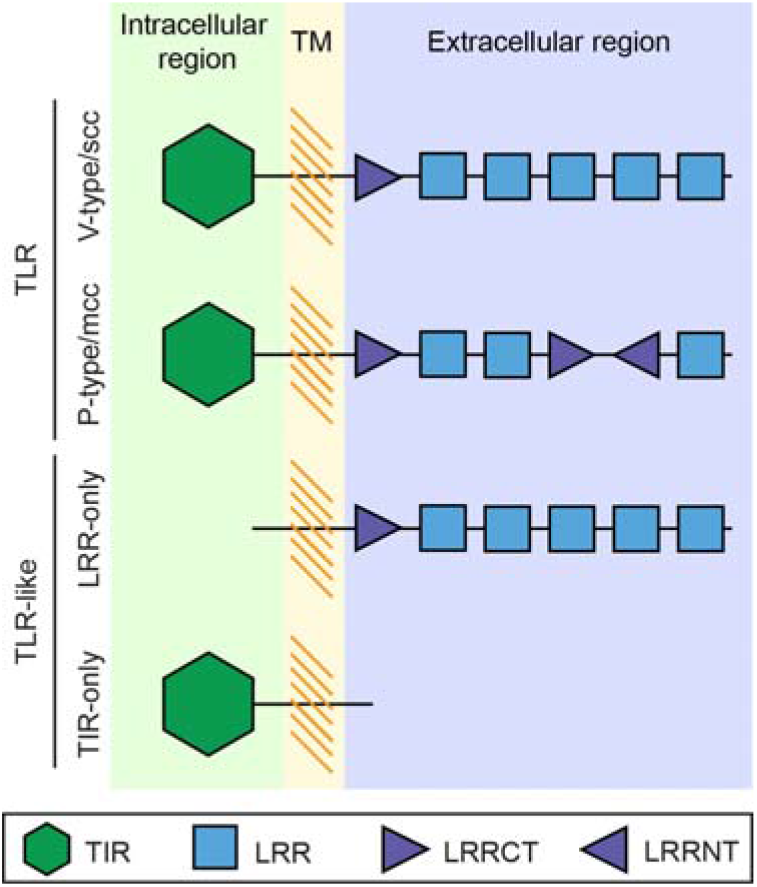
Structure of TLR and TLR-like receptors. TLRs are constituted by a series of extracellular leucine-rich repeat (LRR) domains, a transmembrane region (TM) and an intracellular Toll/IL-1 receptor (TIR) domain. TLRs are often classified into V-type/scc or P-type/mcc according to the structure of their extracellular region. V-type/scc TLRs have only one LRRCT located next to the TIR domain, while P-type/mcc TLRs have more than one LRRCT and, sometimes, an LRRNT domain. Proteins that lack either the LRR domains or the TIR domain are not considered as TLR receptors. These TLR-like proteins are classified in LRR-only or TIR-only. [Adapted from 7, 13]

Based on the structure of the LRR domains, TLRs have been previously classified as vertebrate-type or single cysteine cluster (V-type/scc), and protostome-type or multiple cysteine cluster (P-type/mcc) (Figure 1) [7, 13, 61, 62]. V-type/scc TLRs are characterized by having only one LRRCT domain, which is located next to the cellular membrane. P-type/mcc TLRs contain at least two LRRCT domains and, commonly, an LRRNT [7, 13]. Traditionally, it has been assumed that all deuterostome TLRs belong to the V-type/scc [62], and because *Drosophila melanogaster* TLRs (except for Toll9) and the *Caenorhabditis elegans* TLR belong to the P-type/mcc, they have been suggested to be protostome specific [62]. However, P-type TLR are also present in invertebrate deuterostomes and V-type TLRs in protostomes [13, 14, 63, 64]. Therefore, in agreement with Davidson et al., 2008 [63]; and Halanych and Kocot, 2014 [64], we affirm that the V-P-type nomenclature is problematic and should be avoided in favor of the mcc/scc nomenclature.

Several authors consider that TLRs originated in the lineage to the Planulozoa by the fusion of a gene with a TIR domain and a gene containing only LRR domains [7, 14, 65]. Such proteins, named TLR-like proteins (Figure 1), are involved in immunity [7, 12–14, 66–71] – e.g. in *Hydra*, association of LRR-only and TIR-only proteins activates the Toll pathway [72, 73].

The TLR complement has been previously surveyed in vertebrates [11, 50, 74–76] and in a few invertebrates, especially in arthropods [8, 14, 18, 77, 78]. Humans have 10 TLRs [11, 50], *D. melanogaster* has 9 [8, 18] and the nematode *C. elegans* has only one [79]. Recent genome and transcriptome sequencing of more organisms has revealed that TLRs are widespread across the metazoan tree (summary in Figure 2). Outside bilaterians, TLRs are present in anthozoan cnidarians (e.g. *Nematostella* [27], *Acropora* [69], *Orbicella* [80]), but not in hydrozoans (e.g. *Hydra* [72], *Clytia* [81]). Furthermore, TLRs have not been found in ctenophores [82, 83], placozoans [70] and poriferans [66, 71]. Within bilaterians, previous studies have shown that the number of TLRs in spiralians is highly variable between species [63, 64, 84–87], suggesting that TLR genes underwent several independent radiations [13, 63, 86, 88]. However, the surveyed platyhelminth and rotifer species lack TLRs [67, 68, 89]. In ecdysozoans, besides arthropods and nematodes, TLRs are also present in onychophorans, tardigrades, nematomorphs and priapulids [90]. In invertebrate deuterostomes, the number of TLRs in echinoderms and amphioxus is expanded [62, 91, 92], which is in contrast to the limited number of TLRs in tunicates [93, 94]. Although the TLR sequences of many metazoans have been explored [7, 12–14], more protostome species must be surveyed to gain a better picture of the TLR evolution (Figure 2).

**Figure 2.**
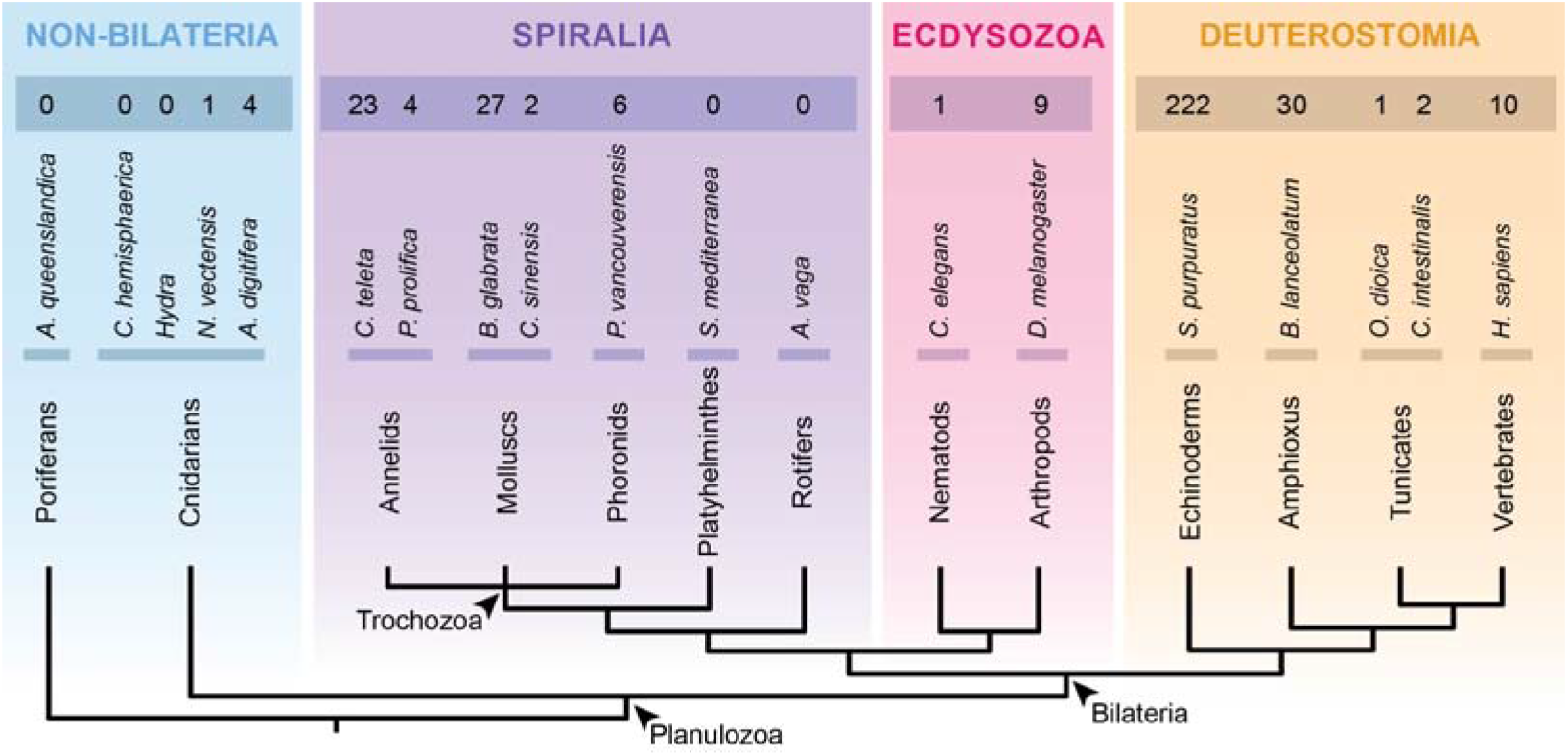
Summary of the number of TLRs across metazoans. No TLRs have been found outside Cnidaria and Bilateria. Spiralians show a variable number of TLRs, being, for example, 23 TLRs in the annelid *C. teleta*, but none in the rotifer *A. vaga*. In ecdysozoans, *C. elegans* and *D. melanogaster* have 1 and 9 TLRs, respectively. The number of TLRs in deuterostomes is also variable, being high in *S. purpuratus* and *B. lanceolatum*, but reduced in tunicates. Phylogeny according to [95].

Although the phylogenetic relationships of TLRs have been previously analyzed, these were mainly focused on vertebrate TLR evolution [65, 96] or including only few protostome species [13, 63, 86]. So far, the results are contradictory and are not sufficient to comprehend the detailed evolution of TLRs. For instance, Davidson et al., 2008 [63] suggested that TLRs are divided into three major clades, although the relationships between them remained unresolved. Brennan and Gilmore, 2018 [13] suggested that TLRs cluster according to the TLR-type (P-type/mcc or V-type/scc) and Liu et al., 2020 [65] suggested that both TLR types would be widespread in invertebrates. Furthermore, Luo et al., 2018 [86] showed lineage-specific expansions of TLRs in some trochozoan groups (phoronids, nemerteans and brachiopods). Thus, phylogenetic analyses including TLRs of species representing the broad metazoan diversity are lacking. In this study, we aim to reconstruct the TLR evolution by searching for TLRs in under-represented metazoan clades and performing a phylogenetic analysis including TLRs of species from the four main metazoan clades (cnidarians, spiralians, ecdysozoans and deuterostomes). Moreover, we aim to reconstruct the early TLR function by analyzing their expression during the course of development in four protostome species.

## Results

Our genome and transcriptomic surveys revealed a total of 198 TLRs in 25 species (Table 1, Figure 3). No TLRs were found in 20 species. Additionally, our analysis also revealed a large number of TLR-like proteins (TIR-only or LRR-only). However, only sequences containing a TIR domain, a transmembrane domain and, at least, one LRR domain were considered as criteria for TLRs.

**Table 1.**
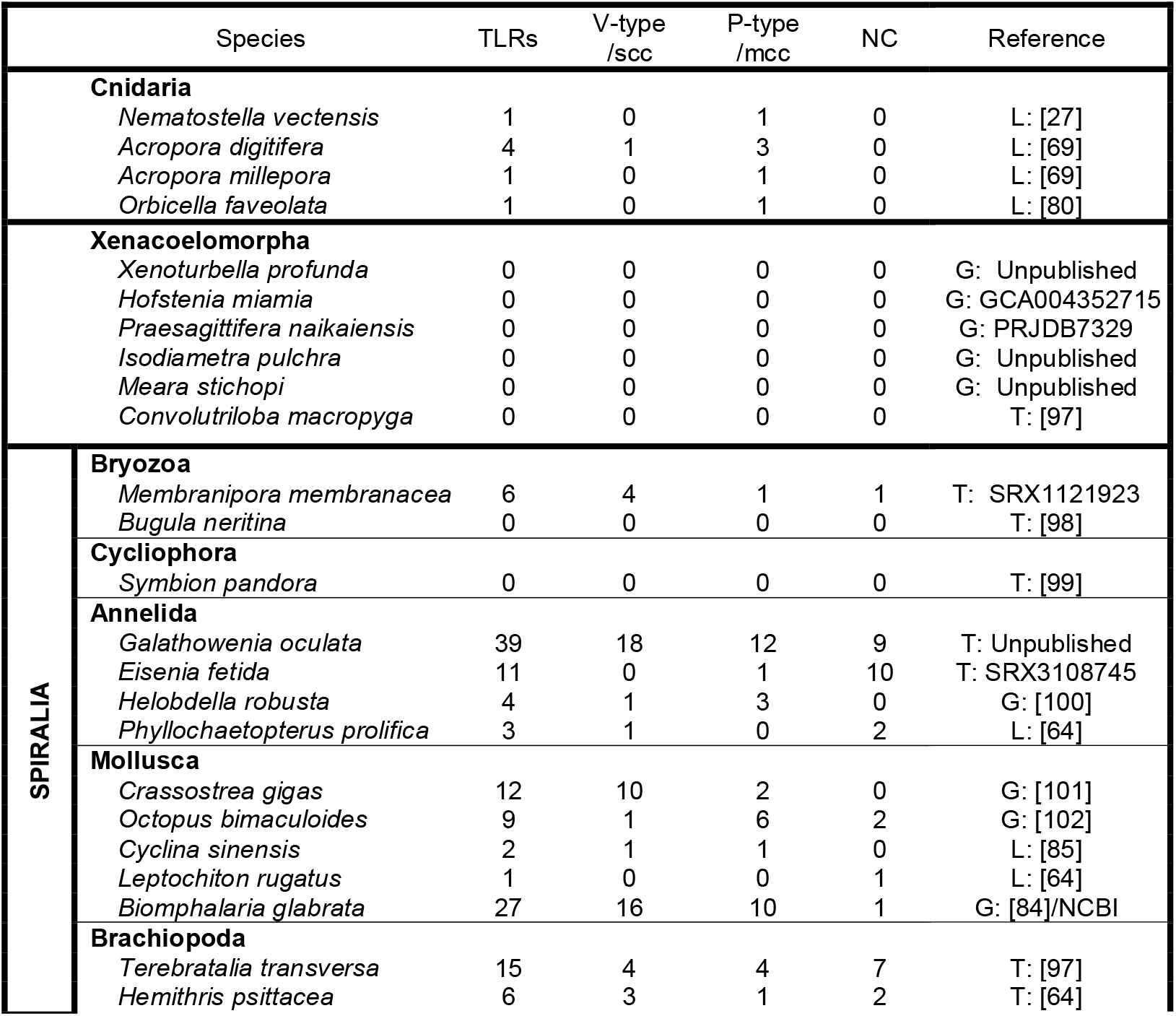
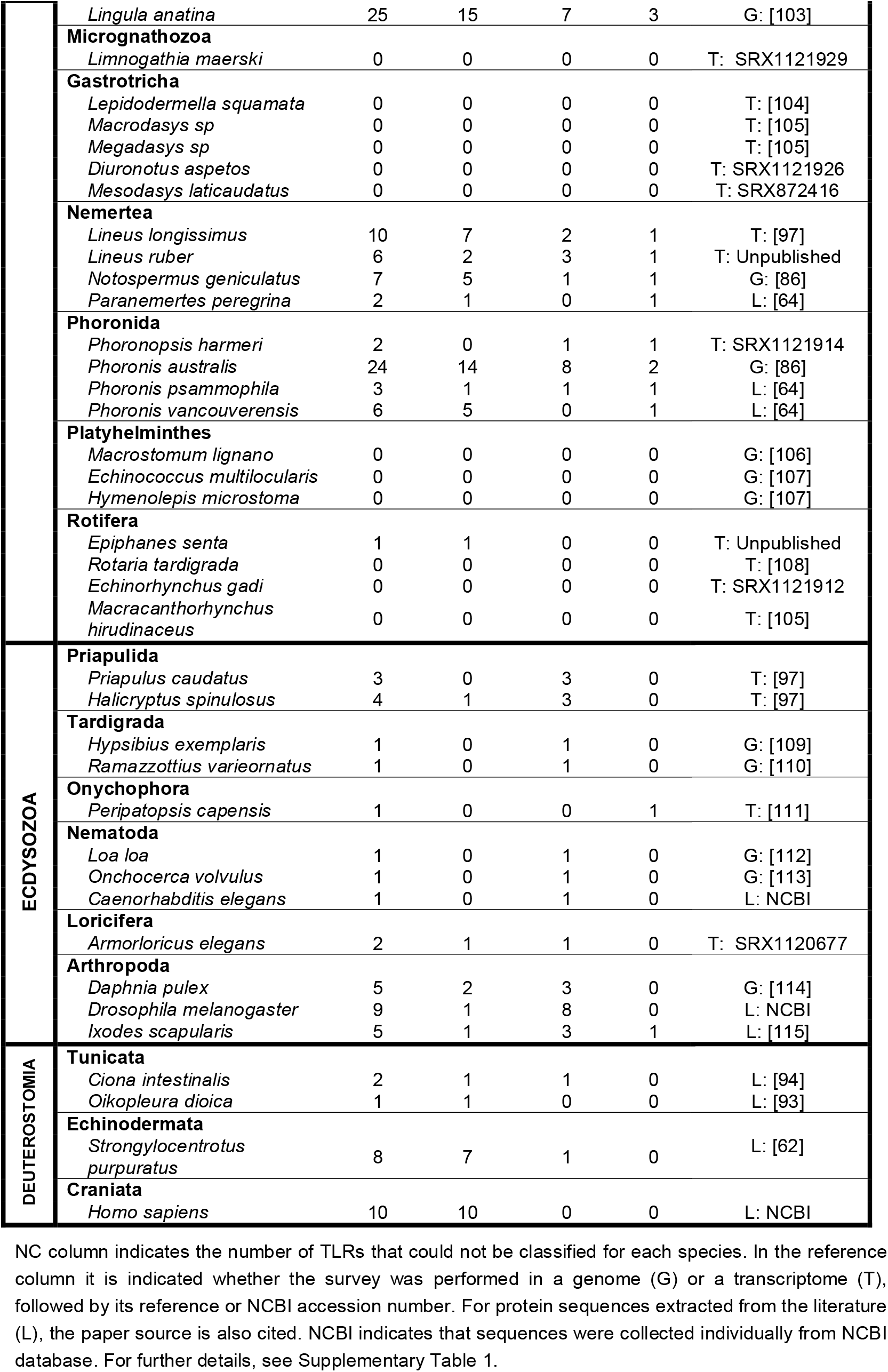
TLR genome/transcriptome survey results and classification of TLRs included in the phylogenetic analysis.

**Figure 3.**
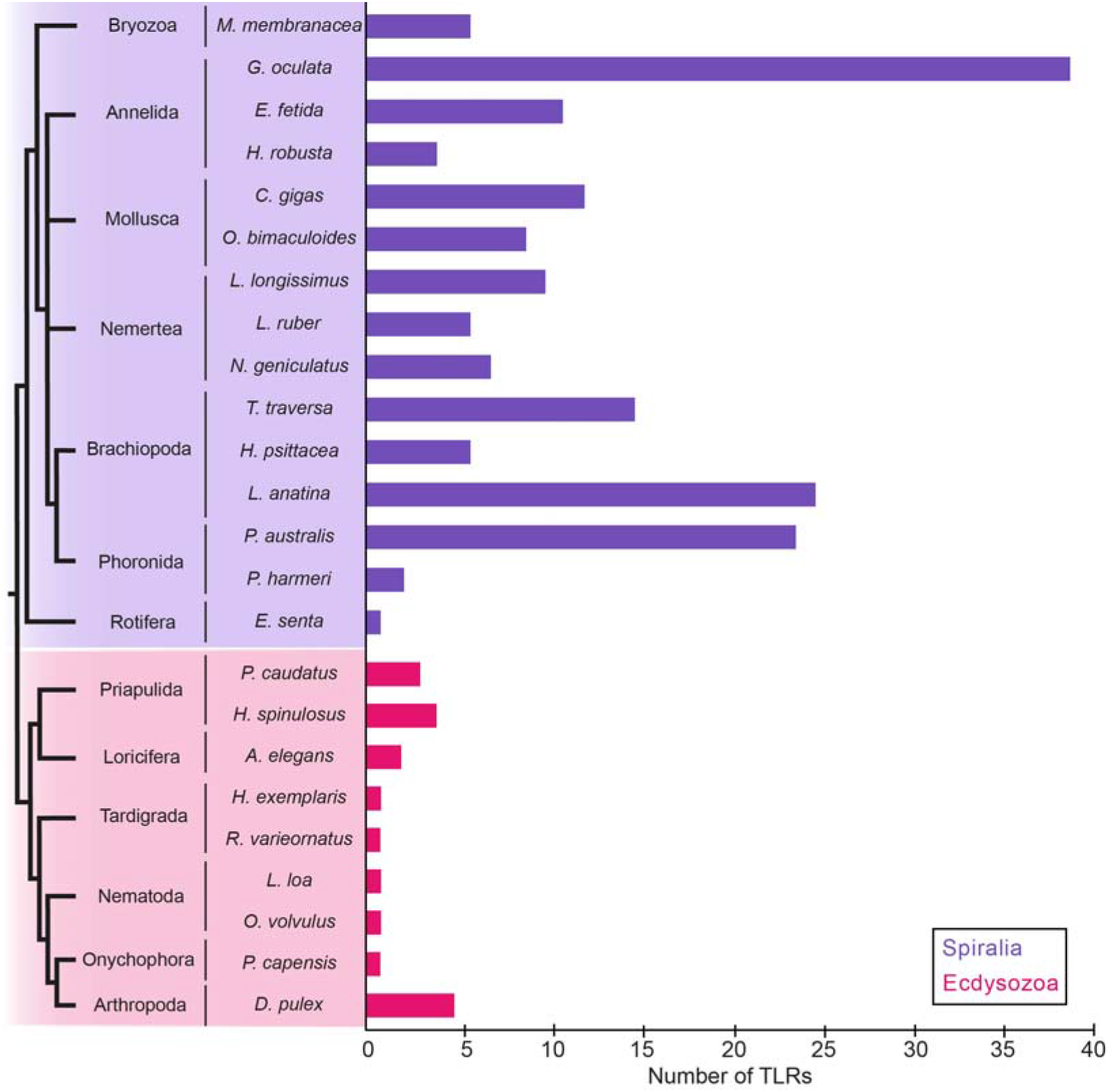
Total number of TLRs in the analyzed species. In general, the number of TLRs in spiralians (purple) is higher and more variable between species when compared to ecdysozoans (magenta). Species in which TLRs were not detected are excluded from the graph.

### TLRs are absent in the genomes and transcriptomes of xenacoelomorphs and in some spiralians

Our surveys revealed that TLRs are absent in the genomes and transcriptomes of all Xenacoelomorpha, Platyhelminthes, Cycliophora, Micrognathozoa and Gastrotricha species analyzed (Table 1). Furthermore, TLRs are also absent in the transcriptomes of all the rotifer species investigated, except for *E. senta* (Table 1, Figure 3). Moreover, although TLRs were present in the bryozoan *M. membranacea*, they were not found in the transcriptome of the bryozoan *B. neritina*. However, although TLRs were not detected, TLR-like proteins were present in all these animal groups (data not shown).

### The number of TLRs detected in members of Ecdysozoa is low when compared to Spiralia and Deuterostomia

The TLR survey of the ecdysozoan genomes and transcriptomes revealed only one TLR for the tardigrade, nematode, and onychophoran species analyzed (Table 1, Figure 3). Furthermore, we detected up to 4 different TLRs in priapulids, 2 in loriciferans, and 5 in arthropods.

### Multiple TLRs are detected in trochozoan species

TLRs were found in the genomes/transcriptomes of all trochozoan species analyzed (Table 1, Figure 3). Our results reveal that, in general, multiple TLRs are present in highly variable numbers in trochozoan species. The number of TLRs is not reflected by the phylogeny, meaning that species belonging to a same clade do not have a more similar number of TLRs than species belonging to another clade. For instance, the annelids *G. oculata* and *H. robusta* have 39 and 4 TLRs, respectively, while the Phoronid *P. harmeri* has 2. Thus, in this case, the number of TLRs is more similar between an annelid and a phoronid than between two annelids. This is explained by the multiple duplications and losses that have independently occurred in the Toll receptor family during trochozoan evolution.

### P-type/mcc and V-type/scc are not specific for any planulozoan clade

Previous studies suggest that V(ertebrate)-type/scc and P(rotostome)-type/mcc TLRs are restricted to vertebrates and protostomes, respectively [62]. However, our results show that both, P-type/mcc and V-type/scc type TLRs, are present in cnidarians, spiralians, ecdysozoans, and deuterostomes (Table 1; Supplementary Table 2). V-type/scc TLRs are the most abundant TLR type in the spiralian species analyzed. However, many spiralians also have several P-type/mcc TLRs. P-type/mcc TLRs are the predominant TLR type in the ecdysozoan species included in this analysis. For nematodes, tardigrades and onychophorans, which only have one TLR, this TLR was always classified as P-type/mcc. Ecdysozoan species analyzed with more than one TLR have one or more P-type/mcc TLRs and only one V-type/scc. Although the vertebrate TLR complement seems to only contain V-type/scc TLRs [14, 65, 116, 117], P-type/mcc TLRs are also present in other deuterostomes, such as the tunicate *C. intestinalis* [94] and the echinoderm *S. purpuratus* [62] (Table 1, Supplementary Table 2). This suggests that P-type/mcc TLRs were lost in the lineage to the Craniata.

### TLRs form three well-supported clades

Our phylogenetic analysis showed that TLRs group into three clades (Figure 4A), which we named clade α (89 TLRs), clade β (102 TLRs) and clade γ (79 TLRs). Although these three clades are well supported (>60), some of the internal nodes have low support values (<60). The phylogenetic analysis showed that clades β and γ are sister clades and together form the sister group to clade α. All three clades contain both P-type/mcc and V-type/scc TLRs, which makes it difficult to reconstruct whether P-type/mcc or V-type/scc show the ancestral state of TLRs. Furthermore, 2 deuterostome TLRs (from *H. sapiens* and *C. intestinalis*) and 11 spiralian TLRs (2 from species of mollusks and 9 from brachiopods) could not be assigned to any of the above clades. The 9 brachiopod TLRs form a clade with a high support value (>60), but do not group with either the mollusk or the deuterostome sequences. This TLR brachiopod clade is the sister clade to the three main clades (α, β and γ). For these sequences, the alignment showed brachiopod-specific deletions in the amino acid positions 150-220 that are not present in the TLRs belonging to the three main clades (Supplementary Figure 1). To investigate whether this insertion is causing the clustering of the TLRs into three clades, we performed a second phylogenetic analysis (Supplementary Figure 2) with the same parameters of the main analysis (Figure 4A) but excluding the 150-200 amino acid region. The second analysis (Supplementary Figure 2) is able to reconstruct clade α with high support value (>60). However, clade γ is nested within clade β and both of them have low support values (<60). In the second analysis (Supplementary Figure 2), as in the main analysis (Figure 4), the 9 brachiopod sequences cluster together and form the sister clade to the three main clades. However, in the analysis shown in Supplementary Figure 2, the mollusk and deuterostome sequences are included in the clade γ. In the main analysis (Figure 4A), no distinctive motifs were observed in the alignment that justify the exclusion of these sequences from the main clades.

**Figure 4.**
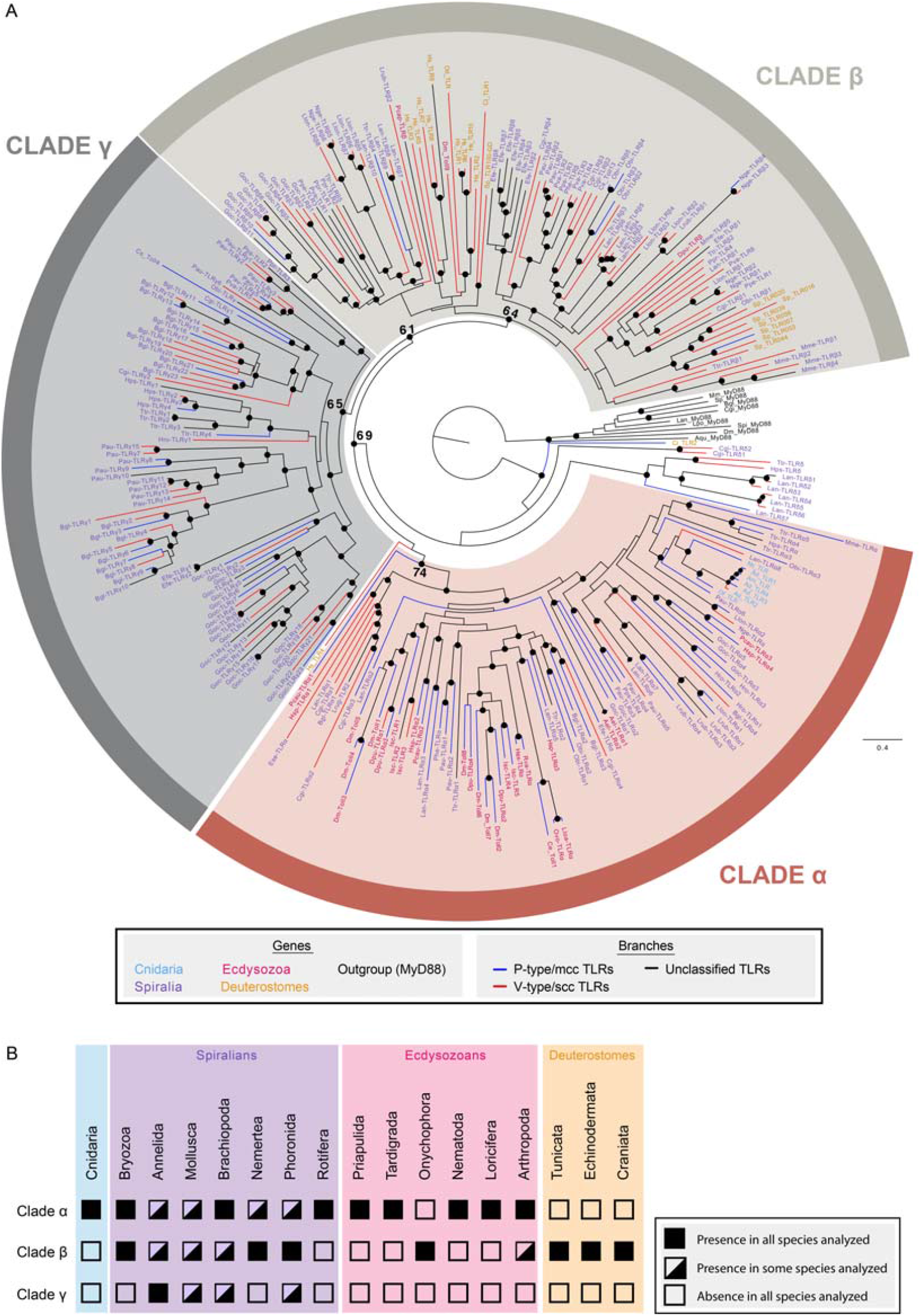
TLR phylogenetic analysis and distribution of P-type/mcc or V-type/scc. A). Phylogenetic analysis of TLRs based on maximum likelihood Bootstrap values are indicated next to the main nodes and all nodes with bootstrap values >60 are marked with full dots. Tip labels contain an abbreviation of the species name and the gene name given in this study (for sequences searched *de novo* here) or in the original study (for sequences obtained from the literature). Numbers in the gene name do not imply gene orthology. Species abbreviations: Ael: *A. elegans*; Ad: *A. digitifera*; Am: *A. millepora*; Bgl: *B. glabrata*; Ce: *C. elegans*; Cgi: *C. gigas*; Ci: *C. intestinalis*; Cs: *C. sinensis*; Dm: *D. melanogaster*; Dpu: *D. pulex*; Efe: *E. fetida*; Ese: *E. senta*; Goc: *G. oculata*; Hex: *H. exemplaris*; Hps: *H. psittacea*; Hro: *H. robusta*; Hsa: *H. sapiens*; Hsp: *H. spinulosus*; Isc: *I. scapularis*; Mme: *M. membranacea*; Nge: *N. geniculatus*; Nv: *N. vectensis*; Lan: *L. anatina*; Lloa: *L. loa*; Llon: *L. longissimus*; Lrub: *L. ruber*; Lrug: *L. rugatus*; Obi: *O. bimaculoides*; Od: *O. dioica*; Of: *O. faveolata*; Ovo: *O. volvulus*; Pau: *P. australis*; Pcap: *P. capensis*; Pcau: *P. caudatus*; Phe: *P. hermeri*; Ppe: *P. peregrina*; Ppr: *P. prolifca*; Pps: *P. psammophila*; Pva: *P*.*vancouverensis*; Rva: *R. varieornatus*; Sp: *S. purpuratus*; Ttr: *T. transversa*. B). Presence/absence in the metazoan groups included in our study.

Clade α includes TLRs from all cnidarian, spiralian and ecdysozoan species analyzed, except for the onychophoran TLR (Figure 4). Because all cnidarian TLRs cluster together, it is likely that only one TLR was present in the last common ancestor of Cnidaria. Clade β is formed by TLRs belonging to deuterostomes, spiralians and three ecdysozoans (two arthropods and the onychophoran TLR) (Figure 4). This suggests that at least the ancestral TLR of Clade β/γ was already present in the last common ancestor of Nephrozoa (Protostomia + Deuterostomia). Furthermore, lineage-specific expansions of clade β TLRs are detected in spiralians and deuterostomes. Clade γ TLRs are present in all trochozoan groups except for the nemertean species analyzed (Figure 4). Clade γ contains TLRs that radiated independently in several lineages. Our alignment shows that 159/181 TLRs belonging to the clades β and γ contain an insertion of 6 amino acids in the positions 349-354 (Supplementary Figure 1). In Clade α, this insertion is only present in Pcau-TLRα1, the sister TLR to all the remaining TLRs belonging to this clade. To exclude that this insertion causes the clustering in three distinct clades, we performed a third phylogenetic analysis (Supplementary Figure 3), in which we applied the same parameters as in the main analysis -shown in Figure 4A-but eliminated the 6 amino acid insertion regions. In the third analysis (Supplementary Figure 3), the three clades could be reconstructed with good support values (>60). However, due to low support values (<60), the relationship between the clades could not be resolved. Moreover, the clustering of the TLRs into the three clades (α, β, γ) was maintained with respect to the main analysis (Supplementary Figure 3, Figure 4A), except for eight phoronid and one human sequences. In the main analysis (Figure 4A), the phoronid sequences cluster together within clade γ, with high support values (>60). This clade of phoronid TLRs is the sister clade to all remaining TLRs in clade γ. Nevertheless, in the third analysis (Supplementary Figure 3), these phoronid TLR sequences constitute a well-supported (>60) clade within clade β, but it is not the sister clade to the remaining TLRs in this clade. In the main analysis (Figure 4A), the human sequence is not included in any of the three main clades, but in the third analysis (Supplementary Figure 3) it does cluster in clade α.

### TLRs are expressed during development in the ecdysozoans *P. caudatus* and *H. exemplaris* and in the spiralians *C. gigas* and *T. transversa*

In order to study the temporal expression of TLRs during ontogeny, we analyzed stage-specific transcriptomes of the priapulid *P. caudatus* [118], the tardigrade *H. exemplaris* [119], the mollusk *C. gigas* [101] and the brachiopod *T. transversa* [120]. All the analyses were performed using both RSEM [121] and kallisto [122] methods.

The expression of the only TLR present in *H. exemplaris* was analyzed in stage-specific transcriptomes of 19 stages (one biological replicate) (Figure 5A; Supplementary Table 3) [119]. Expression of *TLR*α was detected (TMM ≥ 0.15) in time windows during development (zygote, morula, gastrula, elongation, segmentation and differentiation).

**Figure 5.**
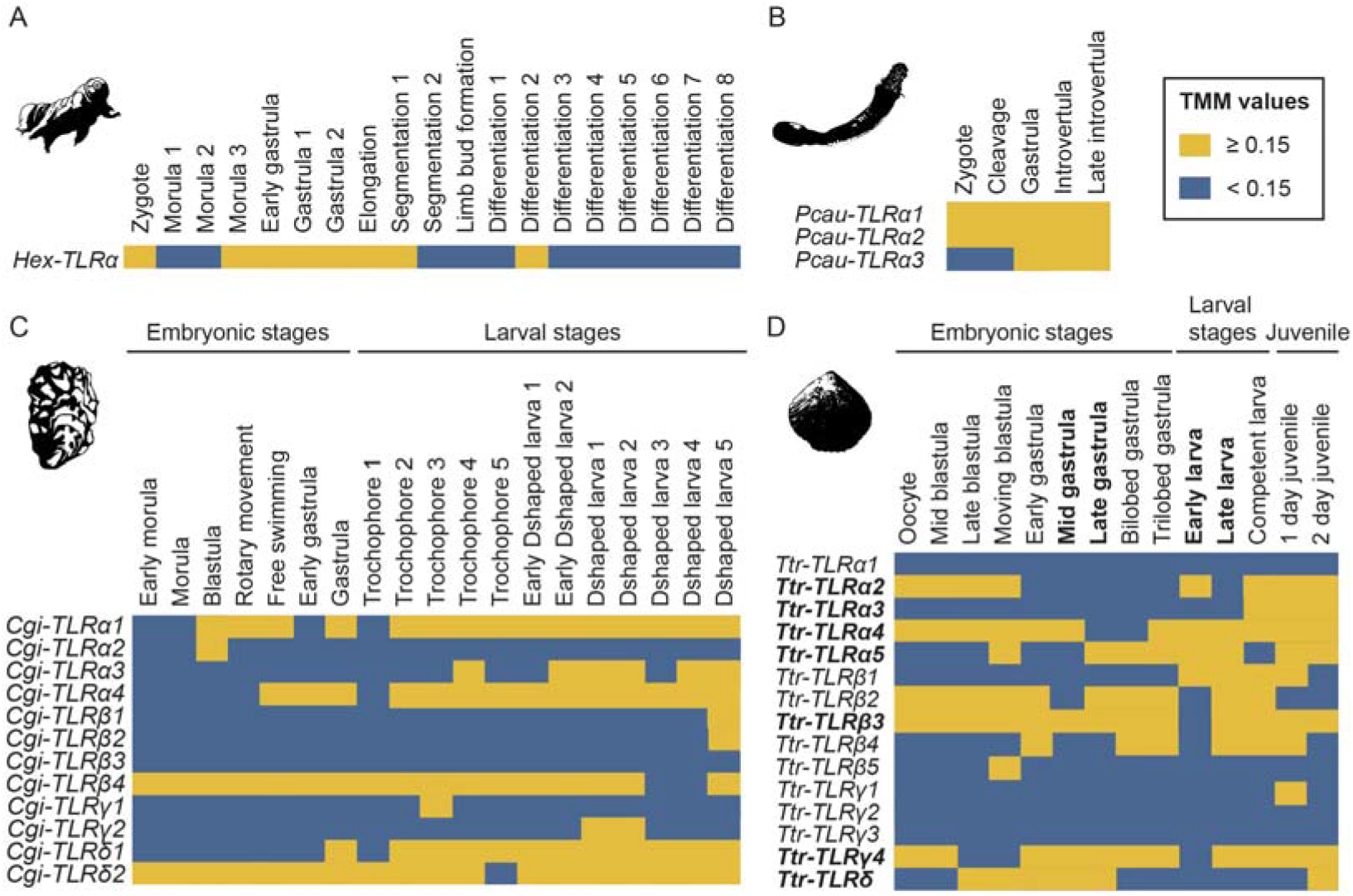
TLR expression in developmental stage-specific transcriptomes of (A) *H. exemplaris*, (B) *P. caudatus*, (C) *C. gigas* and (D) *T. transversa*. Heatmaps corresponding to the average of the RSEM analyses are shown. For heatmaps corresponding to Kallisto analyses see Supplementary Tables 3, 4, 5 and 6. Bold indicates stages and genes for which *in situ* hybridization was performed. TMM: Trimmed means of M values.

Three TLRs were identified in *P. caudatus* transcriptomic survey (Table 1). The expression of these TLRs was analyzed in five embryonic stages (two biological replicates) (Supplementary Table 4) [118]. Our results indicate that all three TLRs found in the transcriptomic survey are expressed during embryonic development (TMM ≥ 0.15). *Pca-TLR*α*1* and *Pca-TLRα2* are expressed in all developmental stages analyzed, whereas *Pca-TLRα3* is expressed only in the later embryonic stages (Figure 5B; Supplementary Table 4).

The expression of the 12 *C. gigas* TLRs (Table 1) was analyzed in stage-specific transcriptomes of 19 stages (one biological replicate) (Supplementary Table 5) [101]. Our results show that at 11 of the 12 TLRs are expressed during development (Figure 5C; Supplementary Table 5). Some TLRs are expressed throughout development (*Cgi-TLRα1, Cgi-TLRα4, Cgi-TLRβ4, Cgi-TLR*δ *1, Cgi-TLR*δ *2*), while others (*Cgi-TLRα2, Cgi-TLRα3, Cgi-TLRβ1, Cgi-TLRβ2, Cgi-TLRγ1, Cgi-TLRγ2*) are only expressed at certain developmental stages. *Cgi-TLRβ3* expression was not detected at any of the stages analyzed.

15 TLRs were found in our transcriptome survey of *T. transversa* (Table 1). Expression of these TLRs was analyzed in stage-specific transcriptomes of 12 developmental stages (with two biological replicates) [120]. Our results suggest that at least 12 of the 15 TLRs are expressed at certain stages during *T. transversa* development (Figure 5D; Supplementary Table 6*). Ttr-TLRα2, Ttr-TLRα5, Ttr-TLRβ1, Ttr-TLRβ4, Ttr-TLRβ5*, and *Ttr-TLR*δ expression is detected in time windows during embryonic and larval stages. All these genes, except *Ttr-TLRβ5*, are expressed in juveniles. For some genes (*Ttr-TLRα4, Ttr-TLRβ2, Ttr-TLRβ3*, and *Ttr-TLRγ4*), expression was detected throughout development. Moreover, expression was not detected at the embryonic and larval stages analyzed for *Ttr-TLRα1, Ttr-TLRγ1, Ttr-TLRγ2* and *Ttr-TLRγ3*. Similarly, *Ttr-TLRα3* expression was only detected in the competent larvae and in the juveniles.

Our analyses show that TLRs are expressed during the development of the spiralians *T. transversa* and *C. gigas* and the ecdysozoans *P. caudatus* and *H. exemplaris*, which suggests that these genes could be involved both in development and immunity during ontogeny. Furthermore, these analyses show that the TLRs expressed during development are not restricted to one TLR clade in the tree shown above, but they are found in all three main clades (e.g. *Ttr-TLRα4, Ttr-TLRβ3, Cgi-TLRγ1*).

Furthermore, in order to validate our stage specific transcriptome results, we performed whole mount *in situ* hybridization (WMISH) for the *T. transversa* mRNAs of *TLRα2, TLRα3, TLRα4, TLRα5, TLRβ3, TLRγ4* and *TLR*δ (Figure 6). Consistently with our stage specific transcriptomic analysis, our WMISH results show that *Ttr-TLRα2* is not expressed at the late gastrula stage (Figure 6A), but the expression is present in the mesoderm and in two pairs of lateral domains in early larvae (Figure 6B). This gene is not expressed in late larvae (Figure 6C). In agreement with our stage specific transcriptomic analysis, we did not detect *Ttr-TLRα3* either in late gastrulae or in the two larval stages analyzed (Figure 6D-F). *Ttr-TLRα4* has a dynamic expression pattern during *T. transversa* development. This gene is expressed in the mesoderm at the early gastrula stage, but, consistent with the stage specific transcriptome analysis, it is not detected in late gastrulae (Figure 6G-H). In early larvae, *Ttr-TLRα4* is expressed in the inner lobe epithelium and in a medial V-shaped mesodermal domain (Figure 6I). In late larvae, this gene is expressed in the brain and in the pedicle (Figure 6J). Consistently with the stage specific transcriptomic analyses, mRNA of *Ttr-TLRα5* is detected in a uniform salt and pepper distribution at the late gastrula stage and the two larval stages for which WMISH was performed (Figure 6K-M). Congruently with the stage specific transcriptomic analyses, *Ttr-TLRβ3* is expressed in the anterior region of the animal in late gastrulae (Figure 6N). However, although *Ttr-TLRβ3* expression was detected in early larvae in the stage specific transcriptome analysis, expression was not detected by WMISH (Figure 6M). Furthermore, *Ttr-TLRβ3* is not expressed in the late larvae (Figure 6P). The expression of *Ttr-TLRγ4* and *Ttr-TLR*δ have a uniformly salt and pepper distribution at the late gastrula and early larvae stages (Figure 6 Q-R and T-U). This salt and pepper transcript distribution is similar in late larvae, although it is absent from the pedicle lobes (Figure 6 S and V). These results conflict with the stage specific transcriptome analyses, as, in this analysis, neither *Ttr-TLRγ4* expression was detected in the early larvae nor *Ttr-TLRδ* in any of the two larval stages tested. Differences between the results of both analyses could be explained by differences and variation of the developmental stages of the specimens used for the stage-specific transcriptome and the WMISH.

**Figure 6.**
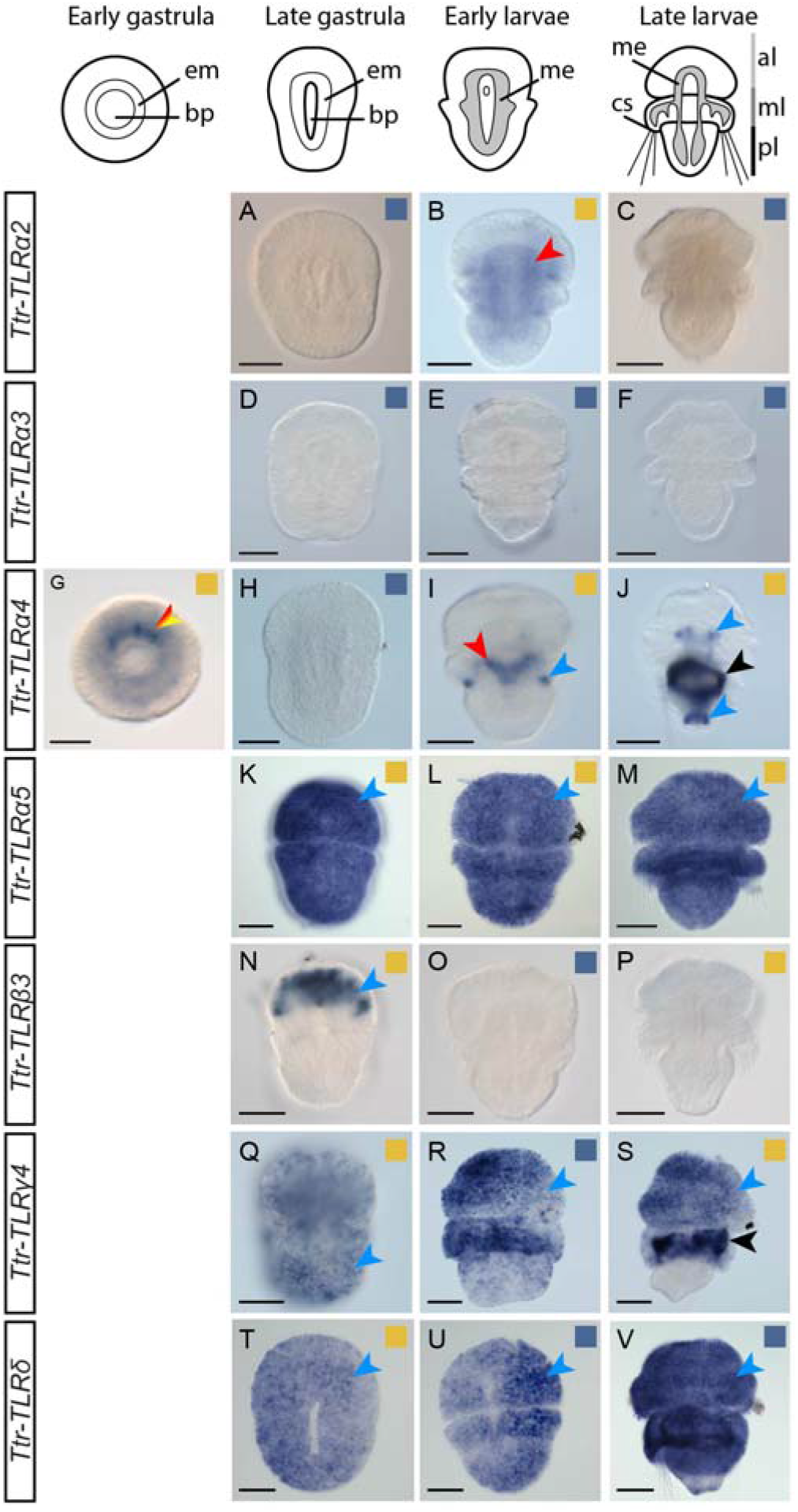
Expression of TLRs during the development of the brachiopod *T. transversa*. Whole-mount *in situ* hybridization (WMISH) of TLRs in *T. transversa* embryos and larvae. Above the WMISH plates, there are schematic representations of each developmental stage analyzed. These representations are not to scale. The name of each gene is indicated in the rectangles on the left. All panels show dorso-ventral views and anterior to the top. Squares in the top-right of each plate indicate whether the expression was detected (yellow) or not (blue) in the stage-specific transcriptome analysis. Ectoderm, mesoderm and endoderm is indicated with blue, red and yellow arrowheads, respectively. The red and yellow arrowhead indicates endomesoderm. The ring-shape staining present in the late larvae *Ttr-TLRα4* and *Ttr-TLRγ4* is background staining (black arrowhead) [123]. Scale bar indicates 50 μm. al: apical lobe; bp: blastopore; cs: chaetal sacs; em: endomesoderm; me: mesoderm; ml: mantle lobe; pl: pedicle lobe.

## Discussion

### The evolution of the TLR family is characterized by losses, expansion and conservation

As shown in previous studies, TLRs are absent in the Platyhelminthes *S. mediterranea* and *S. mansoni* [89]. Here, we show that this receptor family is also absent from the genomes of three other platyhelminth species (*M. lignano, E. multilocularis* and *H. microstoma*). Thus, TLRs are absent in species belonging to four different platyhelminth lineages (Macrostomorpha – *M. lignano*; Cestoda – *E. multiocularis* and *H. microstoma*; Tricladida – *S. mediterranea*; and Digenea *– S. mansoni*) suggesting that TLRs could have been lost during early platyhelminth evolution. This hypothesis is reinforced by the lack of TLRs in *M. lignano*, member of Macrostomorpha, an early-diverging platyhelminth lineage [104]. In rotifers, even though TLRs could not be detected in *A. vaga* [67], *E. gadi, R. tardigrada* and *M. hirudinaceus*, our transcriptome survey revealed one TLR in the monogonont rotifer *E. senta*. This suggests that TLRs would have been independently lost in some rotifer lineages. So far, we did not detect TLRs in the genomes and transcriptomes of the species belonging to Xenacoelomorpha, Cycliophora, Micrognathozoa, and Gastrotricha, suggesting that TLRs were lost in these lineages. How the immune response is achieved in animals that lack TLRs is unknown, but it could be triggered by other components of the Toll pathway e.g. TLR-like molecules [14, 67–69], similar to what has been shown for LRR-only TLR-like and TIR-only TLR-like in *Hydra* [72, 73].

Another outcome of this study is the remarkable expansion that the TLRs family exhibits in trochozoans. Evolution of this gene family in trochozoans is characterized by multiple duplications and losses, having as a consequence a very variable number of the TLRs complement in trochozoans. Moreover, in our phylogenetic analysis, TLRs of the same species and clades mostly group together, indicating the existence of multiple independent duplications (Figure 4A). The same has been shown also in previous phylogenetic analyses of TLRs (Figure 6) [13, 63, 86].

In contrast to trochozoans, our results show that the number of TLR in ecdysozoans has been relatively conserved during evolution. At least, few TLR gene duplications have occurred in this lineage, including recent independent duplications in arthropods, priapulids or loriciferans.

### The evolution of the three clades (α, β, γ) of TLRs

There are very few studies assessing the phylogenetic relationships of TLRs within the main metazoan clades (Figure 7) [63, 86]. The study of Davidson et al., 2008 [63] recovered three clades of TLRs. However, the relationships between the clades remain unclear. Furthermore, the composition of the clades slightly differs in both analyses (e.g. while our study shows that deuterostome TLRs belong to one clade – clade β – their results suggest that deuterostome TLRs are present in two clades – clades A and B) [63]. However, their phylogenetic study is limited by the number of sequences and species included. Similar to Luo and Zheng, 2000 [124]; and Luna et al., 2002 [125], our results suggest that ecdysozoan and deuterostome TLRs evolved independently from a common TLR precursor.

**Figure 7.**
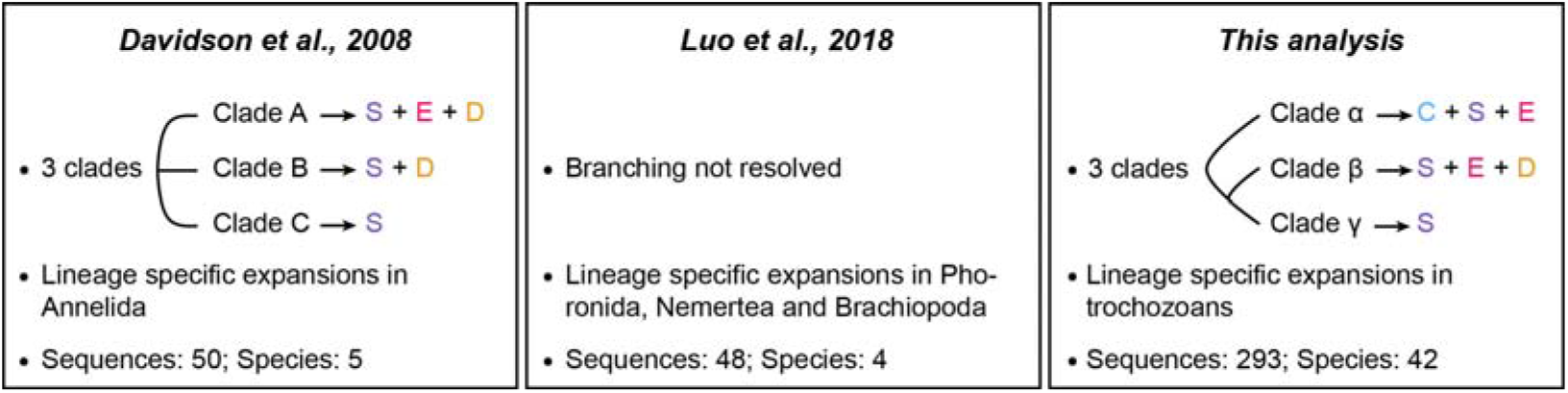
Comparison between Davidson et al., 2008; Luo et al., 2018; and this study. The main conclusions and the number of TLRs and species included in the three studies are compared. Cnidaria (C), Spiralia (S), Edysozoa (E) and Deuterostomia (D).

Previous studies suggest that TLRs originated likely by the fusion of an LRR-only and a TIR-only TLR genes in the lineage to Planulozoa (Cnidaria + Bilateria) [7, 14, 65]. Here, we hypothesize that the planulozoan stem species had only one TLR (Figure 8), the *proto*-TLR. This is supported by the fact that all cnidarian TLRs included in our analysis cluster in a monophyletic group within clade α, which is consistent with the results of Brennan and Gilmore, 2018 [13]. During cnidarian evolution, this gene was lost in some lineages, e.g. *Hydra* [72], *Clytia* [81], and multiplied in others, e.g. *A. digitifera* [69].

**Figure 8.**
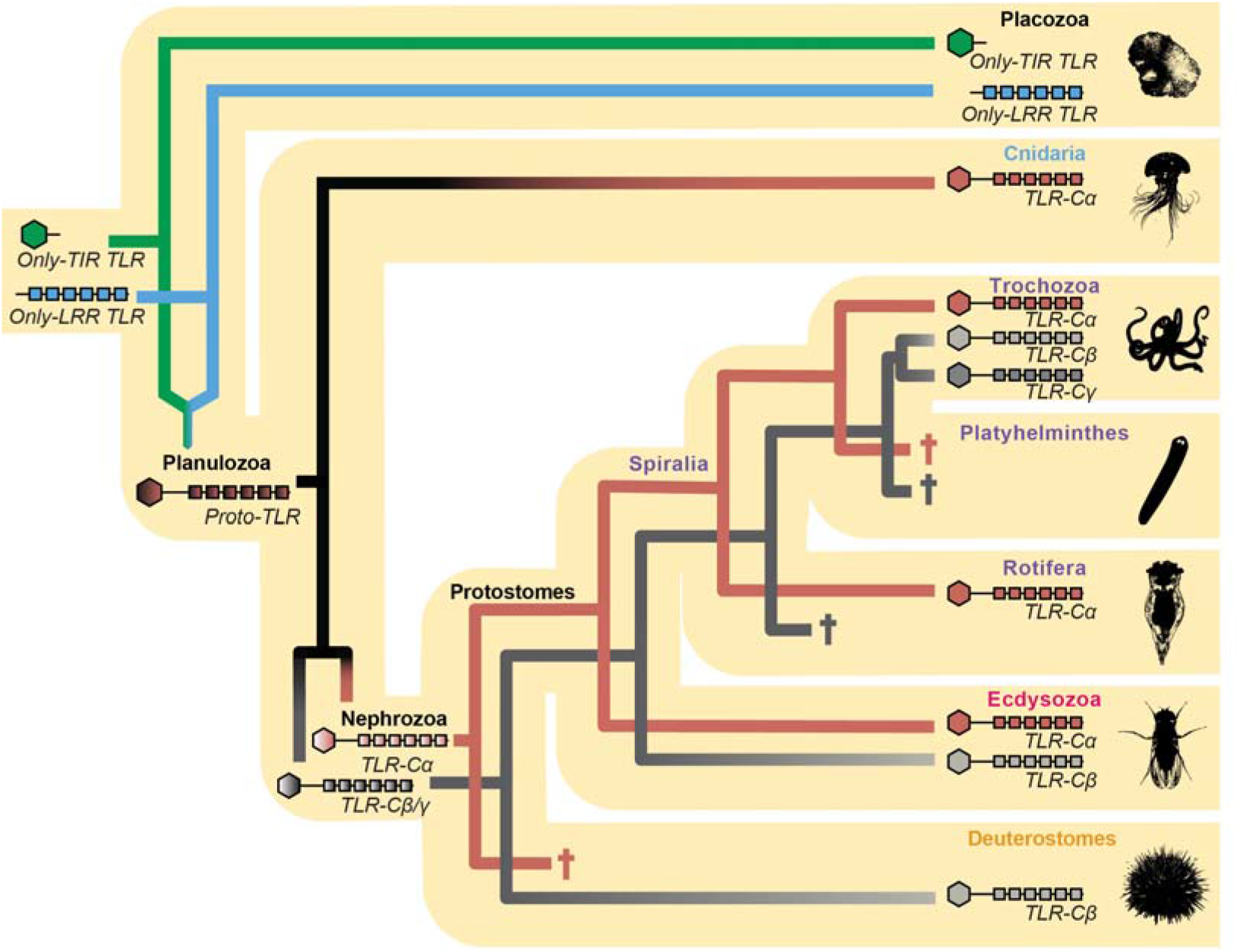
Origin and evolution of TLRs. Gene lineages are depicted in different colors (*Only-TIR TLR*: green; *only-LRR TLRs*: blue; *TLR-Cα*: brown; *TLR-Cβ/γ*: dark grey; TLR clade *β*: light grey; and TLR clade *γ*: mid-grey) within the metazoan tree. Gene losses are indicated with a cross. The question mark shows uncertainty in that specific statement. Phylogeny according to: [95]

After the split into the cnidarian and bilaterian lineages, the *proto*-TLR was duplicated in the lineage to the Bilateria, giving rise to a clade *α* type TLR gene (*TLR-Cα*) and the *proto*-TLR gene of clades *β* and *γ* (*TLR-Cβ/γ*) (Figure 8). However, our results indicate that *TLR-Cα* was lost during early deuterostome evolution. Later, expansions of *TLR-Cβ/γ* generated the TLR diversity found in deuterostomes. Furthermore, as vertebrate TLRs diversified within the vertebrate lineage, it is impossible to make one-to-one orthology gene assignments between the vertebrate TLRs and the invertebrate TLRs [65].

The protostome stem species and the spiralian stem species had likely two TLRs: *TLR-Cα* and *TLR-Cβ/γ* (Figure 8). During early trochozoan evolution, the spiralian *TLR-Cβ/γ* gene was duplicated, giving raise to the ancestral TLR from clade *β* in trochozoans (*TLR-Cβ*) and the ancestral TLRs from clade *γ* (*TLR-Cγ*). This is supported by the fact that clade *β* and clade *γ* are sister clades and clade *γ* is only present in trochozoans. Later, episodes of gene duplication generated the larger diversity of TLRs from clade *β* and clade *γ* in trochozoans. These expansions could have occurred due to the necessity to adapt to microbe rich environments [126, 127]. Losses of both TLRs seem to have occurred in non-trochozoan lineages, e.g. in platyhelminths and rotifers.

Our results show that the ecdysozoan stem species had two TLRs (Figure 8) belonging to clade *α* and clade *β*/*γ*. Although, in general, the number of TLRs is low, few duplications of *TLR-Cα* occurred in some lineages (e.g. arthropods, priapulids, loriciferans). Furthermore, our analysis shows that the surveyed priapulids, tardigrades, nematodes and loriciferan lack TLRs from clade *β*; whereas clade *β* TLRs are present in the majority of the arthropods and in the onychophoran surveyed. This would imply that TLR clade *β* would have been lost independently in the early branching ecdysozoans but not in the most late-branching lineages [95, 128].

### Are protostome TLRs involved in immunity and development during ontogeny?

TLRs are well known to play a key role in adult innate immunity in planulozoans [11, 22–26]. During ontogeny, this gene family has also been shown to be involved in a great number of developmental processes both in arthropods and vertebrates [2, 8, 9, 36–38, 41–44]. Here, we identify TLRs expressed during ontogeny in four protostome species (the ecdysozoans *H. exemplaris* and *P. caudatus* and the spiralians *C. gigas* and *T. transversa*) (Figures 5 and 6). Expression of TLRs was observed for some TLRs in short developmental time windows (the *H. exemplaris Hex-TLRα*; the *C. gigas Cgi-TLRα2, Cgi-TLRα3, Cgi-TLRβ1, Cgi-TLRβ2, Cgi-TLRγ1, Cgi-TLRγ2*; and the *T. transversa Ttr-TLRα2, Ttr-TLRα5, Ttr-TLRβ1, Ttr-TLRβ4, Ttr-TLRβ5*), suggesting a possible role of these genes in development, as genes involved in developmental processes are usually expressed for defined periods of time in tissues in order to participate in specific developmental processes [129–131]. For instance, expression during early embryonic stages of the *T. transversa Ttr-TLRα2* (Figure 5) might suggest its involvement in dorso-ventral axis specification, as it has been shown for the *Drosophila Toll* [8, 9]. Later, in the early larvae, transcription of this gene is transiently activated in the mesoderm (Figures 5 and 6), suggesting that this gene might be also involved in mesoderm development. However, our analyses do not exclude the possibility that these genes might also be involved in immunity, as these TLRs could have a dual role, as it has been shown for the *Drosophila Toll* [10] and the only TLR in the cnidarian *N. vectensis* [27]. Discerning the role of TLRs expressed in broad time windows or during the whole development (the three *P. caudatus* TLRs; the *C. gigas Cgi-TLRα1, Cgi-TLRα4, Cgi-TLRβ4, Cgi-TLRδ 1, Cgi-TLRδ 2*; and the *T. transversa Ttr-TLRα4, Ttr-TLRβ2, Ttr-TLRβ3*, and *Ttr-TLRγ4*) is complex, as these genes could be involved either in immunity or in development, or both. However, detection of immune processes in our analyses is not possible with the data available. Therefore, further investigations are required to gain more knowledge on functions of TLRs during development. Immune roles of the TLRs during ontogeny should not be underestimated: Many marine invertebrate embryos and larvae live in environments rich in microbial pathogens [132, 133]. Pathogens cause mortality of embryos and larvae but also provoke anomalies during development [134, 135]. Therefore, these embryos and larvae need immune defenses to fight pathogens [133]. Actually, few studies have shown that the Toll pathway is involved in immunity during ontogeny in arthropods, mollusks and amphioxus [18, 135–137], and other immune-related genes have also been found to be involved in immunity during mollusk and echinoderm development [136, 138–140]. Additionally, in planulozoans it has been shown that TLRs are involved in adult immunity [11, 22–26]. Thus, TLRs are probably also involved in immunity during ontogeny across the metazoan tree.

## Conclusions

Based on our data we propose a scenario in which TLRs evolved from an ancestral *proto*-TLR that originated before the split into the cnidarian and the bilaterian lineage. Duplications and losses characterize the evolution of TLRs in the main metazoan groups. The *proto*-TLR duplicated in different metazoan lineages and gave rise to three TLR clades. This TLR complement was expanded during Trochozoa evolution, while it was lost in some non-trochozoan spiralian lineages (e.g. platyhelminths, cycliophorans, micrognathozoans, gastrotrichs and some rotifers). Ecdysozoans possess a low number of Clade *α* and Clade *β* TLRs; whereas all deuterostome TLRs belong to clade *β*, being originated by radiations in the different lineages. Furthermore, our data shows that TLRs are expressed during ontogeny in two ecdysozoan and two spiralian species, suggesting that these genes could be involved in immunity and development.

## Materials and methods

### Genomic and transcriptomic surveys

We surveyed TLRs 20 genomes and 25 transcriptomes (Supplementary Table 1). Overall, only high-quality transcriptomes (BUSCO values >70% - Supplementary Table 1) were selected, but lower quality transcriptomes were also included when they represented a species from a low investigated clade (e.g. the *loriciferan A. elegans* transcriptome (BUSCO value 36.2%)). In order to search for the TLR sequences, hmmer profiles for the TIR and the LRR domains were generated using HMMER software version 3.2.1 (www.hmmer.org). The hmmer profile for the TIR domain was blasted against each genome/transcriptome in order to obtain a database of proteins containing the TIR domain. Next, the LRR hmmer profile was blasted to the TIR domain-containing sequences database. These sequences were validated by BLAST [141] (www.blast.ncbi.nlm.nih.gov) and SMART [142, 143] (http://smart.embl.de/). Sequences from the same species with >90% similarity were considered to be polymorphisms and only one of them was considered for the analyses.

### Phylogenetic analysis

TLR sequences obtained from the genome/transcriptome surveys and NCBI (www.ncbi.nlm.nih.gov) (Supplementary Table 2) were aligned using MAFFT software version 7 [144], applying the L-INS-I algorithm. The MyD88 protein, an adaptor of the Toll pathway, was selected as an output. The TIR domain of well annotated MyD88 proteins (Supplementary Table 2) were included in the alignment. The alignment was trimmed manually in order to obtain a fragment containing one LRR domain, the transmembrane domain, and the TIR domain. This was followed by a second trimming step performed with TrimAl software version 1.2 [145] using the gappyout trimming model. The final alignment used to perform the phylogenetic analysis contains 375 amino acids. The maximum likelihood phylogenetic analysis was performed using IQ-TREE software [146] in the CIPRES Science Gateway V.3.3 [147] (http://www.phylo.org). LG+R8 was selected as the best-fit model (according to BIC) and was applied for the phylogenetic reconstruction. Bootstrap values were calculated running 1000 replicates using ultrafast bootstrap.

### TLR classification

TLR sequences from the genomic/transcriptomic surveys, as well as the ones obtained from the literature and NCBI database, were classified into P-type/mcc and V-type. In order to do so, the number of LRR domains was analyzed with LRRfinder software [148] (http://www.lrrfinder.com). Next, sequences were classified applying the same criteria followed by Brennan and Gilmore, 2018 [13]. Some TLR sequences were incomplete and they could not be classified into P-type/mcc or V-type.

### Stage specific transcriptome analyses

In order to assess the expression of TLR genes, we examined publicly available stage-transcriptomic data of various developmental stages for the spiralians *C. gigas* and *T. transversa* and the ecdysozoans *P. caudatus* and *H. dujardini*. For *C. gigas*, we examined 19 developmental time-points from early morula to D-shaped larvae, being the transcriptomic data previously published in [101] (accession number: SRR334225-SRR334243). For *T. transversa*, 14 stages from oocyte to 2-day juvenile were analyzed, being this dataset available from [120]. For *P. caudatus*, only 5 embryonic stages (from zigot to late introvertula) were analyzed. The transcriptomic data was obtained from [118]. The 20 *H. exemplaris* embryonic transcriptomes analyzed (from zigot to differentiation) were obtained from [119] (accession numbers: SRR1755597, SRR1755601, SRR1755603, SRR1755606, SRR1755610, SRR1755612, SRR1755621, SRR1755623, SRR1755627, SRR1755631, SRR1755637, SRR1755644, SRR1755647, SRR1755650, SRR1755656, SRR1755662, SRR1755666, SRR1755706, SRR1755715, SRR1755719). We first performed quality-trimming on downloaded RNA-seq raw reads using Trimmomatic v.0.38 [149], removing low quality or N bases (parameter settings: LEADING:20 TRAILING:20 SLIDINGWINDOW:4:20). To estimate the transcript abundancies, quality-trimmed reads were aligned to reference transcriptome assemblies (*C. gigas* [101], *T. transversa* and *P. caudatus* [97], *H. exemplaris* [109]). We applied two quantification methods: an alignment-based method using Bowtie2 [150] and RSEM [121], and the ultra-fast alignment-free method kallisto [122]. Both methods reported normalized expression values in transcripts per million (TPM), and we further executed cross-sample normalization among different developmental-stage samples by TMM method [151]. To define a criterion for gene expression value in this study, we performed *in situ* hybridization of selected TLR genes at different developmental stages in *Terebratalia*, as well as examining expression values in our analysis corresponding to *in situ* hybridization data of *Hox* genes in *Terebratalia* [120] and *Wnt* genes in *Priapulus* [118]. We considered expression for values ≥ 0.15.

### Animal collection and embryonic cultures

Adult *T. transversa* specimens were collected in Friday Harbor, USA. The eggs were fertilized, and animals were fixed at different developmental stages with 4% paraformaldehyde for 1h at room temperature, as described elsewhere [120, 152]. Next, the samples were repeatedly washed in Ptw and stored in 100% methanol.

### Gene cloning, probe synthesis, in situ hybridization and imaging

Specific primers for *T. transversa* TLRs were designed using the MacVector 10.6.0 software. TLRs were amplified and inserted into pGEM-T Easy vectors (Promega, USA) and transformed in competent *E. coli* cells. Minipreps were prepared using NucleoSpin®Plasmid kit (Macherey-Nagel) and sequenced in the Sequencing facility of the University of Bergen. RNA probes were transcribed using digoxigenin-11-UTP (Roche, USA) with the MEGAscript^™^ kit (Invitrogen, Thermo Fisher). Whole mount *in situ* hybridization (WMISH) was performed as described in [120, 153]. Probes were hybridized at a concentration of 1 ng/μl at 67°C during 72h. Next, they were detected with anti-digoxigenin-AP antibody [1:5000] (Roche) and developed using NBT/BCIP (Roche). Samples were washed twice in 100% ethanol and re-hydrated in descending ethanol steps (75%, 50% and 25% ethanol in PBS). Samples were mounted in 70% glycerol. Samples were imaged using Axiocam HRc camera connected to an Axioscope Ax10 (Zeiss, Oberkochen, Germany). Images were analyzed using Fiji and Adobe Photoshop CS6.

### Illustrations

Figure plates and illustrations were made with Adobe Illustrator CS6.

## List of abbreviations

AMPs: Antimicrobial peptides
BCIP: 5-bromo-4-chloro-3-indolyl phosphate
BUSCO: Benchmarking Universal Single-Copy Orthologs
IL-1: Interleukin-I receptor
LRR: Leucine-rich repeat
LRRCT: Leucine-rich repeat C-terminal domain
LRRNT: Leucine-rich repeat N-terminal domain
Mcc: multiple cysteine cluster
NBT: nitro blue tetrazolium
NOD: Nucleotide oligomerization domain
PBS: Phosphate-Buffered Saline
P-type: Protostome type
PTw: PBS with 0.1% Tween® 20
RSEM: RNA-Seq by Expectation Maximization
Scc: Single cysteine cluster
TIR: Toll/IL-1 receptor
TLR: Toll-like receptor
*TLR-Cα*: clade *α* type TLR gene
*TLR-Cβ*: clade *β* type TLR gene
*TLR-Cγ*: clade *γ* type TLR gene
*TLR-Cβ/γ*: proto-TLR gene of clades *β* and *γ*
TLR-like: Toll-like receptor-like
TM: Transmembrane
TMM: Trimmed mean of M values
TPM: Transcripts per million
V-type: Vertebrate type
WMISH: Whole mount in situ hybridization

## Declarations

### Ethics approval and consent to participate

Not applicable.

### Consent for publication

Not applicable.

### Availability of data and materials

The datasets supporting the conclusions of this article are included within the article and its additional files. TLR and MyD88 sequences obtained in the genomic/transcriptomic surveys and used in the phylogenetic analysis, together with their NCBI accession numbers, are available in the Supplementary Table 2.

### Author’s contributions

AH designed the study. AOA performed the genome and transcriptome surveys, the phylogenetic analyses, the stage-specific data interpretation, the *in situ* hybridization, and wrote a draft manuscript. TML performed the transcriptome stage-specific analysis. AH, TML and AOA discussed the data and revised and contributed to the writing. All authors read and approved the manuscript.

## Acknowledgements

We want to thank Daniel Thiel for instructing AOA in performing genome/transcriptome surveys and phylogenetic analysis and for discussions. We also thank Ferenc Kagan for providing the BUSCO values for the transcriptomes; Ludwik Gasiorowski for discussions; Carmen Andrikou for reading the manuscript and discussions; and Timothy Lynagh for critically reading the manuscript. Furthermore, we would like to thank other former and present members from the Hejnol lab for collecting and fixing the T. transversa specimens and Nadezhda Rimskaya-Korsakova for collecting and providing Galathowenia.

