## Supplementary Figure 1 for "The evolution of the metazoan Toll receptor family and its expression during protostome development"

**Supplementary Figure 1. Phylogenetic analysis alignment.** Regions rich in gaps located in the positions 150-220 for TLRs not belonging to the three main clades are marked in magenta. In cyan, we mark the positions 349-354 characteristic from clades β and γ; and the gaps corresponding for these positions for the TLRs belonging to clade α.

**>Mme-TLRβ3**
NYQTIKNMVYLDLSDNKISIIP----NDIFYQMVKMSHLNLIGNQIVSLDNNMFLYNANL
QSLYLSNNRLTTFDSKLLNHCKALERLMISQNKISLFDVEFVNEIDSLRSVRIDENPFDC
SCGQLFFQSWVQRSKK----KFGQDLQCVQPSNLLNQKISEY---QPQQCV-KWVVDGVA
----LVGLVLIIIVTYQLRWYLTNIPYTRSMRY------RQQDECHYDAMVIFSNKDTDW
V-KELMRELEEDRENKIYIRVRDDISGTITFEECSKVMNQSRKIILILSNSFLAETECIS
EVEFAGNELFSTANGRVMILVLEELQESRIDL-VRSLMIEANVIELLESTRKMREVWQKL
KKFVNHQKP
**>Mme-TLRβ4**
NFKTIKHLTYLDLSDNSLPSIP----NDLFDNMPKLKTLKMISNLISSLDRRQFIHNSQL
RTLDMTKNLLETFHVSVFANDTLIKALSLRTNKISMFDETFTQFVSTLTYLRIENNSFDC
SCGQLYFQKWANSSKK----QYGDKLICHSPGQLRNQRIVIY---QLFDCY----WVLVG
LA--LVGIITTVLLMYRFRWYLAHLRFALSVAE-RLVDIKQQDQCKYDAMVLFSDEETNW
V-KRLLIELEEDRENLIYIRARDDITGDVNFDSCCKIMKQSRKIILVLSNSFLREDECIS
EATFAGSELFSTAKERILILVLEELQEPL-NP-VSSLLVETHYIDLYETTRKMRAIWQKL
RRFVSHRAN
**>Mme-TLRα**
-LTAFPNIKI------------------------AMEELSLSHNTITELPEDPFYWLRHV
VHLNIQHNELTPQNIPIFDRLGKISYLFLAYNNIVYFPPSIR---KNFKALSITGNKITC
-CNGHWMKTWLQEQNE--TIWNSLTAHCKD----QAHPIIQLDPDGFS-CP-LYLAPIII
SIALTIATMMTSVAVYVYSFELKIILFKLNLHPR---SV-DSESLDYDLYLMYNYADSPW
ATEKLLPGLE-
KFGYRVYVPERDMGIGEITAEARANAFASTHRVLVVVSQKFIDSGESMK
EFFHAHEHENSTTRRYLVLVKLEKI--NRTD-IFKKYMSTNFFVSVKS------KFWYNL
RYWLPREST
**>Mme-TLRβ5**
LFHGMKNLKTLLLMHNHLGLSFSDDYTGFLSRLPRLMVVNMSSNGITVLPKQMISNTS
AL
EVLDLGMNHIYSWDSQTFQGAVGLKKLLLNTNRLALFNETSFSDLKNLTVLDLGNNPF
AC
TCDMRWFRDWLKTT--KVHVNNSHTYTCTSPAKMQGTHLIEFELTTLQ-
CVPVWIVSGCF
IS--LLLILIMVCVLYRYRWRIRFALYKCSKSCKAYQRLPQTDRPLYAAFFSFCSEDENI
IEEQILPNIDNDAGVYPLIHRIKYDPSRTYLDCLEKALLTSPATVVMLCQHYKEDRQCEL
ELAAS----LQEEDRRIILVVLDDIVQRKLPVALRVMLNRNEAIEWHRNEQQARMLKGKL
AEALED---
**>Mme-TLRβ2**
NFKAISNLEYLDLDTNNLTFIR----NHTFDHMPNLNTLILSANNLKHIDDQAFIHNYNL
ATLLLQANKFSVFNVTLLEGPKNLRKLCISTNLITHFDSSFVKFMGTLQTVKIANNPFDC
SCGRKFFSDWLNRTKL----HESVGLECTTPENMAQKKVYNY---EDLECT-PLIWSAVV
LC-LIIVTIMIAVPCYRYRWYISHMVIMQAVKD-RAMDIKHSDECKYDAMILSSEADMKF
V-KTLLLHLEEDRSNRLYHSLRDAIPGTYRFESLCEVMRQSRKIIIVISNSYLNSSECMS
EAAFAGEELFGTKKEKIVVLVLEDLNEDLMPS-IAGLLTET-VIDLPESKKTMAPVWDRL

KKFVENKP-
**>Mme-TLRβ1**
FYPQLKQLQKLDISGNSFKYLD----PNAFNEMKYLSSVIAQSNPLSILPDTIFISNMKL
VKVDFSNCFLDELDNTIVKSLPKLEHLYLKYNQFTSFAPSVISVVKALKTLSLLGNPFDC
DCNIGRLQDWLSETEK----IDVINITCGGPEHAADTSIFEY---PPPRCK----LPLII
GC-AVVGFVLLILLCICSRWYISHRKILPELKG-ILKNIRYGYKCDYDAVVCYSDIDQQW
VGGRLVPALESKKA-RLYIYERDSTIGAEKTVQIRDAMERSRNVIIVLSKSYLASEAFLP
EVDIVADVMRQNEKGRILLLALDDLDNRKMDP-IKLLTLTEKTLNVV------------
---------
**>Goc-TLRγ23**
SLSCWPRMRKLLLGNIELQQIFDHGNETIFNNCTYMKTIDLQNTGIKRLPQNTFLDMEN
V
QYINISNNKLTSLDI--LTSTKNL-TLNISSNLLDHLSNKMTQLLDQMEILDLSYNPIMC
GCPQIDFIVWLKNT--YVEIYNLNQYQCVYQDGTK--YINDISLSQLKQCN-IIIQAVCS
IGS-VVVIGIISIIIYSYRHKLEYLLLIARHAAKSKKDKHNDKTFNFHGFVSYSSEDDLW
IVEQLHMKMEQDFGLKLCIHERDFLPGYFITENISSFMEASRKTVIVLSNNYLKSKWCTF
EFELAKCKLIEATFNTMVMILLHDLDQRKVSPALHKYLKQKTYLKWPKDSSQQPAFWL
RL
KEALDQKPE
**>Goc-TLRγ17**
VLNCFPSLTEVRIGGNQL----NT-HIMMFANCTKLTHLDVSHNKLVSMPKDAFHETPNL
IHLNLSGNLFANLEIHEVTQLTKLQVLDLSYNRFQAIPESWRHTIQLLGKLYISGNPFMC
SCDTVDHLIWLQSI--QDLLDDPTHLTCRDTNGKEY-TIMQIHIGRFKECIKSMVQAGCI
PSAIVLVIVGISLYIYKRRFRFQYLALVARANIN--LHAPQIPDYTYDAFISYSSLDIEY
ML-TLYQKLEQEHNYELCIDMRNFRPGNPIDDEITNGIMDSHKIILVISQNFLRSGWCLY
EMQLAHGELAVRGGDGLLLILKEPRPQELITDKLQGLLDSRIYLEWSEEGDRQQVFWQ
RL
RDALGMPLQ
**>Goc-TLRβ11**
VLSNSTMIESVDLSGNLLFRYTDEFLCDLLKNLVNLEEISLFDNYLTHVPSCLFRASSRI
IGIYLSRNRIAYIQKGVFDSLYQLEELDLDDNSITFIDPSNFYNTPSLSWLTIENNRFSC
DCRLTGFRDWTAEH--QDIIEGP----CETPKQLKGEAVHAYTTTWLE-CNTVFIICGSL
----LFFLLVVTGLLFYFWKDIKYIKMVHRAKGKGYIPLNDNNQVLYDAFISYHPEKKFW
VEVDLIPTLEDDVQFNIMYDER-FDTG-SIFTLTEENIAQSRKILFVVSRGWIQAGWNQF
ELDMAMIKLIDDHRDMIIVLLMEHIPKKEMPDKLKMMVKYNKCLKWSDNEHKQRIFRRD
L
KLELGKE--
**>Goc-TLRγ16**
VLNCFPSLTEVRIGGNQL----NT-HIMMFANCTKLTHLDVSHNKLVSMPKDAFHETPNL
IHLNLSGNLFANLEIHEVTQLTKLQVLDLSYNRFHVIPESWRHTEKILGKLYISGNPFMC
SCDTVDHLIWLQSI--QDLLDDPTHLTCRDTNGKEY-TIMQIHIGRFKECIKSMVQAGCI
PSAIVLVIVGISLYIYKRRFRFQYLALVARANIN--LHAPQIPDYTYDAFISYSSLDIEY
ML-TLYQKLEQEHNYELCIDMRNFRPGNPIDDEITNGIMDSHKIILVISQNFLRSGWCLY
EMQLAHGELAVRGGEGLLLILKEPRPPELITDKLQGLLDSRIYLEWSEDGDKQQVFWQ
KL
RDALGMPLQ

**>Goc-TLRα2**
-LHNLATYP-LNVSYNNLNKIEHCLIPYLFDSLYALESLDLSFNLLTSVPSQLFSSLISL
SALYLDHNDIRFLPQGMLFNSTHVGKLTLHQNKIETLQINIFQSLQFLTTITLADNPWVC
NCSMFDFCKWLHSNWT--KVEDKSSLTCKNGSN-----LIQFT--YNCTCKQDNRIVLGI
CGSLITLLSIGLGLVYYYHDNLRFFLYLFGWRF----PVRNNGEAYFDIFICYSSKDNKY
VITKLLRYLETKPPYKVCIHERDFIPGDYIIDNIVRCINKSKTIILVLSNNFVNSMWCLG
EFQMAYHNAFENRHNNIIPILLGDLNLDHLDPTLRTFVGMNNYLRKDE------LFLQRL
LVALPEPSN
**>Goc-TLRγ10**
ALTCMESLEKLNIANNNF----NETDINIFENCTKLHILNLSYNELENIPKDTFNETTNL
ANLDLSGNKLSNIAF--LENQRNLTFLNLAGNSIQYISPPLTQDISRLMKINLDDNFLRC
GCEDIVFIDWLKNN--EDQIINWEKLKCIDDAGLY--NIQTINTEWTQQCNMNIIIAMSI
MTGFILFTTIIACCLYRHRYKVHYLYLLFRSWFH---KPDDANQYNFDGFISYSSLDKTW
ALETMYANLATKYGYNICVDERNFRPGQHLVDIIIETINTSNKIMLVITQNFLRSGWCLY
EMKMARGELATRGRDCLILILKDPIPQELITPTLRQLLESRIYLEWSEDRDRKALFWRKL
CDALGEPRH
**>Goc-TLRγ9**
ALTCMESLEKLNIANNNF----NETEINIFENCSKLHTLDLSYNELENIPKDTFNETKKL
VNLNLSGNKLSNIAF--LENQRNLTFLNLAGNSIQYISPLLTQDISRLMKINLDDNFLRC
GCEDIVFIDWLKNN--EDQIINWEKLKCIDDAGLY--NIQTINTEWTRQCNMNIIIAMSI
MTGFILFTTIIACCLYRHRYKVHYLYLLFRSWFH---KPDDANQYNFDGFISYSSLDKTW
ALETMYANLATKYGYNICVDERNFRPGQHLVDIIIETINTSNKIMLVISQNFLRSGWCLY
EMKMARGELATRGRDCLILILKDPIPQELITPTLRQLLESRIYLEWSEDRDRKALFWRKL
CDALGEPRH
**>Goc-TLRγ3**
SLDYFPSLKILLLGSNGLGPLIHDNDGRLFANLSSVVSLDIADNSIQTISPNACSNMSNL
QFLNLSQNEMFTFHL--ISHIRSLKLLNLSNNRIHLFSQDTMDMFDDLATVDLTGNLFTC
GCSDL-YIIWLTDTMIRSRWLLYEHYTCLFNNG--TITLSQVDVSQLWDCHKPYIVMASM
II-
FALLVAVLVKLLHYHRWTLQYWYFMFKRAYRRQQELEQNLVKTYDAFVSYHTNSAQW
VYEHLLP-LERDENLKLCIHQRDWIPGQFISEIIVESVKQSRKTLMVVTKEFAESKYCLY
EMQMARNVLFDEGVDALVVVLIDEILSRRINSTLRYLIQRKNYIQWPDEN----FVPKM
KAALARE--
**>Goc-TLRγ1**
VMHYFPSLKYLNMANTNLKMHLKDTNGSFFSKLSNLQTLDISKNKMETFSKKTFCHLT
KL
KHINLKSNNFVAFEL--MNSV-HFLQMDLSDNKISHLTKSNLRLFELMTTINLSDNAFMC
ACEEQEFLKWLKERRIVDIMQHYSKYDCLVTQTQNKVAIRSINMDEFNQCD-
AILMAKVV
AGLVAILVGVLTKVIHYNRFTIRYWKFGIAWMWRRRREEQDQVEYKYDAFVCFQNDDI
DW
IYNELRPNIEQEGAFKLCIHHRDFSPGEFIIDNIVNAIEGSRYAILIISKNFLKSGFTKL
EMQLAMKVMIMRQAEMIIPVMLDDVQHPDMYRALKYHIEKKTCITTNE------HFWEKL
RAALKRE--
**>Goc-TLRγ5**
VFDCLPKLKYLNLANNKF----TNIELELFSNCSNLTYLDVSYNQLKTLPENLVAQTANL

QHLNLGGNQLRQFDV--
LVPVKTLSTLNVSNNLLGYLKDEMCLQLSKMHRLDLGNNPWLC
TCDNVEFLRWVQQS--TNMLLKPDELICNDPHGNFV-LMANINISKLSGCYTTTIITVSA
TIVITLLIILLAVLAYRRRYKIQYIYLIIRAKMRGFR---QDRQYTFDGFMSYSSLDCNW
VTGVLHKTLEDELDYKICIDQRNFMPGSYIAEAIAEGINESKKVILVITQNFLRSGWCTY
EYNMARGELANRGRDCILLIMKDPIPKEHITQTLQTMLESKIYLEWSEDDDKKQLFWRK
L
QDAIGEPQG
**>Goc-TLRβ10**
VMSNCRCLKIVNLKENLLFSY-EKELCRLFKWGKSLQKISIPSNYLAVLPKCIFRGLSQL
TSLYLEKNRLHVIGKGLFTDLINLEFLDLSFNAITYMDSGNFLAMTRLKSLNLNRNNFHC
TCQLLPFRNWIREK--VNLKNFRYNDTQCSLLDRKHVFVHNYTISWLE-CNKVFVVSASS
IGGIFIVVALIVTLLYNYWRDIKYYRMVHKARRHDNRQE--IAEIEFDAYVSYHPEKELW
LRVDLINNLEDDISFKVTFDDR-LEPGRSVIGSMAEAIHKSRKILFVVSRGWLRAITTQL
EIDMALVKMIDDHRDMIIVLLMEHIPKDEMPDKLKMMVKHNTCLKWSDNEKQQAKFWR
DL
KLELGKH--
**>Goc-TLRγ20**
LLHCRQNLTVLKFAQNDLSPVF-NSKQPIFAGCNNLKELDISRNSISEIPNNAFVDLKNV
EEIDLSGNELTNVVV--LENCKQLRVLNLSSNPLTNLDDPTMTVLDKLQTIDLSRITLGC
GCNDVAFVHWAQTT--RVKLFNSDTYKCTYLDSSR--ALMKVSIFSLRACMKDVIIASTV
PTITLILIFIAGLYVYHKRWRIQYHCLLLREVARRYE---QLDELTYDAFVCYCSQDEDW
VAEILRPKLEDELNFKLCIHEREFIPGMDIQDNIVSFMQDSRNTILVLSEHFVESRWCQW
ETRMARNKLLDSPRDNLIMILLQDVLKQKMNPTLKSLIEMKTYLRFPQKAEELPVFWLR
L
KNAMSEHVK
**>Goc-TLRβ9**
VLNHSNAISLINLKGNILFRY-DSQMCSMFKGRANLSHIDISNNYLFRLPSCIFKGLKNL
KTLYLQDNRLTYIQHDIFIDLYKLHRLNLSNNAITLISPITFAPLANLKVLRIGHNNFQC
YCEMKELRNWLGHN--IKKL-GHHKEKCSGPLTRQDEFIHSFTVSWME-
CNGLSTFGSIG
IISIVLLSVITFTVLRHYWRDIQYIKMVRRARKHKSHPEN-NCLIEYDAFVSYHSDKQIW
VIRDLVNELENDVTFRVMFDER-
IDLGTNIFTSMEEAIDKSRKMLFVVSRGWVAAAMNKL
EVDMALVKMIDDHRDMIIVLLMEHIPTNEMPDKLKMMVKHNTCLKWSDTERSRAKFWR
DL
KLELGKH--
**>Goc-TLRβ8**
VLNNSHDIYKINLKGNLLFHYFDYQLCDLFQSKSNLSHIDISKNYLSRIPACMFTGLSKL
HILYLQENRLTYIHKDMFKDLHNLQRLNLSNNAITSIDASAFVPMTLLNRLWINQNNFDC
NCDMKGFRNWLGHN--KKILSGSIKEHCSHPLIRRNEYIHNYNVPWME-CNGLSTISTIT
ISLMVVLSVATLTVLKYIWRDIQYIQMVRRARKHGNYPLG-DIQTEYDAFVSYHVDKQIW
VMRDLVNELENDIQFRIMFDER-
IELGRNIFTSMEEAIDKSRKMLFVVSRGWVAAAMNQQ
EVDMALVKMIDDHRDMIIVLLMEHIPTNEMPDKLKMMVKHNTCLKWSDTERSRAKFWR
DL

KLELGKH--
**>Goc-TLRα5**
-FSSLPALPKLHIANTSLTNIN---------------------------------DYFAKV
TLLDVSNNSISHISQDVLQGMTVLKTLYLHGNKLQYIPEYMMNLK--LDHLSLSDNPWAC
DCKNAWIKPWLNANVS--ITIGFEGIKCHGG----GKQLLHYD--FEAMCNL--VVIGVP
VF-IVIFIILMVAIVTNYRQVLTLMIRHHIF--------EEPAGKTWDAFLGYATDDVEY
VQNVIIPLLE--PKYKLCVHNRDFQPGVPIIDNIAEAVDKSQRTIMILSPNFLQSQWCLS
EFRIAHMQYLNHPSKLLIPILLDDFSPAECTA-IKCHLQAHTYLEAKD------WFDRKL
LQQMPKVSL
**>Goc-TLRβ6**
LLEKSKDLIQLDLSDDMLFTYSDNELCRIFSSQSKLEILTLAGNYFSSLPVCMFQNLHHL
KDLDLRKNRIPLIQRNLFSDLRNLSVLDLRENSITFIDVTDFLKLNKLKTLYLKNNLFAC
TCDLRPFQSWVLGVSQ----KTDGPLRCSSPEQRKNNTVRNFTATWIE-CNELVMYLTIT
LSSILIITSIILLTLYNFRNDIRYRRLLQQVKRKTYTKLQDA-AIKYDAYVSYHV-EKQW
FMEEISKKLEKEIQFNL-IHDDNIVAGESIFGSMSKAINRSYNIIFVISRGWIHDPARAI
EIDEVNGILNREKRHNIILLIMEHIPPEQIPGNLKMMLRNNVVLYWNEDPKRQRIFWRDL
ILELGKTKD
**>Goc-TLRγ15**
FINCFPSLVELNLSGNQ-----GMIDVVMFDNCTNLRVLDISNNKLASLPYDMLHHVPYL
EKLNMAGNFFTHLDILNFMETKTLQSLNISSNHWRTFPDTWQKVISGLVELDIHGNPLV
C
TCDTIDHLIWLQNI--QVALYKPSTLTCMDLYGQEH-NIMDINMYKYRDCL-PYLMAGFI
PCTVTLVLIGLILYAYRRRYRLLYWLLELQAKLR--QPHAEERNFVYDCFISYSSNDIDW
MIEMFQKL--EQHNYKMCIDMKDFRPGSPLVDEINQGIIQSRKVILIITQSFLTSGWCNY
EMDIAHGELALRGEDCLILVLKEPRPQALITPALQRLLEERIYLEWSNDHDRQAVFWRR
V
QDALGEPLQ
**>Goc-TLRγ19**
FLACRPNLTIALLSQNNLSPIFDTPKQTIFHGCSKLKVIDISENQIKTIPFSTFIDLIQV
EIINISGNYLRQLDV--LELCTSLNLLNLSSNLLTTLSSRMTSSLDTLQTLDLTHNSLMC
GCNDIAFIDWAQQT--NVRLHNGHRYTCTDKDSHQ--QLLDISVYHLSECKKEILIASIV
PSVGLVIIFSIGLLIYNKRWRLQYRYLVAREMVRGYV---EIINLPYDAFVCYCSQDQTW
VAEQLRTKLEDEFNFKLCIHDRDFIPGMDIQENIVKRLEESRNTILVMSQHFVESRWCQ
W
EARLARNKLLDSPRDNLIMILLDDVLKGKMNRTLKSLLEMKTYLQYPHNEGEKQLFWM
RL
RNVLMENRR
**>Goc-TLRα4**
-LTQLPTIPPLLLSKNSVSRFE-------------------------------PYLANV
SVLDLSWNGLHEIVIEALNSTRDIEQLFLDNNALTELPKSIKDMPPMLQLITMHGNHFRC
ECESAWMKSWLQAQVDNGRVNSSLKIQCADTQT----EIIH-D--FHELCL-NFLVVGVP
LA-LVSLIVFAFAVLYKFREVIILTIR----KK----PTEKPEGKLYDAFIGYATEDVLW
VQDVLIPILE--PQYKLCVHNRDFVPGTPILDNISEGIEKSQRSIMVLSPKFLDSHWCLE
EFLQAHRQYMAHSSQILIPILLDDFEPDNIVAYIRCYLQSHTYLQHAD------LFARKL
RIHMPKLTV
**>Goc-TLRγ4**

IFNCFTALRYLDIANNQL----KSKDSVFFANCSHVEHVDLSSCTIGEVPRHLLAQLPNI
TYFSLSGNRLRKLDI--LNK--KLNLLNVSSNLLSAISPGMLSQLTHLHTLDMSSNPLQC
MCDTTTFMTWVQQS--TELLHNPSDLLCMTSAGDMV-AIVHVDVAAIHQCILPTILATTL
TTTGILGIIISLLIYRKRYRIQYIYLIIRSKLS---ES-KQARFPFDAFISYSSLDSRW
VVNTLYSTLADTHAYNVCIDQRNFMPGAYIADAIVEGINDSNKVILVISQNFLRSGWCVF
EMNIANGELANRGRDCLILIIKDPIPQELITKTLQALLESKVYLEWSEDPDRQRVFWLKL
MNAIGPKRD
**>Goc-TLRγ6**
----FPKLKTLQLRNNKV------SNVEMFANCTALAHIDLSHNGIYSLPLQLFRHTPNV
NYVNLAGNKLHTLA-FEVVFLTNLALLNLSNNNIQILTRNFQENVDQMSFIDLDNNALIC
TCDNVDFVRWIQGS--WHFLLQSNQIECKDGGGVS-HSIIDIDVNHFHGCIRSTLIASLV
PNVFFILCILLGVLFYRKRHKLHYLYLLARAQMRQRRNV-
DRGNYCFDGFISYSSLDTDW
VIEHVYNELADHHGYNICIDVRNFMPGEFIADVIIESINQSYKVILVISENFLRSGWCTY
ELNMARGELSIRGRDCLVLIFKQPIPRELITPTLRSLMETRVYLEWCHEADKQQVFWRK
L
LDALGQPRQ
**>Goc-TLRβ7**
------------------------------------------------------------
----MKENRLTYIHKDMFKDLHNLQRLNLSNNAITSIDASAFVPMTLLNRLWINQNNFDC
NCDMKGFRNWLGHN--KKILSGSIKEHCSHPLIRRNEYIHNYNVPWME-CNGLSTISTIT
ISLMVVLSVATLTVLKYIWRDIQYIQMVRRARKHGNYPLG-DIQTEYDAFVSYHVDKQIW
VMRDLVNELENDIQFRIMFDER-
IELGRNIFTSMEEAIDKSRKMLFVVSRGWVAAAMNKL
EVDMALVKMIDDHRDMIIVLLMEHIPTNEMPDKLKMMVKHNTCLKWSDTERSRAKFWR
DLKLELGKH--
**>Goc-TLRβ4**
FFTDFPSLKMLDMSNNDFSFL-HQVLAKMILGFENLEILRLSNCQLSDVP-NFFDGGEKV
TELDLSWNLIGHFRNGVLNKLRSLRKLYLQHNHITHIDPLNFERMNDLQYLDMTNNRFT
C
DCEQREFINFVKGNQ-HRIIFHGRRTKCH-PDSAYNTFLLHYTSPWIE-CDGNFIMILSF
TIILVVIITCIFIFLYNYT-SIRYRSALCKIKCNQYKSL--GHQYDFDAYVIHHPTDISW
ILYELIPHVEQHFSFELCIEERNFLPGPFKTDNLARAILRSRRALLIISKDFLESDWFRL
ELEMAQLQHLNGREKYIIIIFLDEIPASQLPMKLKCLMGFTTSFVWPKNRSKRNEFWRG
LLLELNKKPP
**>Goc-TLRγ22**
FFDHFPKLKTLLLGNNELGLLFASDDFLFFRNSTTLESIDLAGNNLSRIPPLLFKDTKNL
QYLNISNNYLDSFNI--LSTLSHLKYILLSNNKIRVLSSTTRHQINTLAHVDLSGNPLLC
DCNNLDFLHWLRDA--PLVFDNKDSYQCTGMNNFR--KVYDINVKDFEQCKINMIKTIAS
TAGTALITMAVVIAVYRKRYRLEYLWLVSKATVKKREGNDENGRYIYHGFVSYSGRDD
LW
ICDQLHIHMEQVMGLSLCLHDRDFIPGEFITDNIIKSMEASRKTIIILSNNFLESRWCEF
ELQMAESRQAEMTYNTVITILLHDVNQNKIGPLLKKYLKQKTYLAWPRDHHQRPAFWL
RLKDAIDREPD
>Goc-TLRβ5
LFRS-SNLSEVSFSGTYVGHSDVTLLKDIFHGHHNLQTLRLDTTHIQELKSGTFWSLTRL

EVLILTSNHLKTLPTDIFQGLVSLQYLDLEDNDFIHLDPNMFTQLPNLKLLWISSNSYHC
GCDLRPYREWLNHS--KVVL---PPGRCYSPSKLNNQIVDEFSLPWLE-CDDTLLISGTL
AG--
LVIILSHGYAVWRFRFDMKYWYYITRAKRAAEQPLQDGGNIQWDAYVTCTPQDQKF
IYDHVIPNLEEDFKFKLCYGPRDFLGGSEI-GNRENALNNSHRAIFVISKEFMKNNWGKF
ELEMTQLKLFDDHKYMTILFFMETIPKSEMPELLKLLKRHSLCLYWKSENREQNVLWK
RLKLDLFKARQ
**>Goc-TLRγ14**
VVNCFPSLKYLYLSGNNV----QGHSPRMFENCSKLKVLDISQNKLQVIPFHTFNETPNL
EEIHLSGNYFSDLEILDFKDTKSLRLLNLSHNQFHALPEFWKDTIAEFKHLDIHGNPLVC
SCDTVDHLLWLQSI--RPLLYDADNLTCKNNQGRQ--FIMEINISEFKECIKPLLLAGCI
PSAIIVITLTLALCIYRRRYRLHYLTLILRARLRKYINSEQRQDFLYDSFISYSSLDVTW
MVDILYKNLSERLNYELCIDVQNFRPGEAIVDEILAGVLESKKIILVISQNFLRSGWCNY
ELKIANGELALRGEECLILILKEPLPKELITPTLRRLLKSRIYIEWNDIEDRQQLFWRRLQD
AIGEPMA
**>Goc-TLRβ3**
SFKGLLSLEKLSLARNHLDNS-DGTIVTLLKSLPRIKELDLSWNHLTYIPKSSLDSMENL
TKLDVSGNRLTSFTVDKVRTNTKLNQLNVSRNALASIEALEIGKVTLLNNLDIRYNKFKC
GCELRYLRNWLLEKNHKFSL---YSEKCFAPIEMINTSVMDFKINWIY-CDHLIISLSSA
GG-FLAIILFCIVLSFVFYWDIKYWWALRRKGLGGYLPLDEESKLSYDAFVSYHTSSESW
VADKMVKNLEDDVNFKLCLHGRDFLPGRYIADNIIVTMRNSAKIIFIITQKFLESQWCGY
ELEQAHIRQFDEEKHLVILIFLEKIPKAKLPKKIRLLMRHVTYLEWDKTSRAQNLFWKKLK
LCLLDKPT
**>Goc-TLRβ2**
SFKGLANLEKLSLARNYIGSS-DDIIVMLLHLFPKVKDLDLSSNHLTWIPKTALDAMKDL
TILDFSVNRLTSFPLENVNHNTKLERLNVSRNALVNIEAPQIKADTKLNNLDIRYNKFFC
GCDLRPVRDWLLAQRFKVSI---FSERCSAPTELRGTPILYYKINWIN-CDSLIISLSST
GG-
FLVIVILCIVTVCIFYWDIKYWWALRKRGVRGYIPLDQQVQLSYDAFVSYQTSSQEW
VAEYMTKHLEDDVNFKLCFHGRDFLPGRYIADNIIVSMRNSAKIIFVITQQFLESQWCGY
ELEQAHIRQFDEEKHLVILIFLEKVPKSKLPKKIRLLMRHVTYLEWDNSSRAQNLFWKKL
KLSLLDKPT
**>Goc-TLRβ1**
SFQGLSSLKDLNLARNNLDFSVQSLLSKMFSALKTLKKLNVAFNHFMSLPGDVFDGLQ
SL
EYLDLSMNKLSFFGKRYIRNVKSLRVLNLRRNMIIKFDMNILKPPYKLVLLNISENKFIC
SCDLRGFRDWLDSKPKHITI---FDQNCSDPDTMKTSRINDFTIPWID-CDNLTISFVSA
GG-FLIIFSVTVVVLVHFRWELKYWWFLRLSRRRNYIQL-
HDDGFQYDAFVSYHEESSRW
VYDYMPKELEDDMSFQLCFHGRDFIPGQSIQTNIANSISQSRKIIFVITQGFLDSNWCTY
ELEMANIHQFDKKKNLIILIFLENIPKYKLPKKVKLLMKNVTYAEWEENNRSQRIFWKRM
KMALMDQPT
**>Goc-TLRγ12**
FFDCFEKLRVLNIANNDI----VNLTFTVFTGCNRLEYFDASYNNLKDIPTSAFQQVLNI
KQLVLSGNHLRDFDA--LQGLNNLQTLNLSDNILTNLPLQVRNSLDDIASIDLYGNRLSC
SCETIYFIEWLHNF--KENVYKYEGLECSYIDGVF--PVASVSIFRLWHCWAPLIMSIVS

----TIAAILFIFAIYKSRYKIQYRYLIVKGKFKRYQAAPSHDNINFDAFVSYSDHDIIW
SVETLYCKLAHEWRHNVCIEGRSYRPGGFRNEVVMEGINESNHILLVISQSFLKSGWC
AF
ETRIAHGELVHRGKSCVMLILKEPKPESLIGTVLRSLLDNGCYIEWSNNPDKQRLFWYK
LQDFLGEPIN
**>Goc-TLRγ2**
AFCFLSNLKMLVLNDCKLNDILTDTETSLFKNLLMLESLQLRYNNLTYFNVSHLNSLSHL
KHLDLSQNKLSKFDFAQVSK-PNFTNLDLSSNRFRMFSSDSMKSLDAIKTINLHDNILTC
ACNMLKFMLWMHNN--
KGHMVQYSTYQCVFSHNDTEMNLIDIDMVSFSECHKDYIIFSVI
SVIITLLCCLIGLILYKKRWTLRYWYFILRQSWRRRRDA-DVMHYHFDAFIVFHSDDYEW
ILNELLKQSEEPHGLKYCIHLRDWRPGNFVSENIVQSVERSRHTVLIVSKNFTKSKFCY
Y
EMNVARSLLTSHGRDVIIAILIDEIRKAGTTATLREILRQKTYLQWPSEN-----
FWERYHTMMDDNEE
**>Goc-TLRγ8**
FLTCFESVETLIIGKNYV----NGREFNVFQNCSKLHYLDLSFNGLTGIPWRAFNETPSL
IALNLSGNQISYPRF--LEAASNLTSLDLSNNKIHYMDESMRTNIGTLMSLNLDNNILLC
NCNGITFIEWLQKY--KQNIVNWDKLKCIDSKGTEK-KLIDINITLKKECLMDIIIASST
VSGIFLLVVFIAVCTYRRRYKLQYLTLLLRAWCR---KPDDGDQYNFDCFISYSSLDRIW
TLETLYTTLATKHGYNICFDERNFMPGQHLVDIINESIFTSRKIILVITQNFLRSGWCLY
EMKMARGELAARGRDCLILIMKDPVPKDLITPTLRQLLDSRIYLEWNEDVDRQQLFWR
KLRDVLGEARH
**> Goc-TLRγ21**
VFHCWPNVKTLLTGGNDLSHVFYEENIDYFENCTHLEVLDLANNKINKVSKNLVKTAIN
LRQLNISRNQLSMFDV--LDNNQQLELMNLSNNILGSLPISLTLTLDHINIVDLQNNPLLC
SCATLDFISWCQTT--KVHLNNLPSYTCTDGTQRQ--YLMDVSVSHVDKCKEPVVIAAVS
TSLITIIFVIIAITIYTKRWSLNYYLLLSKIFLRRRNRYSENTSYRYDAFVSYSSNDDDW
ILRNLHPILEDEHGLKLCLHQRDFIVGNDIQDNIIESIEASRKTIVVLSNNFLESKWCYF
ELQMARNKVLDGKLDLLVLVLLDDLAKDLVTATLRTLLATKTYLRAPREPQQEALFWLK
LKDAISTDRS
**>Goc-TLRα3**
-LDKFPKLPTVNVRDNHITILP--------------------------------SYMPMI
KVLDASYNSIHNINIDSLSNLTALEIFLINDNDLNYLPNTWGSQTSSLKKLCYHDNPFHC
DCSSLWMEKWTTTFED-LLCSE-D-IVCQNG-----RPITDQG--INEFCHI---FASVI
IGILVTTIIV-TFLLFYFRQVIVLWTRR---RC-----ADDPANKLYDAFVCYADDDFDW
VNDNIIEHLE--PKWKMYIPDRDIQPGELRVILIQEVIETSQRTIMILSQNFLQCTELVN
TFRFAHQQYMVDPSKVLIPILAPNFEVGTMPLFVSCYLKAHTYLEASD------FFMRKL
QLQMPKIRV
**>Goc-TLRγ13**
VLNCFHSLIELNISGNVI----GQSKWKMFQNCTQMTRLDMSFNKLTSVPLGAFNELSGL
KDLSLAGNDLRSIDI--IQNNAELRFLNFSHNRLDTIPDKWREYLDEVKILDIRNNPFIC
SCQTIEYVEWFQRR--KSMIFQPNALSCVDITGKH--SLMDIDIAEYEECFHPFIIAGTI
PSVVIIIILLIALFIYKRRYRLQYLSLVLKAKMKNYVNAKESKTYLYDSFISYSSNDVYW
MVETLHNKLDDELGYKLCIDVRDFRPGNPIGDEIETGILQSRKIILVITESFLRSGWCTY

ELNIANGELALRGEECLILILREPRPEQLITPTLQRLLKSRIYLQWPEEEDKRLVFWQKL
QDALGQPYS
**>Goc-TLRα1**
-LSSVPGLP-LNMTGNNFGNITNCYLNDVFNGIYNLKTLTLKDNFITTLTSGPFKSLVFL
DTLDVSNNILNTISDNAFSNVRLLKHINMVNNNMTVLKPEVFNDLSKTGTMFLSGNPYN
C
-CESLPLKNWLNTHQA--QIIDIDITLCQKE--TDTVPIYSMA--DANTCDLTFYVIVII
LS--LLFIIICCILVVRFRQAIRLRLYWFKWRF---EAFEDDSKKKYDAFISYTGHDGDW
VREDLLNFLEG-NNFNICLHERDFRAGELIIDNIDRAIEDSKRSIIVLSNNFLNQDYTMY
EFEASYRDWKLGLRDPVIVILYEALNKEKIAEHLQTHLNTRTYIDKSK------YFWENL
LLAMPRQKH
**>Goc-TLRγ7**
VFNCFEKIKIINFAQNKV----YNGDLELFLNCTYLEVLDLSFNALNSVPKNTFTNLVNL
KKLNLAGNKFIHID-
FDLRRFWKLESLNMSRNSMDMLSVRTRKELTDLYTVDLHGNHFSC
KCADIDFVEWIQNN--FGMVAMPDSLPCSDERYVKR-RIMLISVTHLRRCWSHTILAAVV
PVALLLLLLFVVLLAYQRRYKIKYLYLLLRAKLR---DHQDRQVYLFDGFLSYSSLDQNW
AVN-LCEKLEQDFGYNLCVDQRNFGLGYQLVDLIVESINQSRKIILVISQNFLRSGWCLF
EMNMANGELAARGRDCLLLVLKDPVPQELISPSLRALLDTRLYLEWSQDPDQEQLFW
QKLRDALGDPRP
**>Goc-TLRγ11**
MLDCLQSLTVLKMRENQLEVVSNNLNNTLFNNCTQLEYVDLSFNGLTFLPKDSFIGTH
RMKTLNLSGNRLQHSDM--
FPKWQYLDALDLSMNYFTTIGEELRRQMEYLKTLNIQGNPFSCDCQSLDFIEWIQNT--
QVHVTNRDGLSCSLSNQINTIKVLSLNIGQAKKCWSPIILAAVVPCVILIVMITTITIIYRSR
HRIQYYYLIIRAKLR---EQKVQDDYHFDAFLGYSSKDVGW
TIDILYKTLANEMGYNICIDQRNFRPGNYIADTIVASIAQSNKVILVITQNFLQSGWCNF
EMNMAHGELGARGRDCLILILKEPQPENLITPTLKALLGTRVYLEWSDDPDRQRVFWR
LLQDALGQPKP
**>Goc-TLRγ18**
VIHCLPNLKTLLVSENNLSSLFGKYNQTVFYGCSQLQTLDLSKTLIQAIPSDAFKDLLNI
EHLKIAGNDIQTFDV--LEQCKSLILLDLSSNLLSKLSERMMSRLDALQTVDLMHNPLSC
GCRDIAFITWFQNT--HVHFLNGQEYTCTDENYKE--HLSDIYVFHLNECKMDIIIASTV
PSVCLILIFSIGLLIYRKRWQLHYRYLIAREMVQAFT---TMGNYTYDAFVCYSSQDQTW
VYEQLWTTLEEKHGLKLCIHERDFMPGVDIQENIVQSLEESRNTILVLSKYFVESRWCQ
WEARLARNKLLESPRDNLIMVLLDDVLKPKMNGTLRSLLEMKTYLQFPACPDQQKLFW
MRLKNAISEERP
**>Efe-TLRα**
----FPNIHSIHFEGNALTTFMYNDLPYAFLPFPNIEFLSLNGNRITHLKFGTFAKLKKL
TNLYLHNNLIKEIDSSVFDDLHLLNQLTLHSNSLELLENDTMDLLSSLSNLTLDDNPWVC
PCDNATFKYWIQQHSE--IISSPFSLKCNE----T---ILRI---DEDLCYKQYLTGPLY
TASLFCLLLLTLVLVYRYRYIIQVIVYKFELRR----KQQESESCAYDAIAVYDSSNLKW
IKDILIPRLE--PKFKLYILDRDMLPGSVQCNEVVENIKRSRRTLVVLSGAEDLQE-IGF
GFDVAHHRVTQERHHRLKLILLHNVAKQDLHANFKAYLTTGQYFSVVD------LFWQKM
LYFLPR-PP
**>Efe-TLRβ1**

IFQPLLNLQTLNLAGNRLGRVITDIDGQLFRGLSKLRWLRLDNNEMRVMQYTMFSDLTS
L
QMLNLSNNYLSSWAPGIFNASGKLSIVDFSSNKISIVTEQELLTVPTNTSLNLTDNPFAC
YCDLIWFRRWIHNS--STKLAHLQSYICNSPDQMAKKPLLEFNPDDIARCHLMWILLGAC
SG-VVAIMLFFGTMMYRYRWQLRLRLYYAQRRIRGYIEVDEGYD--
YDIYVSYSDSDREW
VRTELMPRFNLLGELRVFIEEADATFGFLEFDTLAEAIYKSKKIMLVVSDEYLHDGRRLF
EREWAIRSRFEKQDDSIIVVCLEPDADVKVPAVLLPIC-
RNQGLEWKQDEAGQEFFWRKL
ADVIQY-RD
**>Efe-TLRβ8**
IFQEAQNIEKINMSFSRI----ESNLIAPFHHLLKLTDLDVSGTGISNLS-HMFADLPNL
KRLRLSRNPILTLERKDFEGIESLRDLDLSSGKLSTISFESLKVWKNLRVVDFSDNPFLC
DCNFIWLRRWLKRANAKVEVRGWDKYYCQTSKG--QVNMFQLSPSDID-
CFQHWLLTVRL
LTSLIWITATSASALHRFRWHLRYWYFMKTVHAQRFKDEEEPDFFAFDAFVGYSSSDS
NW
VITQLLPRLEQECNLRLCIHERDWLPGRDIAENILESIDNSRKTLLIVSNAFAVSHWCHF
EMTMAQTKLFEDDRDNLILVLLEEIADCNMNPRLQLLMQNKTYIEWTDNNIGQQLFWA
RLRQVLAKQSN
**>Efe-TLRβ7**
VFRGLSHLRVFNISRSKS------KTIYPFYYFRRIEVLILRDVGLRNLER--ARYNRRL
RYLDISQNEISVLMSSSLARL-KLEVLLVRDNWLSVIDLETLSTWTNLKRVDFSENTLNC
DCKIIWFRRWLQNRQSNVTVENLHLTQCTAPAKAKDKPVHLLEPTDLE-
CFKPYMVAVFL
TAFFAYLIAPTVSMLHRLRWILKYWYFKHKTKAKEYRDIDDNKPYEFDAFISYSESDRN
W
VVSQLRPRLENEFGLRLCIHHRDWLVGRDIVDNVVDSIEHSRKTVLIVSNAFALSPWCH
F
ELTMAQTRLMEEDRDSLVLILLEEIADCNLTPRLQIQMQRRTYIEWTKQSVGQQLFWA
NLKHALAKPSD
**>Efe-TLRβ2**
----F--LEPQYSSNNPLNN--------------SMTSLDLSGTNLNTLLNGFIECYKAL
RTLNVNQNKLEILTIETFRNLPSLEELRAANNFLTQISESSLKLWTQLKTVDLSGNPFRC
DCKLLKFSQWIKENNFKKLL---DNMQCVSYSSGKRQSIVS-AENRLKLCLEWFLWALTA
TVFVTSFLSTFASIVHRFRWNIRYWIFSHKIKTRRFKQVRDKKHYTYDAFISYSETDSRW
VILQLLPRLESEYHLRLCIHQRDWLAGRDIAENIVLSIEQSRKTVLIVSNAFAVSQWCHF
EMTMAQSRVFQDDRDNLILVMLEEIPDCNMSPRLRMLTERQTYVQWDDHALGQQLF
WVKLQQALAKPAE
**>Efe-TLRγ2**
LFR-ISNITSLVLINNDF----QN----------SLRPLDLSGSGTSFIPKSMFSELSAL
QFLNLSRNIIESFQV--LPPSGNLSLLNLSDNSIRILTSEMISELESLQTIDLSRNPLSC
LCNATEFIAWLKSTK-KVSFENIEEYTCLHPNGTK--SVNSLNMTELMDCKKSFFIVPLI
IGLLLLLLGVALIVIYKRRMDLRPYIMFIRYLS----PA-ESDNYEFDVFVYFHADDVLW
VRGALLEKLQ----HLKVITPDNFRIGASMADAILDGCRKSRVIVLVLSSSFKRDDWC--
-----TLRSYTSHPGSVIPVVINDTDLSDFEDDYSNLIATHS-IDLAD------FIWNEF

TSRVDRQSS
**>Efe-TLRγ1**
LFR-ISNITSLVLINNDF----QN----------SLRPLDLSGSGTSFIPKSMFSELSAL
QFLNLSRNIIESFQV--LPPSGNLSLLNLSDNSIRILTSEMISELESLQTIDLSRNPLSC
LCNATEFIAWLKSTK-KVSFENIEEYTCLHPNGTK--SVNSLNMTELMDCKRSYFIVPLI
IGLLLLLLGIALIVIYKRRMDLRPYIMFIRYLS----PA-ESDSCEFDAFVYFQDDDEPW
VSRVLLEKLQ----HVKVITPDNFPLGAAMVDAILDGCRKSRVVVLVLSSSFKRDDWC--
-----TLRSYTSHPGSVIPVVINDTDLSDFEDDYSNLIATHS-IDLAD------FIWNEF
TSRVDRQRS
**>Efe-TLRβ6**
IFNRVPNLITLNIARSRA------KSLSLLFGLKRLEVLNMRATGVTSLPY--IAKRRHL
RILDVSENEINVLQSPVLQNL-KLEVLLLRDNWLSVINITTLDTWDRLTRVDFSENTLYC
DCQVVWFRRWLRKRK-NTTVENLHLTRCTGPAEVKDVPIHLLHPTDLE-CFSAYLLSVFI
AVFMTTLVTFLVAILHRLRWMLKYWYFRYKARAKEFRELLDQQHYEFDAFLSYSETNY
EW
VVDQLHPRLENEFGLRLCIHHRDWSLGSDIVDNIVNSIERSRKTVLIVSNAFAVSQWCH
F
EMTMAQTKLFEDDRDNLILVLL-----------------------------------------
---------
**>Efe-TLRβ5**
LLWGIPKLKELNVSRSVG------RSIAPLTNQTSLEVLSVRQDDLFDEDVNMLTNVPHL
RHLDLTDNNINVLENIPFHKLKSLEVLLLGKNWLITVNATSLCTRTHLNRVDFSANPLIC
DCGIVWFRRWLNTT--QVIVDNREDIRCSAPEEVKNRILSEIHPTDLE-CFQELLALGLI
LVQMVYLLSLIVSVLYRFRWRLKYWYFRHKTQGNEFENWIETPHYDYDAFISYNESDS
KW
IVTQLSPRLETEYHLRLCIRERDWLVGLDIVDNIVDSIEKSRKTVLIVSNAFALSPWCHF
ELTMAQTRLMEEDRDSLVLILLEEIADCNLTPRLQIQMQRRTYIEWTKQSVGQQLFWA
NLKHALAKPSD
**>Efe-TLRβ4**
IYEGMPNLEQLRLANSRR------KAAIPFSSLSNLVELNLRGVGIRAVHLNITKNLPKL
RRLDLSDNELSHVRASFFRNL-ELDILLLRRNWISTISKTTFGMWSRLKEVDFSENPYFC
DCRMVWLRQWLRNKTRATAVRNKGRMVCIGPHEAKNTKLYLLKPTREE-
CFAYYRLAVLL
FTLVIWFLAPLLSIVHRYRWFLKYYYFKYKIQQRQLHDLLDKKQYSFDAFISYSESDSKW
VINQLRPHLETEINLHLCIHHRDWLVGRDIVDNIVDSIEKSRKTVLIVSNAFALSPWCHF
ELTMAQTRLMEEDRDSLVLILLEEIADCNLTPRLQIQMQRRTYIEWTKQSVGQQLFWA
NLKHALAKPSD
**>Efe-TLRβ3**
VFNCIPNLEELLLINVQV------SVRHPFRNLTKLKKLNMGGTSLGDVEVKMIPDLRQL
EWLSLAHNTIQKLRRGMFQNFKTLKVLSLGNNRITTLNVTSLSLWKSLERIDLSGNPFT
CDCQLVWFLHWLSTT--NVTVSDEKQYQCNSPAALKRKSLKKLHPNEVE-
CFEWWLLAVLL
ISITASASSTIGSVLYRFRWYVKYWYFKYKIQQRQEALSTDNHSYQYDAFVSYSKHDTK
WVVTELRRHLEIEEGLNLCIHDRDFLVGEDIVSNVISSIEQSRKVLFIVSNAFAASQWCH
FELIMVQTRMLENDRDNLVLILLEEIDDATLSPRLKLQMEKQTYLEWTSSEVGRQLFWD
RLRQAVSRPPE

**>Hro-TLRα3**
-LKTLPLLP-LTFSRNMISSMD--------------------------------FYLSST
TVLDLSYNELNQFDLSTLMFSTTLEELYLHSNNLVSVPPEFLQKSPKLKVLTLHDNPWD
C
SYENKFLKSWMMNGNV--SLLHENSILCRTP--LSGKSIFFVK--DEEANDP-RNRVILC
ILIPLFAIEIFALAIFLILKKFKVKLYYLNIHPR----ECTDEYMEYDAFISCAFSDRCR
AIE-LVSTLEN-RGYKVCYPERDFIPGEPTTSFVSK----SRRVIYLLTLDFVNTPRCLF
EFQISLQRNLEVKHKRIIVLLDSSLKVQLLPNDMFNFLTTHHCIDLLK------NWTNQL
FYSLPIKPL
**>Hro-TLRα1**
-YESLPLIPLLYFNKNLLTSFN--------------------------------HVFNDT
KILDLSENKINEISPHTWVQLIQEDSVFLHNNNLSYLPRILEKMNTS-KRVTLHGNPWSC
TCQNAWMLDWLKSHVH--VV-KPEKMVCAYP--HESKSLFEVDF-----CYS---ISCVT
VGIVVLIVAVT-----------YVWYRRFGPQKK--PPVPPNPKLTNDVFIFCSDEEQVP
LVKEIIRWLENEHRFSTICGLRDFDSK-PKVVNINEALTTSKRIIFIVSKDFLKDNWCIS
GTMSA-FSLVEDKRRFIVIFCGVHLQSTNIPVELEIYIRTYTYLSFDD------SFWKKL
LRAMPKE--
**>Hro-TLRα2**
-MSNLPMLP-LTFSHNSIETID--------------------------------FYFNNT
VVLDLSYNKITKIDLQVFKSLKVLQELHLHSNFLTTVPRDFLKNDRMLKHISLHNNSWDC
SCGNKWLKQWMMNQSI--TLLTPDSVLCRTP--LSGRSLFSVS--EEECPKP--SRLLIS
LLIPLLTGVLLLLALFVLIKKFKVELNYLNIHLR----ECIGENMIYDAFVSCSYSDRRR
GIE-LVRLMEG-KGYHVCYHEKDFIGGQSIAANIVEAITFSKRVVCLLTSNFLKSTYCMF
EFQTSLHRNIELKRKRLIVLLDESVEVEVLPNDVHNFLTTHTYIELSS------KWTHQL
FYSLPLNPI
**>Hro-TLRγ1**
FLNFIS-LGELNVEKNELGEQMSDMFGLTFQCYSNLTILNLSNNKIKKLHKLSFKNLRQL
RILNLAENSLQTIE-FEISHMKYLQYLDLSRNLLVSLADDTCYVLSGLSSTSLYGNPLQC
NCETIKFMKWVQSK--KVTINNKNHTNCQNRSS-KFITLSNLAVNELRFCNLPLIIGGSL
VG-
LLIICSFLGFVLYKYRWDIRYFLMSMQKSKRRCTSFERNRYYKYDAFVCYEKSDRRW
VVTELLYNLEADDRFLLCIHDRDFELGLGIKHNISMAIHASRKTLLVLTNNFLKSKWCRH
ELEMASLESLDRECNLVVPVFLEPV---ETWDSLSWLTKRYTYLEW--------FAYDKL
VVALN----
**> Cgi-TLRγ2**
AFASLRKLEYLDISNNEH----------------GLAVLGLNQTNIKVLKVNI-------
----LQRHHLSYLDN------TSLLELSIATNRIETLEPCVLSRLKSIKRLSIARNRLIA
AAYVLEYHSLV-----NVEVINA--------------SLRNFTIS---GCKE--------
---LILKNTVYHPALYRNRWKIRYMRYTLFQRAR---SSSSDDLFLYDAFVSYTSKDRDF
VIKDMIQKLEQDNGVQLLIRDRSFIPGEFKCQQIVRSIQESRKTICVVSKRYLKSAWRDY
ELNMARVEGVEVRKRYVILILLPEVCSGGYPKKISDFLKRDCFIEYPDDPAGYEEFWQR
LCSALQENQD
**>Cgi-TLRα4**
-LTKIPGFP-VSLDRNNISEIYHSTLNNSFIGLLQLKTLYLNNNELQEINRGVFNKLWNL
TELHLEYNNIAYIEEGAFSALTSLSTLFLDHNLLISLPQSATNHF--LSNIRLGENPWSC
SCDVMAFIPMVMNRSM--VISDYSDMFCKETGE--NFSMKDVLVK---RCTSNILKILVI

VAAIILTTFFIIIVICLWR---PIVLFHRKCKCR---RYPEDGDKSFDAFLAYSHKDDDY
VTREFIPRLENELKYRLCVYYRDFPIGGTIADTVASSINRSKRTILLVSKHFNDHEWRNT
AFQHSFGGLFKQKDNHLIIVLLDDAKGMKLDRQLKVLVKSHHVISYRD------CFWEQL
QYKMGSSKR
**>Cgi-TLRα3**
-LYNIPSLRH------------------------------------------WRLNFPKL
KVLDLTNNHISDLIIDHDPDSSDKGVINLQYNNLTSVSDNQLKNFHSF-YIDVQNNPFSC
GC--MRVKHFILNNTKSSEYSYLRGLKCQNP--VAGRELITLS--DADGCGSQ-SGPIII
LCVLVFVLFVCLVVIIRYRVEIKILAFRFNI---PCQQQDNLDNKKFDAFVAYSQQDSDW
VLKNLVWQLETLQRFHLCLHQRDFTVGAPIAENIINSIERSRHTILVISSNFVRSEWCLM
EFRTAFHQSLIEKRRHMIIIVMGDLPHGELDTDIKRCLKTLTYLETHD------LFWDKL
VYALSDKQR
**>Cgi-TLRα1**
GFQDMPNLEMFGILDRKI--------NEIFGSFQKLHTANFSDANLEFIPANWFRQFRSL
RIIDLSHNRIKEIPYRR-NHFGKLIKLILRHNNISRITKTLIEKLSNM-AVDFSKNKFVC
ACESLEVLQFVRNQIEAINYHYLANETCYYPSSLQGMPLRSL---DSLLCPNSWTFQELY
IGLIVLTFITIVCLVVKFRKEIKILTYRLGIRF-PHR---SGRLKEYDAFVSYSALDESW
VMGTLCKRLEG-PPLRLCLHHKHFVLGACISDNIIESVEKSRHTIIVLSQNFLQSEWCLL
EFRKAFHQTLLERRRHLIVILMDQINLDTLEPEMNYFLQSHTYLKRTD------LFWDRL
IYAVSDP--
**>Cgi-TLRα2**
SFKNLQ-LETFLTESNRF----------------ELRILNISHSSLYYVPENWIIYFPKL
EYLDMSHNKIQDIVLSMYDPTSARLTLDLTFNDIRQISVRFLEKIARL-YVIIDNNPINC
SCTDMRVLEYIRNSVK----QYIRDLKCQFPENIKGRRLRDL---DNDGCG-KMLPIIVV
LSILICFLLIFLFLIIRYRLQIRLFCARLSGISN---DMDAEKSFKFDALICHGLFDEEW
ARSTFIENRHK-
SHLKLGFYREDATDQKNNFEKLIDQMKSSKYVVVLLSRQFLEGEFLTP
GFQEALQQSNEHTRKRSILVLMDDIPTQEETICLRRSLQTFTCIHKND------RFTDKF
LYLLSSK--
**>Cgi-TLRδ2**
ILQGVKNIRQLRAVNVQFN--FNLISESLFKNLKYLTNLDISRNSLNFLPQSLRDQKLSL
KELNLDHNMFSSLS-SSLKQFTNLRKLYVRYNLISKINEKDQELFKSLTLIYIEGNPISC
TCSNIQSLKWMKDH--QHLFSDLSKTKCVGSNNL-TVELNEW-LLKFEICQADWLIFSIV
LIVSTLTMLIILAAIKKYHVHLEYVILRVKQRLMPVGHVCVEGDFQYDVYISYNDDDTSW
VANNLNPKLE---NIKAWFKEKDSIPGGWESEEIVNCINDSRKVMFIVSESFLDKGWHSY
AVQMAITHAFHNQRRSIMVIIKDGLPLERLPKEFKHIWWCIEHLRWPEDETNDETLLLNL
SNVLVSE--
>**Cgi-TLRδ1**
MFQGVRNLHHLYVLDVGLNNTAHSISNSLFKNLKNLLTLDFSKNGLAFLPSLLMDQKHS
L
TEIRLDHNRFSAVP-SVLTELKELKTLYVRFNLISKFSRNDQRLFQSLSSIYIEGNPITC
ACTGVQSLKWMKAH--QNIFYDLNKVLCVESKIP-IVQLYEW--RKFENCQTDWLVFSVC
LLFFTIVSLTIIASVKRYRVHLEYVILRLKNRWKGV-QKSNEDMFLYDVYISYNDADCSW
VIETLYPKLE---NIKTWFGDKDSIPGRWKSEEIVGCINESRKVMFIMSESFLERGWHSY
AVQMAITHAFHNQRRSIVLIIKDGLPLDRLPNEIKNIWWCIEHFRWPENEQHDEMIFSTL
SKILKPK--

**>Cgi-TLRβ4**
LFK-APNITNIELFDNQIS---GSTLKTLLWNLIKLQKLNLQGCGINYLARGTFDRMPDL
RTIILKGNSLYGWDPTMFNKLFNLRALYLSGNSVAVVNRTSLIGKINLKFIDLADNPFAC
TCQQLWFRDWLKTAKNITVAFYPKRYVCRSPPKWDNTLVALFNYTEED-
CREPWILIGSV
LGSVVFVCMVVVIVIYTHLPTVRNIIYLIRLRRKGYVRLVNSEEYMFDCYVVSCETDEQW
VFQTLSSTLEVKHSYRLCIPTRDFDIGASIADQIEEKMRECKKIIIVMSNDFAQDEWCQF
QLEKAQERIRNQGEEAVVSIMLHDIDHKHMTSTIKNLLRKSSYATWVKGKIVSKLFWDIV
VAAIEK-PP
**>Cgi-TLRβ3**
LFNNTRNLRVLDMTGVQFSHNLEEKMFQLFKPLTGLEELTLKKTSLSTFPVSVFQFMP
NL
TKLSLQDCYFNQSYLRKLSAPASLKVILLDNNLITSINETNI---NNIDQMSLKSNPFLC
TCDLVFFRKWIETNS-KRLLGWPNDYTCNLPQEWKGKNLADFHLSYLS-CHPPYIIMAIS
ISFAVLAIATVSCIIYKKRWHIKYYLYLLRAKKRGYEVL-GGDDFAYDVFVAYNSDDRIW
VISEMIPRLENEEHLKLCLHDRDFQVGKLIVDNITDAMHRSRKILIILSNSFAQSHWCRF
ETMMAQLRSINHGENTVVVVILENILTKNMNNSLHMLLKSTTFIEWTNERAAKEMFWTR
LVSSIKT---
**>Cgi-TLRβ1**
SFNHLQSLKHLKFDDNNLGGLISDNIGTLFAGLHKLETLSLSKNFLHNLPISIFKDLSSL
QTLTMKSNRISGWNNGLFKQTSALRSLDLSDNSISLVNSSSLADLSNFQMLNLSNNPL
ACTCDLRWFRDWVNQT--RVNIANVGNYVCNSPNVWKGKPFLSFDRTKIN-CV-
LYFVVGVS
IA-SGLAVLVFCVIIYKKRWWILYRCYRLKNCC-RYQPIQDGQELVFDAYISYADDDYKW
VLEQLLPDIDSKGEFKLYFHDRDSVPGSSMISSISDNIEMSRKVIIVLTEKYLSSARHKF
EIDLAVMLKSQGVIDDIIVINVCGVSFACIPKSLQRKVSKDEFLLWKDDVDAIWLFKQRL
KAELKR---
**>Cgi-TLRβ2**
IFRYCRGLNVLDMTRIRLTTYDGVMLYELLHHLTNLTKLVLQSTLVLTLPENLFSRMPFL
GSLHLDHCYLSQWKLGVFRNVASVKTLYLDHNEIAIINQTSFELLRGLKQLSLGYNPYL
CTCDMVWFREWTGNNT-KVMLNWPYAYKCKSPREWATKLFSDFSLSYSY-
CHPPYVIAAIS
TAAGVVLIVIVVGLFYHYRWHIKYFFYLMRARKRGYEPLPGDDDFIYDVFVAYHSDDRV
WVISELIPCLERKEKLRLCLHDRDFEVGKLIVDNITEKINSSRKVLLLLSNNFIQNRWCKF
EMAMVHARNVEEDRSDIVVVILENIRTQNMSNSLHVLLKTTNFLEWSNKKSAKELFWK
RLVASVK-PES
**>Cgi-TLRγ1**
FFIPFVGLEILNLSNNALSQMFSDENGDFFQSQRRLTDLDLSLNRIAHLPGHVFQHNSKI
SRLNLSFNSLSDFNV--INHMKHLSQLDLSHNQLTQLSKNVRASLDAIAKVYLLGNNLKC
ICGTLDFLKWLRDSK-SIYFVGINNYTCLFENA--AASFNEIIVQVLEKCSSTLIIVLMT
TLIIVTMTTTVSRILYRYRWKLRYMYYVAKEKYKTHSEEKDRSSFRFDAFISYAEEERLF
VFK-LVKYLEEKCNLRLCIHHRDFIPGTGIADNITNAIHCSRHTVCFMTSHFLQSHWCMF
ELNMARMEAIYARQNVLFLVALEK-
TMKHLPLQLMDLVDSNSYLEYPGEESGIEAFRTKLGETLAS-SD
**>Obi-TLRβ5**
IFKSCPQLTNLILKNVSN------YPNDLLKPLTKLENLMITDGQVSKVPD--ICNMNNL

TELSIFYTTVSKWNNANCSVMRVLRKLVLKDNKIIYVNPKLFSHLSNL-HIDLSRNPFVC
DCKALWFRDWSRKN--AGRLKNYRNYRCFSPNSLGHIHLRNFSLSWDY-
CENSIAIAGVS
VGVLAVMFVFLAILSYTKRWSIRFCIYQSLVRKRKYKALVNNGRYKYDAFICYCSTDVS
WVLNKLLPIIEEENHFNLCLHDRDFLVGNDIVDNIVDSMQQSRKVVLVLSNDFAQSSWC
QFEASIAQQKILEEHYDIIIPVLLNEIPSNLQTKRLGVLLKQKTYLEWPNDEQYEGMFWE
RFIGRLNANNE
**>Obi-TLRα2**
-LTQL-GLP-IDLSGNKLNYLYNSIVNKTFIKFTNLKKLYLQNNLIEALQQKSFEGLKNL
LELILYNNKIQYIPENTFSETPKLKYLDLRNNKLQTITSEMF---KALQKIYLSDNPWSC
ECNDISFEKIFAKDTE--LLVNGEQIFCRKYDA----NIFNYGQE---FCQNVLISSVSA
VSSVIILIFLITFLVYAYRQEIQLLLFHFGYRFK---LIIDEENKLYDAFVSFDNSLDLF
VLNELLPQLEQNPPFKLCVHFRDFEVGLQITENIINSIENSKRTILLITDNFLKSEWCKY
EFQTAHYDGLSQKMNTLIVVLFENINEELLDPDLKLYLKTKTYLKYDD------WFWNKL
RFALPAKKD
**>Obi-TLRβ4**
IFKSCPQLTNLILKNVSN------YPNDLLKPLTKLENLMITDGQFSKVPD--ICNMNNL
TELSICYTNVRKWNKPNCSVMRVLQKLELKHNKIRYVNQELFSHLSNL-
NIDLSNNPFVC
DCKALWFRDWSREN--AGRLKNYRNYRCFIPDTFHHISLENFSLNWDY-CENFIAIFGGI
IAVLVVIFVFFAILSYEKQWSIGFCLYHSLVRKRKYKTLVNEVQYKHDALVCYCSADVN
WVVNKLLPIIEEENHFSLCLHERDVVVGNDTVDNIVDSMQQSRKVVLVLSNDFSQSSW
CQFEASIAQQKILKDHYDIIIPVLLNEIPSNLRTESLVDLMNQKTFLKWPNESKYE-------
---------
**>Obi-TLRβ2**
LFR-FPSISLQLIVTTLLSD--SKSINQSLAFLPNLVTLQMSFSNLKTIP-KVICNMVNL
TSLNLKGNAIVWWNDTNCFVMKKLHFLSFSENRISIVGDKTFLLINNL-RWDLSLNPFLC
NCENAWFKSWVEKNHQKFLYYPK-DFTCDTPADLRGKQLSDIDLGNNI-CGVVGITIGIV
LGSLMFVFVVVASISYYKRWALRYICYLLKSRKKQERSQQDEKSYVYDAFICYHNSDSK
YLLEKLQPKLEEENNFRLCIHDRDFVPGWDIVDNIVESIEKSHKIVLLLSNNFALSEWCQ
FESTMAQQRLFNEKKNTLIPILLEPIKIKNQTSRLTILLKEKTYLEWTDDKNGQKLFWARL
LNTMRGP--
**>Obi-TLRγ**
FFTFFTGLRLLNIENNAIGQVAEEGFSKAFLNLTNLEELYLSNNHIRYLSNNSLLQLKNL
RILNLHINLLESFDV--ISHMLNLSYLDLSKNILQELSENTFNAIEKISTVNIQSNNLKC
GCAQIRFLTWLHKVRNHLKI---MYSKCTHPNGTV--KLTDLIITYLNSCS-SFIIITVA
CLIVLLGCFSFGALVYHFRWKLRYLYYMIRERY-AYQRI-QTGEYLYDAFVSYAEEDRGC
VFEYLIPELEEKDTFKLNIHHRDFPAGKQIAENILSAIQSSRKCLILLSRSFLSSEWCMF
EYNMAKMECVHAERDLVIVIMLEELSVDILPLQLQHQIKMQSYICFPTTNPTSDVFWNN
LKKSIQE---
**>Obi-TLRβ1**
FSKELTRIRKLYLDSNKLGKFLTDKKGFLLSG---------------QLSKKIFQNNKHL
KYVYFRGNKITGWENNTFVTNLTLMELDVSNNFIFTFDSDSLKYINRKKKFNVTGNPFA
C
DCNLRWFRDWLNTT--TVDIVDKNGLTCNSPPDWQDKQLLDFTRSKID-
CTDLYYILGGV

GG-GFLLTVVIVLFAYTKRWYIRFKIFKLYQYVQEYEAI-PGDDMYFDGYISYSDKDADW
VEKYLMHTFDNNGNFKLCFRNRDFAYGKYIIGMIESSLAVSKKMIMVLTPYYKKDKRCE
FELQLGIMKLNI---KNVMPIVLKNLQPNQIPNSLKEIFETNKFIEWDNN-----------
---------
**>Obi-TLRβ3**
IFNACRHITLLKFQAVSI----NASINELLMDLKNLEYLELTGNTMRNVPD--VCDMKRL
HTLILSRTWIRKWQTTNCTVMNILRVFSLSYNRIFNVNKTSFALFSNL-KWDLSHNSYIC
NCRILWFRDWMRQN--SARLHYPKSYLCSNPAPVRFLQIAKYYVSWDY-
CANPIAVSGCV
LGTLSVIFIITGLVSYIKRWSIRYWVYLFFARRRKYQLL-ETSEHNYDAFVCYCGSDVGW
VTKYLLPILEEENDLHLCLHDRDFAVGNDIVDNIVDSIQQSRKVVLVLSSDFAQSQWCQ
FETSLAQQRLFEEKKDIIVPILLEEIPTELQTMRLALLLKQKTYLEWSNETRGQMLFWER
LVEILLETKE
**>Obi-TLRα3**
-ITVFPTMP-VWLQFNNITKLV-------------------------------PYLSQI
THLNLTKNSITSLNYTVMRNMVNLKQMILDWNLLTTLPKGIQNVQ--FEVLSINHNHFFC
DCTNIWLKKWLQKSRE--SILNWRRIVCNTV--DKVLDIVVVP--NDKICNPKTLTLGLS
LALAILILLVCLFLLIHYWLEIKVILYYLNIHPSLGSANTLNEQKKYDIFISYPDQTYQF
ATGPLLSTLQS-RGYSICLPDRDFVVGSAKEENILRAIKSSVRTLIVITKSHVEDEWQLF
TLRTAVQCSLKKPFNYLLCILDG-VDKSKLDLETQAYVTSHVVIDKDD------LLWKKL
FRSIPPART
**>Obi-TLRα1**
-LSD-SMLP-IYLSGNRLISLSRSNINGTFMTLINLKQLYMHDNDLTILTKETFQGLENL
EVITLNSNSISYIAPGMFAPMPKLKIVDVSSNRLHILDNSFL---KYLESIAIHNNPWIC
KCPFVMLQELYINKPD--LVVLSESVICDHEDV--AYPLFEF---DVQHCL-KVICALAI
FSAVFLTLIIAVISIACYREELKVWLFQYGWRIP---AKLDDSNRRYDVFVAYTSKNAMF
VEHELTPRLEREPPYQVCLTYRDYDVDISYAQNTINCIQNSKRTIMLVSNDFFQTEWFR
YDFQINNHDILKTLSERLIVILMEKVDRKKLECDLMFYAKTKKFLKYQD------HFWDKL
YYMLPKVRG
**>Bgl-TLRγ10**
-----------------------------FENLSDLKKLHLI------------------
--------QMRDI-------------------------------------TVNGATLSV
----------IKI---SLSGDHAGDST--------------------------------
------------------SLALPSSLFAFT---------QD---YKIDVFLGYSDTDYRF
PCQDLRAYLEDTLKLTTFLNDRDLLATLNKASGIVEAINSSWRVLLVCSEGFLKDEWSL
F
TMRSAMYAQSPANPGRVV-VMVHQRCLRLLPTELLSAVEEDNILV----------SEWK--
---------
**>Bgl-TLRγ3**
FFDSFPEVEVLVLDHCELDSEFSQHSYSLFRNLVKLKQLDLSYNALDILLPNTFSANLNL
RSLNLAFNRFRTIP-FDLSQTLGMNKLDMRQNSLETLSSKDMALLDELQKLLISGNVLSC
GCEHIQFLQWLHLT--DVRLDENRNYTCINNQG--LSSTSAYNIEVLWECWGYFNIALGM
FAFVNIGFVFVFMLTKNKTLIISGVLQLFT-EFKKRP--V---DYQYSVFIGYSDFDYQF
ACLTLRKFIEDDLKLSTFVGDRDLLPSIAMAEGIMAAMDSSWRIVLVVNKSFVNNNWFL
F
MVRSAVFSVSPANPLRVV-ILVEECCLPRLPSELLSSVPEDNVFVV--------TEWK--

---------
**>Bgl-TLRα2**
-LTDVPKIP-LRLDGNNLPSLRNSTVNNTFKGMKSVRSLFLNNNLLTIISPGVFSGLENL
ERIFLQNNFISLIDPQAL--LPYLYLINLRENDLNTLPIDGLGFVRELKRFSLSQNPYSC
QLDFVCFVLFIRDSAD--CIEDISDIKCSSNSL--GFTLLDFQIE---LCSE--TYALIA
ACVVIAFGLALLIVAYMNRDFLQVLCFRFGLRVM---KATEDNDRPYDAFISYSSKDEDF
VIHQLAPRLENDKKFQLCVHYRDFPVGACIAETIVRSVEASKRTILVVSDNFLDSEWCR
FEFQTAHQQVLNERRNRVILILMHDLDTEKLDSTLKVYMRTRTYLKYDD------WFWEKL
MFAMPDVQH
**>Bgl-TLRγ23**
FFDTLSTLKKLNISNNLLGSFLVLVSPRIFSSLRNLTVLDLSENFIIDFTCDLFSNLTSL
EYFNISKNALIRFEV--ISRMSNLIFLDFHLTRMTGLTSEFRDSIDRLSSIDMSNAPISC
NCKNYDFMTWMTSS--KAFSQGFKNYICVYPDQTGHV-
VNDFDMNLLNQCASVLLFSMIA
IAMIVVVGAVVGGIVYKYRWKLRYLYNAAYLQFKSSRRG-
EDDEFDYDAFISYDQEDGVF
VTQTLVPELEKR-EIHLCIHASEFTAGEYISSNIVKAVNRSRKTVVVLTQNMLSSYWCNF
EIQMANMEALHTGRRVLVFLLVDNIPTKDLGLELLYYIRSNTYIPFPKDFNGMSWLWDK
VANDIRND--
**>Bgl-TLRγ20**
FFKNLPTLTYLNLSINLLSRCHNVKKKYIYEALVNLEVLDLGLNNIDEFTPHILDHLISL
KKLFLDYNPLKSFDV--ISNMPQLEYLSLRHSRLHRLSVYTMKAIDEITSIDMAFNPILC
ECSNLDFIRWMTAS--SAFDPKFESYFCMYSDGSMQF-IDDFTLMILSECASVIIFFSVS
SGTTFLIILILIAILHRFRWKLKYMYYAAYLHYKSAR--DNGKAFSYDVFLCYHEDDESF
VLDTLCVELEKR-
GLKTLVHKRDFVSGKPIVSNIVEAVNCSRKTLVVLTDNMARSKWCQF
EVQMATMEAVSYKRPVLIFLLMSDVPCCIMGAELSYCVQNNTYLQYPSPSSEMDNFWI
KLVSDLKN---
**>Bgl-TLRγ19**
FFENFASLEFLDLSANTFGRKVRKGSKPIFSSLKNLRELNLRFVDLITVDKNVFEGLENL
EILHLQLNGIYYFEV--VSYLKKLQFVNFSFTELTGLRPQVTNFFDSIATLDFSETPIHC
YCANLEFISWLSRALQYIRFQRLKWFKCVYEDTTEKY-FHDFLHQFLGECTPVTLFFIVT
SATFLLVCIIIALVVYRFRWKLKYFYYSAYLYFKSYKRFHDDKDFEFDVFVSFANEDERF
VLKEILPELTTR-GLKVHIHTTNFRAGEYITTNIVNAVQCSRRTLIVVSSNLQKSQWCHF
ELQMANLESVHTGRPVMVFLLMESLPEDVLSREMLYHIQNNTYLQLPDEVRVMDIFWT
KLCSDLKD---
**>Bgl-TLRα3**
-LEKVPEIP-VYLDGNSLNKLTRSYLDGLFDNCTSLLHLRLDYNYLISISKSLFDKLIEL
RSLYLNDNLINFIAKEAFANLNSVEIITLDKNRLIMLD-------SSLKSLTLSGNPWQC
QCNTSTVLRVLHALND--IIVDRGNMCCYYVGTVQ--KLSELDRTPYELCVDTLLVCLVV
AMFVLLLIVVVLVIIFKGR-EVQAWVYNLGVRVK---DKTDAGNKYFDAFISYSNKDSEF
VSKVLVPALDE-KGYRLCVHYRDFPVGQNITDTIFRAIEESSRTIMLLSRHFVESEWCRF
EFQTAHYHILKEGSHRLVFILLDDLSDDELDPDLKVQLKSKTYLKFGD------WFWEKL
FFALPDVRK
**>Bgl-TLRγ22**
FFDSFTSIRELNLSNNLLGEFFQSNETLVFSKLKNLEILDLSSNGIHLLHFDLFMDLPRL

HHLNLAFNTLTTFGV--ITKLSKLMYLDLTKTGISKIPETARTFIDGL-SVFMGKCSISC
ECDNLDFLLWMVNS--KAFDKTFKNYMCFYMDSSS--PITDYTIEILRKCTSEMLFFMVG
CGTLFLFFLLLFGIIYRFRWKLRYLYYAAYLHYKKSGGE-GGAKFKYDAFVSYDHADEET
IVIHVCNELEARGLKLCVHGRDFRAGDYIASNVVKAVCSSRKTLVVLTKNLMNSYWCK
YELQMANMEAVHTGRQVLIFLLVENIPQGELGVELLYNIRNNTYIPYPTEPAFWDALWN
KLANDIRD---
**>Bgl-TLRγ21**
FFSELSSLLHLNLSSNLLGSFFRYESETVFYPLTNLMTLNLSFNDISELRPNIFANLINL
RQLQLQKNNLQKFDV--ITSLIKLVRLNLKLNRLSTMSSNITDHIDTLKVVDLSFSPISC
QCNNLAFINWMVNS--KAFHPNFINYQCVDSNTIQ--NITDYTVEKLNECSSVTIFLISS
GFSFVILCFVIGSVIYRFRWRIRYLYYAAYLYYSKTNSG-RDSDYKYDAFISYDQNDWKF
VVNKLMPEMEKR-RLKVCIHSKDFVAGDYIASNIVKAICSSSRTVVVLTRNMIKSYWCGY
EIQMANMEAVHTNRKVLLFLMMEDIPSSELSVDLLYNIRNNTYLQYNQDGVHMSRLWD
KLAYDIKH---
**>Bgl-TLRγ18**
FFNNLTSLKHLSLFQNLLGDCLNDKNGLIFSQLTELKVLNLSFNNLYYLGWEVFQGQAD
IEVIDLSVNRLDHITF--VSHMRKLRHLDLHKNDIETLPTGLTDHISSLKTLDMRQNPISC
GCENLDFLQWVVNT--
RVFGSDLYLYYCKFPDSDRAVRVPGYVVKRLVSCSSAVLYTVVS
CVTVLIMLILLAAVIYRFRWTLRYWYHAAKLKISSNQQM-DSDQFKYDVFVSYASKDIDF
VVKELCPRLKER-
NITVYVHGEKFKVGCYIADNIYTGIRKCRKTLVVVTQNMLASRWCNY
ELQIAREQARNTGRNVLVFLFLEELPTSRMGMGVLTHIKSSTYIMYPKLPQHRGAFWD
KLADDLRSS--
**>Bgl-TLRα4**
-FLEMPIIPNLYLDHNPLQSLN--------------------------------PYLSRL
SEIYIDNCLLTTVMPSAIAALKNIRVMTLHNNLLQKLPTSTRNITEKATNITLHNNRWAC
SCESLWLPRWISRHKA--VLWKPGNILCDYFQK-----LEDVS--EADNCK-SAMDNFLT
VILFVLSTVATVILFFCYNTDICAIVYKLGIEFR----LYGDQYCPFDILISYGQDNYKW
VVDTLVPYLEKPGGYRVCLNHREFPSSDCVLETLPTAVRLSRSAILVLSKEFLQKEWC
ML
EVRVAIQRLLLVGS-KLLIICMDKVNVDELSPELRAYIHTHHYLRYDE------DFWVKL
DLFLPRKLI
**>Bgl-TLRα1**
----IK-LRHLDESVAKL--------VTLFDALYVLEHMNFSTIGITTFPREWRRFFPKL
TYIDLSNNFISQVQFQNFPS-KTVVTFNLQRNNITVINMDVLNSWEKL-EVDIRNNPIHC
GCELESFLPHLQDTTTLAPYEYVKEMECSTPDALKGRKLYSL---HSSPCP-VYQVALIA
LGVTLSFLLVLVIILVRYKFEIRILLYRLHVRL-PCDADE-RHSKTYDAFISYSNDDDSW
VFENLVKFLENEKPFRLCIHQRDFVPGKTIFDNIVDSIEASRHTIIVLSPSFMKSHWAME
ELRQAYRQSLVEKTRHLVVLLLHKV---NL--N---------YCSF------------------------
**>Bgl-TLRγ16**
FFMNFSSLDQLFLGHNTLGDFLSHYNITPFIYLKQLTRLDLSYNGLTKVYRNLLSGLNAL
QELHMEENIMWDFNI--IDHMSNLRLIDLSHNQIKELPIHVREHIDNLKLIDLSFNPIRC
ECQYLYMILWMVSS--RAFNPAFENYMCVYPDGSYKI-IDDYTLQYLNACADYSVLLVVI
FSTLTMIILVIAGILYRFRWHLRYLYYAAYLKVKEGHHNQETRSYVYDVFVSYAHQDETF
VVQRLMPELSNR-GLNVFVHGRDFVVGHYIASNILTAIRESRKTLVVLTKNLINSTWCNY

ELQMANMESVHTGRQVLVFLIKDSLDITDLKTDLLYHIKNNTYIDYPHGPLALNLFWDKL
SLDLKN---
**>Bgl-TLRγ14**
MFQYLDELQELRLRNNQLNNFIK-TRKPVFQYLKQLKILDLTNNALTVVQSSIFEELGSL
EIIDLSRNNMRHFNL--LTNMSSLNFLNLSHTQLSSLSVETRQNIDLLTRVDMSRNPVRC
ECDNIDFLKWMVSS--RAFDVNLTDYMCQYKDTST-IVIKDYTLVYLARCADSTLFLVVL
SVTLCMVSFVVAAVVYRFRWRLRYMYYAAYLVVKGKRKDNEAELFRYDVFISYASEDE
EFILGKLLPEFDSR-
DLRVLVHGRDFAVGEFIASNIVTAVKESRKTLVVLTRNLLNSTWCNFELQMANMESIHT
GRPVLLFLIKESIPTTELTSDLLYHLNKNTYIVYPQ--EITDVFWDKLARDLLQ---
**>Bgl-TLRγ8**
FFDGFFGLEKLFLSKCELQRDFALHSSRVFQNLSNLQSLDLSYNYLNDLSQGTLYYNP
KLVWLNLSDNQFNRIP-
FDLKDTPNLLELDVRNNAISTVSKSITTELDQLANFWLSGNILSCGCQDLNFLHWLSST-
-MVTLDQGGNFTCMDRNG-ERSYTMRYHVDTLWECWGFLYLAIII
LCFYVTGVFLLVLVQRNKTFLVSFFLQLLG-NFKLKR--G---DYPIDVFVGYSDEDYHF
PCRDLRLYLEDVIKLKTFLNDRDLLASLSKASGIVDAINSSYRILLVCSESFLKDDWSLF
TMRAAMYAQSPANPSRVV-VVVHESCLHLLPTELLSVVNEENILVV--------SGWK--
---------
**>Bgl-TLRγ7**
FFDELTGLEYLALSKAGLNRDFSSFSRRLFQNLSNLTRLDLSINYLNALSKGTFSPNSKL
QWLDLSGNQFKDIP-FDLQYTPNLLELDVSSNALTTIDDDIARDLDHLVHLSLGGNILSC
SCSDLRFLQWLNLT--SVTFDHSRNYTCLNKDG-EKAYTLFYDLDSLWECWGFLYVAVII
VCLYVIGFFVILLLLRNKHFLVSYFLKILG-NIKLKR--T---DYPIHVYIAYSDIEYKF
SCSDLREYIEGTLKLNTFLNDRDLISSLNSAADIVKAMNSSWKILLVCSASFLNGDWAM
LTLRSAIYAQSPTNPARIM-VLVHQNDLLLLPHDLLSVVDDENMLII--------SEWK--
---------
**>Bgl-TLRγ6**
-------MENLALSNCRLERDFSQHSHVLFKNLTRLRQLDLSSNSLNYLSKNTFLFNSHL
QFVNLSRNLFREIP-FTLRYTPELRALDLSVNSLSSIDVSTTKDLDHLVKLYLQGNVLSC
GCNDITFLQWMKTT--LVTFDLNGNFTCINEKG-ERTYILFHDLESLWECNGFLYLSVII
MCLYFIGLCIVFIIYRNKQFLISYLLQTFV-GFKSTR--K---DYKIDVYIGYSDRDYKF
PCKDLREFFENSLGYKTFLIDRDLIASVDKASGIVDALNDSWRILLVCSESFLKEDWSMF
TMRSAIYIQSPANPARVV-VLVHKDCLHLLPTTLIGSVNEEKIIVV--------SEWK--
---------
**>Bgl-TLRγ1**
FFDDFSSLRYLILQSMMNEDFFRVSIDRIIQNMPELRYLDLTDNKLNFLPPNLFSRNSHI
THVILAKNRFSSFP-ITMDLVPNLKTLDLSGNAIIYLTEEETSSLTKHSYLLLAENNIAC
VCSQIKFLLWLNI---TF-LDNKGAYSCTSQDG-QLILTVLWDVLGFYQCYGYFMISIVL
LLVMSFIFLMAYLVHRFRTAIEAYLVRIFIKAVRMKS--SD---YKTHVFIGYADEDVGF
VRHILLRYLEEDLKVSTFVHHRDLGPGYTDQQ-MFESISDSWRILLVITQRYLNNYLSDI
IMKYASHSMSPANEKRLV-LLVQESQLYNIPGYLYDVLEDSRIIV---------SDLSA-
---------
**>Bgl-TLRγ5**
YLDTLPALENLALANCQLDREFSIHSGRLFQNLTRLQQLDLSSNLLNYLSTDTFMYNKH
LKWLTLAQNQFREIP-FSLKYTPELEVLDLRQNSLNTIDMASIHQLENIVKLLLSGNDLSC

GCNDLQFLQWMRST--AVTFDQDGNFTCTNKDG-KTTYTLAYDIEYLWECTGYFYIVLIV
FCLYLIGCSIVFIMMKNKHFITVYILKRIF-GIEHTR--R---DYPIDVYIAYSDTDYQF
PCNELRQFIEQSLGMTTFLIDRDLNASFDLALGIVNAINKSWRVLLVCSESFLREGWSM
FTFSSAIYAQSPANPARIV-ALVHRDCLPLLPMELFGCINEDNILYV--------SEWA--
---------

**>Bgl-TLRγ13**

FFPQ-SSLISLNISNNILGEYFALGRKKIFLGLGYLRFLDISMNLIYKLPRDFLSGLKSL
EVLLATKNRLQALNV--LSQMSSVWFMNFSQNSITWIDKVTRDDLDLLASLDISFNPLPC
TCDGIEILNWLAFT--NVRLVNQMYMKCQTSTG-ETVSLGDLRAQQVQACASAIILVISI
SSTVVVTLMVSLATLYRFRWKLRYLRNIALTKY-GFRPKKTGKKFQHDAYILYEDQTIKF
VFRDFIQELEVKRGHRLLLVDRDIMPGTIMTTAILSAVQNSYKTIPVVTPYFFDVWYSEY
AVQMAIMEEHYEPRQILHLCLYQATDPKDMPKDLLSVMKRNRYTEFPPETEMVKQFW
DQLSSTIQQE--

**>Bgl-TLRγ2**

FFDSYPALELLALESCRIDGLLSQHSFRVFQNLHSLQSLDLSFNSLDMLSPQTFSTNPN
LTSLNLAGNRFRNVP-FDIKLTPNVKFLDIRQNALTTIDISSRKALDELNRLLLSGNILSC
GCENLLLLQWLQET--RVELDGNRNFTCMNIKG--
LSSTLAYNLDGLWECWGFFNLSMAL
LCFTLLAYILFFTWIKNKTVILSSILQIFT-DFKKKP--S---DYQSGVYLGYAESEYKF
PCSELRQYIEDELCLNTFIRDRDLLPSLDIAQGVMDAINSSWRILLVINERFLHQDWFLF
TIRAAIYSISPANPSRVV-VLVEKNKVHSVPTELLSSVPNENIIVVSQ------------
-----Q---

**>Bgl-TLRγ9**

FFDGFSGLETLALSKCLIQRDFAFHSHRLFQNLKELRQLGLSFNSLNAFSNATFSFNSN
L
QFLNLSDNQFNYLP-FNLKHTPELRVFDVTNNSIITINVDARHELDRLARLFLRGNILSC
GCSDLLFLQWLKNT--LVELDQGGNFSCIDKDG-ERSYTLCHDLESLWPCWGFLSIAVII
VCLYVIVFFIVFLYIKRKTFIITYFLQLLG-HFHRSR--Q---DYKIDVFLGYSDTDYRF
PCQDLRAYLEDTLKLTTFLNDRDLLATLNKASGIVEAINSSWRVLLVCSEGFLKDEWSL
F
TMRSAMYAQSPANPGRVV-VMVHQRCLRLLPTELLSAVEEDNILVV--------SEWK--
---------

**>Bgl-TLRγ15**

FFHNFPNLKKLMLGNNKLETYFNLPNYTLFSKLKKLKTLDLSDNAISKMPTDILAGLTSL
KVLYFEHNTLWTFNL--LSHMMNLRYVYLRHSQVNSLSEDVRQHIDSIGRFDLSFNPIHC
DCENYDFLKWMMNS--RAFDPKFTNYMCQYPDSSYK-NITDYTLRILRKCTDSFIFLFVL
AATFVMIAFVLAGIIYRFRWKLRYIYYATYLRLKSVDEE-NSEQFRYDVFISYAHQDEEF
ILKVLYPELGSR-GLNVHVHGRDFVAGEFIASNIVTAVRESRKTLVVLTLDLLKSKWCNY
EIQMANMESVHTGRQVLVFLLKDSLNNKQLGTELLFHIRNNTYIVYPQNDEELAVFWDK
LYKDLRK---

**>Bgl-TLRγ17**

FFSCLNSLRNLTLSVNMLGDFIGSSKERLFENLSSLSYLDLSFNSIDKMQVYFFHGLSN
VTEIDLSRNKISEFNV--ITKMNQLRRLNLSDNKISRLFSNVTDQIDRIKQVDLSKNPIDC
TCANLEFLKWMVNW---VNVSQSQGYLCKQDDGSI--AMPDYTVLSLNQCASVVIFLIII

GATLVLACVIVGMIIYRFRWSLRYWYHVAYLNYQQKRKSDRRQKFEYDVFISYVHNDE
TFVAQTLSTELEKR-
HVKVYMHGQKFVAGNYIASNIVQAVKSCRKTLVVLTNKYVRSQWCYY
EVQMANMEAISAGRPVLVFLIKEKIPNHKLG-EILTFIKTNTYIPYPQEDRELKIFYDKL
ASDLL----
**>Bgl-TLRγ11**
---------------------------------------------------------------------------
HITRLQLLDFSRNSITWITESTRDDLDALAELDLTFNPLPC
TCSGIEFIKWLATT--KVKLIDQVNLRCRLKDG-GSTSVGDLMLLFLQSCISSWILSVSI
LSAVFMAVVLGLVLMYRYRWKLRYLRNVAIAKF-
GFEPKKHQGLFKYDAFLVYDSDDMQFVLNECVQELEVRRGIKLCIGDRDFMPGTYVAS
DIVSAVQNSYRTVLLVTPEFYDDDYVEYAVNMAINEEIHTSRQVLYLCLYQPVALAEMP
RDLVAILKRNEFIEYPPEEGLIENFWDQLTAAVRQE--
**>Bgl-TLRγ12**
FFRP-NSLISLNISNNILGESFALDSGKVFSRLGYLRFLDISMNLLYRLPRGFLSGLKSL
EVLLATNNKLQALNL--LSHMSSVWLMNFSQNSITWIDKVTRDDLDFLASLDISFNPLPC
TCDGVEVLNWMAFT--NVRLVNQMYLKCQTNTG-EIVSFGDLRAEQVQACASAIVLVISI
SSAVVVTLMVTLALVYRFRWKLRYLRNIALAKY-GFKPKKTGKKFQHDAYILYEDQTINF
VFNDFIQELEVKRGHRLLLVDRDIMPGTYMTTAILSAVQNSYKTIPVVSPYFFDGLYSEY
AVKMAVMEEIYEPRPVLHLCLYQPTDHEGMSKDLLSIMQRNHYTEFPPDPELVKQFW
DQLSNVIQQD--
**>Bgl-TLRγ4**
FLDELYGLENLALSKCQFDRNFALKSARILQNITKLKVLDISNNSLNGLSKGTFSRNSEL
LYLSLSGNQFKDIP-FDLKFTPNLKILDLSSNIITTLTTDTTDALDLLNQLMLNGNILSC
GCHDLSFLQWLNST--LVSFDNNRNYTCMNKDG-
ERTNTLTFDLESLWQCWGFFYVAMITLCLYVTGAVLIFLMLKNKNFLVSYFLQIFG-
NFKHTR--S---DYKTDVYIGYSDEDYRF
PCIELREHLERNLKLSTFIIDRDLLASLDKASGIVDAINSCWRVLLVCSKSFLKDEWSIF
TMRSAMYAQSPANPAKIV-LMVHTSCLSLLPADLLSVVNDENILVV---------SEWK--
---------
**>Ttr-TLRγ4**
FMSSFPNLKYLSLAYNNLGHMFDDVSKCVFLSLTELQTLDLSHNQIAKLPVDLFLNQHN
LKELILNHNKLKTMG---VASMASLQYMDLSYNEIRD--KSML---ESIATINLTKNAVSC
TCLNVVFLTWLINT--HINISGKETVYCLQQQK----MLIHFNFN---DCKKDYVIIGSV
VG-LVLVLIVIAVIVSNDR--VKYHIYLLKYKLR---NISRN-TEEDRIFISYCSEDRIW
VLRKLKPELEA-MGYKLFIHELDFEVGNFIADNIVHAIDTCFKTILVLSDNFVSSGWCMF
ELKMTLAK----S-DCAIPIYYKPVTKKNNTLLLKYLNKVKTYMKWPEDDREQYYFWQRL
KHALDKQED
**>Ttr-TLRα5**
-LTVMPQAP-LLLDNNNIEHLE--------------------------------YYLNDV
TKLILRHNAIADVPPNFVKLVTDMTLLDLSYNRIRYIDDDVLSSLKPTLSIAINHNPLAC
DCHSHSLKQWVSDHRK--RIVNLADITCFGG--AGGISILEAS--DLSICLD---IILPS
VIVPVVFICILILLVYIFRNEIKVILYKFNLHLN-----EEDETAVHDAFISYCSTDENW
VIKELANKLEMN--YKVCHHQKNFEPGVAIADNIVKSIDQSRRTILVLSNDFLNSDWCKY
EFQAAHYRALKNRQKYLIIVMLHKIDVSKLDNTLRLYVKTNGIIKVNE------LFWQKL
FYEMPIRTL

**>Ttr-TLRβ3**
LFEDLGLLKELVLKAADIANFRSKDIRAMLNVLIGLEHLNLEKVRLYSIPPTTFHHMHNL
SKLVLSDNFLSHLPEDLFFNLTNLKVLQLNHNRISQVSTKTFGFLDSLESIDLSGNPFAC
GCSLHWFLQWMNSTNVKVVGSQRFSYKCSSPPALRGKSLHEYYYKYRQNCLPIILVA
SVSGSCFLALLVSVICIVYRSRWYIRYLFYLLRARRKRQRKRNDEKDFAYDAFVCYNKD
DQDWVVRRLLPELEYNGEFKLCLHDRDFMPGIDIIDNIIESMEQSRRTILILSNSFAQSQ
WCQWELSMAQHKVLQDEGDILVLVLLEQIRSDNMSLKLHYLMRTKTYIEWTDNEDGR
KLFWEKLKGTLKAKPE
**>Ttr-TLRδ**
SLSGMEKLTTICLASNCLGSVP----PEIM-SARNLKQLDMSYNKISSIPP--IGGLKEL
RYLNMKSNRLRQLPN--LCQLKHLEIVCFSENTISDPNVDELLDMSKIKMLCLHSNRIPA
N---K-VQNLLKKASRDIRLEN-----C----------VEETHVKYLKKCL---------
-------------------------------------AMDENAVHKWDVLILHDDKDEEI
IENEIRPKLEEEMDFRVCIPYRDETMGMSKVAERSNLINFSKTIMLVITEKFNSSK--IL
GLDEVMLNGLDSETKCLIPVLWSKV---QVPKELKGRTMVRR----DS--VQEKYFWQKI
RKAIQSH--
**>Ttr-TLRα2**
-LNKIPKIS-IDLSGNNIPLIRHSHIDGSFTNMSNLLLLYLNNNNLKVLSRYTFEALPVL
EELYLHGNKLTFIEDETFLGLKKLRIISLKSNLIKTLPYTDF---SHLTSVSLAENPYDC
DCNFSRFKSWIFSSLA--TVIDSNDVFCVFYGLFPGSRLFNF---DLNYCELSSMIAIII
ILIVFVVIVALATVAYYYRNLIKVWLYNYGLRPR-----PDDSDKIYDAFVSYSSFDEST
VVHTLAPKLETNPKYKLCLHYRDFPIGSSIAETIVESVENSKRVIMLLSENYLSSEWCIY
EFKTAHHQVLKDRTNRLIVILYDEINMDNLDPDLRLYLKTNTYLCWKD------WFWQKL
YYAMPDVSD
**>Ttr-TLRα3**
-LTSLPFTEDLDLQNNSIRELT--------------------------------PYLKHV
KILNIANNKLEIVSAEAIQSLKTVQKFNLSGNRLTKL--NVNHFKTNLETLDIQDNQFTC
NCEDQWFQEWLLQINN--AVVNADSVRCHNK----DVAILSAS--HTDCGLANHTILTIC
VSVGAVMVCVAVVMVYIFRKEIKVLINHFSWHPR--RENDDNRHYLYDAFISYNLLNLDF
VRNSLIKNLE--PRYQLCIHNRDFLLGNEIADNIVTSINASKRFIAVVSKAFIESEWCQY
EFQFAHNDAMKDKRNNIIIILMEDSDLGEIDNCLKIYLRTHTYLSYKD------LFLQKL
LYSMPQVRT
**>Ttr-TLRγ3**
LMSSFPNILYLSLANNKLGELQEKQFKDVFYPLTKLEEINLSGNNITYFPVNVFLAQTKL
KRLLLHDNSLKVWH-INMSTMTSLEYLDLSENQITIIGQMSMNYFKTIVSINLNDNKIDC
LCFNLEMITWIQK-S--KFIHQRDNLKC--GDTK--KSILSYDPK---SCEVAELVIGVT
LGISALFIGVLFLIFYKIHW-LKYKLHLLKWRWRGMMA--DQNIEQEMIFISYENRDRCW
VINTLLPKLEG-MNYKTYIHNRDFTVGRPIADNIVHAIDICARTVLILSDHFAQSEWCVF
ELNMALV------KNSVVPIQYAP---------------------------------------------
**>Ttr-TLRα1**
-VEDVP----------------------------SLEELSLQFCNFSVITRNMLQNYPNL
KTMYMHNNRINYIETAALTRKVRINVITLDNNRLTHID-------KQVIIVRLQGNPWDC
QCHLKPLSDYVRNHIK-------GNITCYSPPS-----LASTPLQQVNTCEQGLLFLPIM
LGLLLALVLASTCCVYWYRYEIKIMWNKY-------RKAYKTEKHTYHAFVSHSSVDFKF
VKDNLLVSLE--PTYKLYVYYRDSIPGSTIVEDIVKAIDDSAITIILLSQNFLHSDWTKL
EFKQSYFKAMKSKSNNMIIILMEDIPLDSIKPQIKAYIRTKTYIHKND------RFFEKL

TSSMPKEEM
**>Ttr-TLRγ2**
-------------------------------------------NDISYFPDNIFIHQTKL
KKLILRRNAFQVWN-VNMSTMLSLRYLDLSKNLLTVIGETSLTFMDTFMTINLEDNLFIC
SCPYLPTISWIQEN--NKSIRQAQNLKCKMGEN--EIKLMSYNAE---SCHVIYDIIGIT
SSISVIIVMATIFISYKFHW-IKYKFHIIKWRLRNCFGIHDQPANNERIFISYENRDRRW
VLDTLVPKLENTMNYNTCIHAWDFMPGYPIADNIVRAIDICTKTIVVLSDHFAESNWCQL
ELQMALV------KHSVIPIRYAPIEKQNKTRLLKYLTKANVYIDWYDMHNKEDAFWDKL
KYTLDRDDE
**>Ttr-TLRγ1**
-------------------------------------------NDISYFPDNIFIHQTKL
KKLILRRNAFQVWN-VNMSTMLSLRYLDLSKNLLTVIGETSLTFMDTFMTINLEDNLFIC
SCPYLPTISWIQEN--NKSIRQAQNLKCKMGEN--EIKLMSYNAE---SCHVIYDIIGIT
SSISVIIVMATIFISYKFHW-IKYKFHIIKWRLRNCFGIHDQPANNERIFISYENRDRRW
VLDTLVPKLENTMNYNTCIHARDFMPGYLIADNIVRAIDICTKTIVVLSDHFAESNWCQF
ELQMALV------KDSVIPIRYAPIEKQNKTRLLKYLAKANVYIDRYNMHNKEDAF----
---------
**>Ttr-TLRβ5**
-----------------------------------------------------------LTHLIVTGNKLTTLSPELF---
THLKYLDVSNNSITSFSREIVGDLGYIERFIFDDNKIECDCELSHFQQWLLTTLI----
DTSKTERCYN---YEGVRIIDYQPTWID-CDNTYVVVGSI
GS-
FCLLVATVAALLVYYRWDVKYWFILRKIKAKRYHNMHDENNVMYDAFVSYSSLDEGW
IYNELIPNIEDDIKFQLLMDQRDFLPGHYIIENIVQGIDSSHKVLLIISLNFIESQWCTF
ETRFAEQSSIETG-QRLILIFLEPLKKSEMSRHLQRL-----------------------------S---
**>Ttr-TLRα4**
-LNALPYVP-LYLQDNHITHLT--------------------------------DYLALI
TELNLDHNAISEIPLAFLNSIPKMKTLKLAYNQIKYFPEEIEETR--AFNWSMHHNPIAC
NCYSLWLKKWVSANRK--RIDNLHDIVCFSG--AGGVAILEAS--DHLICID---IILAA
TITPASIIIIMVLLGCIFRKELKVILYKFNWHPK-----RENESLPFDAFVSYCSADEHW
IVTQLAKKLESNPPYKLCLHYKSFEPGVAIADNIVTSIDNSKRTILVLSDKFLESEWCRY
EFQAAHYRALKNRRKYLIIIMLNKIDPSKLDKNLRLYLKTNGYIKPTE------LFWEKL
KYELPMKSS
>Ttr-TLRβ2
IFMNLTQLQWLVANHNNIGMCLTKGMSKLFQNLKSLQWLDLSSNQIETLPKELFQNLK
SLKYLNLSSNRISYWASEQFTALKKLQTLDFNSNVITTINKSSIGQLENV-
HLNLSNNLFSCDCDLRWFRNYINYT--KIDFTYIKDYLCAAPPDFQGKHFLKFHSNMII-
CSPYLIRYISIGGAVVLIVLMISLATYNWRWYLKLKLFRLKNTLRGFQ--
EDDDIVTYDAYLSFAEEDRDWVTRTLLPKIDNEGRYRIYYDDRDDMPGDNIINAIDSGIE
KSEKSIVVFSKKYATNGRIDVDLTLI----
LDKPHQRVILIMLEEVPLRMIPRCLHSTLWSNQHLLWTEDVNGQALFWEKLNNKLMED
--
**>Ttr-TLRβ4**
------------MGDNSLYR--PNDLPALFAPLHSLTYLSISKNKLDYLHEDTFNGLYNL
EKLILTTNKLEYLSTDLFKNTTKLKTLYLAKNSLKTINAGTFEKLTFLKDINLGENQFDC
HCDIRPLRDWLKYKQKKKAIKIQGDLNCTTPPNLRNSLIVDYNPSWLD-CDNEYLLISSS

C--SMGFVLITITVIYIFHWNIKLFFAIRKANRKIDGENNPLLRKRYHAFISYANDSLWW
IKKHLLPNLQDNFEFNLCIRDRDFRAGQAEVDNIIDGMQNSTCTIFLITAEFIDSGWRQF
EMNVILRGLIDDPNNRFILVFLEDIPNNKLPIVLSTLKKNVDCLYWP--KVKRIQFWAKL
KVRILGK--
**>Ttr-TLRβ1**
IFFNISTLQILNLNKNLLSELP----DVLFTNLENLQCLDLSSNLLEVIPEKLFANLKSL
TDLNLANNMLYNTN-GIFHPLIHLTFLNLSSNSLTMITKDTLAGPKKLKTVDLNKNVFKC
TCDLQWFVDRLRQSKQCPYIVQLREYKCTN---LPGTCVANFMPSQWE-CHSIFIVVISV
LGS-ICITLLLMGCCYRYRFYLLHFCFVLKRLRETYEELYDNTQYRFDAFICYNDEDLNW
VQSQLLPKLRE-ATIKICINFMHFRIGAPRIDTIMEGIQTSRKTVLVISRHFLDDDWCLF
EMNVAAHRLFEEGKDNLVIIFLEPIQYSEMPLTLQAVVRTKRYLEWSTNEQGKDLFWET
LCYLLKTRPS
**>Hps-TLRγ4**
FMSSFPNLAHLSLAKNKL----KK---NVFWPLKLLEYLDLSDNQISMLPKGVFSQQSSL
KYLILKDNALTKLS-LGLKNMKCLKYVDVSVNKLETLEPQTRSFLEKKMFIFMEDNVFQC
SCSNIDMLYWMRNMNLKSSVQRWSQVKCHNYEN---VNLTDYDIS---KCDSIRTLVVTL
IGVLV-LMLVMGVVIYKSDR-LRYKWHLLKWRLRNNR---D-HRQNFKIFFSYGSRDRQW
VWEVLKPKLEQ-DGYSLFIHEIDFHVGECIADNIVYAIDVCDQIVFVLSDNFVSSEWCMF
ELNMALV------KHCIVPIRLSPIMKRN--RLIKYLTKTRTYLEW-KDKESADEFWARL
YSRLNRK--
**>Hps-TLRγ3**
YLSSLPNLAYLSLAKNKL----NI---DVFTPLKLLEYLDLSGNQIAILPKNVFSQQDNL
KYLIMKNNALKTLN-
FQLKNMNSLEYIDASENKLGTLGQQTRFFLEMMMSISLEDNVFQC
SCSNVDMINWMTKTSLKVSRQRWSQIECFNLRS---VNLTDYDIS---KCDSVTTIIATI
VGVLAPAMFCMGLVIYNYDR-IRYKWHLLKWRIRNYQAVVR-
HRERFQIFLSYDSCDRQWVWKVLKPKLER-
EGYSLFIHEIDFHVGECIADNIVYAIDVCDQIVFVLSDNFVSSEWCMFELNMALV------
KHCIVPIRLSPIMKRN--RLIKYLTKTRTYLEW-KDKQSADEFWARLYSRLNRK--
**>Hps-TLRδ**
DFSNLEKLRTVILLCNRLEKFP----TSLL-DVKSLAQLELANNRIREIPP--IGQLREI
KFLSVKCNRLTSLPE--LAKLEVAEVICFSENMIVDVPVESLLRFKNLKTLCLHSNRIQN
H---K-VMQLHE----DVRLEN-----C----------IGD-DIQ---KCR---------
----------F---AYHVR--------SSNFRL-KCRNM-DKTTWKWDVYIAYAEAAEHI
VDEELVPKLTN-MGLTACVYYKDSQPGKDIMADRRDMIDRSKKILVLLTKDTSYSDF-IS
EIQHIVSEGPKDQTARLIPVQWDE---AIIPDELKKVVVTSR----RT--AQEKVFWSRI
EKALKS---
**>Hps-TLRα**
-LTSLPQVP-LYLSNNRITELS-------------------------------SYLGSL
TKLHLDHNSLREINPNFLSQLKNLTFLSITWNNIKYFPESIKGT---LFNLSIHHNPIAC
DCHSLWLKKWISRSRN--RFDNLKDIVCVDG--AGGSPVLEAQ--DNQICL---KMILLG
TIVPFVVIIIIVMLVFIFRKELKVILYKFHWHPR-----REDYTLPYDAFVSYSSGDEHW
VVSQLTKKLEGSRPFKLCLHYKSFEPGVAIADNIVTSIDSSRRTILVLSNNFLNSEWCKY
EFQAAHYRALKNRKKYLIIMLNEVNTDKLDKNLKLYLKTNGYIKPSE------LFWEKL
QYEMPVLEP
**>Hps-TLRγ1**

ATSAFPDLTYFSLAKNKLEMMFNINSTDVFHPLEKLQFLDLSENGIIDVPSNVVEKQVSL
RMLNLSNNNMQTFKV--LFSLRNLTYLDISNNLLKTIDVISTIGLDQILRINMDKNDFEC
VCSNVRTIDWIRKDSLRSTILRRDDLQCKVIKEGGWNKIVDY---QLNDCDTDLIVLPIA
SA--LAVITFIFGIIFWKRNQIKYKLHLLKWRW-GFLKT-NPHVERGQIFISYDHRDGDW
VRNTLRPNIQE-MGYRPYLHEIDFVPGESIADNIVHAIDVCDKTVVIISDYYAESQWCQF
ELQMAITKGL----GYVIPIKYAKLKRK--NKLFQYFMKCVTYLEWPADDDTRAKFWIRL
GRAIAKE--

**>Hps-TLRγ2**

YLSSLPNLAHLSLAKNKL----NI--TDVFTPLKLLEYLDLSGNQIAILPKNVFSQQDNL
KYLIMKNNALKTLN-
FQLKNMNSLEYIDASENKLGTLGQQTRFFLEMMMSISLEDNVFQC
SCSNVDMINWMTKTSLKVSRQRWSQIECFNLRS---VNLTDYDIS---KCDSVTTIIATI
VGVLAPAMFCMGLVIYNYDR-IRYKWHLLKWRIRNYQAVVR-
HRERFQIFLSYDSCDRQWVWKVLKPKLER-
EGYSLFIHEIDFHVGECIADNIVHAIDTCDQIIIVLSDNFASSEWCMF
ELHMALV------KHCIVPIRLSPIVEHNN-RLITFLTKTRTYLEW-KNKPSGEIFWARL
YGTLNRE--

**>Lan-TLRδ7**

EVENF-ELRSLCLACNVIDNLP----GKFF--MKQLQELDVSYNRLTQIPA--IRNLKNL
EFLRLTGNRLQTIPS--IESLDRLLYLCLSENALVDIPTNALHRLINIKSLCLNSNRLPC
E---V-VIQVIQESSPKVSLREN----C------K--EIRD-------------------
--------------AFRAK---------------QYFEK-DGKDFESDVYVMHADEDYAL
VDQEIVPHLEER-NLKVTVNIQALRPGLPVSDQLVHFIESSRKILVVFTKNDVFEHTCLT
KVKAALEKRKKES-ETSPIVLVE-CPKSKVPHEFKDLYVIHR----RT--THEKHFWPNI
INAITQ---

>Lana17091

-LIHMSKID-LDVSNNYLETIP--------------------------------SD-TMY
REVYLDNNSISTFPTSQV--LPHLTTLSLRYNSIRTISMRQIEKDT-VNDLYLGGNPWRC
DCHARSIKHWLLNNSN--IIRDLDDITCVSGELTLGKSIKNVP--DNNGCPI---IAAIV
GGLVFVVIIACMLLLYKCNLKVRIWLYKFRFRFK--D--KQDSDKIYDAFISYSSLDEKY
VVQTLVPGLENTPPFKVCVHYKHFIPGASIAESIVEAVENSKRTIMLLSQNFIHSEWCTY
EFKTAHHQVLKDRSNHLIVVVLGDIP-SDLDSDLKLYLSTNTYLRADD------WFWEKL
LYAMPKLEN

**> Lan-TLRδ1**

DFKC-LKLKALYLNANYIRELT----PNIL-KLEELIIFDGSYNELQVLPN--IDQLQSL
KYIRLKQNQLRRLPE--LGNVKSLEVICVSENRLQDIPAEKLAKLPKL-RLCLHSNRLGQ
----K-VVRTLKEAKFEVRFDN-RSVD---P------KISKI---KVEGCH---------
------MTVLT-----------------------------RETPVFDVFMLYSEEDEKV
ITDEFLPKLEKKAELKVCFASRDYIPGHFELKEALTNMRKSRKIIALLTEHFDEQK--AV
EINHAVDADLARQSCSVIPVVWGNV---KMPVQFKRIVPLRR------------VDWDRL
ITAIKE---

**>Lan-TLRα2**

-ITTLPAMP-IYLQNNKLEIIT-------------------------------DYFSRV
HTLVASNNSIAKITSRIFWY---ISHVQLDGNNLKSLPQDIESMKHNITSLSLSRNPWTC
SCENLWLKSWLLKKRK--VI-HMDSVICTNE--VKGKPISQVT--EEMLCHPSYIQVAVS
LGVLLLLTLITIAVLYKYRFEVKVILHRFNWHPR-----QEMTEKLYDAFISYSSEDRLW

VHTTLAPTLENQLPYRLCMHCRDFLPGEAIDNNIIQAIQNSRCTILVLTKNFLRSNWCIF
EFQQAHYQMIHNAHFKVIVILKEDIPAEEMDDDLRAYLRTHTYLEAKD------WFWKKL
LYVMPTMNK

**>Lan-TLRβ6**

---------------------------MFRNLTQLRKLYIQNTGLSFLPPNVFVNNGMM
SELQLQSNFLSTWDPIVFQPLLSLRKLFMDHNNIRILNETSFFIWDNLTDLNLAGNPFSC
TCENLWFRNWIQST--NVKLLQLHAYLCYEPKKLSKSPFLDWHPTKAQ-CTPAWVIASAI
GVPTLLFLALVIVVSHRYRWYIRYWCFTLRSRYKRLEPFEDNGTYVFDAFVSYNCHDR
PWVIQRLLPKLEYDAGFKLCLHDRDFIVGHDIVDNIVDGIDVSRKTILVLSNNFAQSQWC
QLELTMAQHKLFDENKDILVLILLEDIKPENLSNRLTLLLRKQTYIEWPSEEEGQDLFWE
RVKAALQKPSG

**>Lan-TLRβ9**

IFNNVPTLTELCLDNNQFYRILLDILRDLLRPLRHLRCLSLTANKLTELPLGMFDGLANL
TTLDLNLNSLRSLPVEIFRHQRQMTDLHLDRNSIFTLSGHMFANLTALKNFNYARNKIIC
DCNIRSFQSWLATT--SVNV---PRELCFGPEWAQKTPIKEFRPSWFA-CDD--VYLAAI
SGGCVFFVIFLSAVLYSFRWDILYIYAIYRASGKKSKLGRREPHKTYDAIAMYSPTSVTW
IKKHLIPNLEEDIRFKLCINDRDYIVGDPLVDNVETNMEKSRRILFLLTREYFESQLHET
EINLAQVKLFDGFVDKIIFVFLEEVPKTTFKEPLKTMMRHGNCLHWPRKKRERTIFWKR
LKLALLEAKT

**>Lan-TLRβ8**

IFKNVPTLTELYLDNNQFYRILPDILKDLFRPLRQLRHFSLG------LPP---------
-----------------------------HQSMYRVPGNFA-----L---DPNGHK---
-RQLKSFHGSLA----MTTV-------CIWPPYREDA------------CF---------
------FVIFLSAVLYSFRWDILYIYAIYRASGKKSKLGGREPHKTYDAIAMYSPTSVTW
IKKHLIPNLEEDIRFKLCINDRDYIVGDPLVDNVETNMEKSRRILFLLTREYFESQLHET
EINLAQVKLFDGFVDKIIFVFLEKVPKKTFKEPLKTMLRHGTCLQWPRKKRERTIFWKRL
KLALLEAKT

**>Lan-TLRα4**

FLTSLPNFPNLEVQRNQL---------------------------ERFPN---IALPNL
WILSLKDNSITEIKNESLQHVPNLRYLSLEGNGITHIPEGFFNHTPHM-TANLTGNPIKC
DCSQRWIKDWMLQQEKRTIF---VEAFCSNSSG--AINIKDF---DFEACVPTFYVTLLV
VLVLLIIVAILILLIYICRKELQVWIIRGWWKA-NVLTNSAQRTYKYDAYIAYCDNNYSI
IRDHFIPRLEQKHGYRLFIRDRDSEAGQPIAENVANAISKSYCTIALLSNSAMESEWFPV
EFELTHSLSVEDKSRRLVIVKVGHLSKEALQKSIQLYLTTKTYLSWTD------DFWDKM
HKILPDKRE

**>Lan-TLRα3**

-LTGFPSLP-LYVNRNQVETFP-------------------------------TLFSDL
QELQAADNNIGSISNSSFTAFPKLQYINLDRNGIAEVVVGTFDSL--LSMVSLKGNSLHC
DCSQRWIQDWIIRNLT--FI-----ATCNDT-------------DFTQCDT-NVVALAV
VLVVLAVFIALVAITFFYRTEVEVLIYRYKQ--R-----DSDTDKDYDIFISYSNDDSVF
VRNVIIQKMETEWGYKLCIHERDFLPGEYIADNIANAVEKSRRTLTLLSDSYLHSEWCVF
EFAMAHQQSLKDRCRRLVVVKLSDLDSNLLAKEVGIYLKTNTFLHKGC------MFWEKV
RGTLPAKPL

**>Lan-TLRα8**

-LTALPFAP-LEMAGNLIEVLE-------------------------------PYLANA
TKLILSNNAIQTIDPAVFGLFAELRTLHLDGNHLTHLPKEITSVN--ISEIKLDKNYLSC

DCKSTWLKRWLNENGK--NIPRFTELTCAVG--QNGQRIIDVP--DSSTCDPLIVPIAIC
LAVVLVILAVNLIV-YRFSIEIKVLVYKFNWHPR--D--DDGPEKIFDAFVSYSSQDYKW
VVHNLRHTMENVPPYRLCVHDRDFIVGETIFDNIMNSVQQSKRMIMVLSQNYVDSEWC
MMEFRTAHQKVLKERSKYLIIILFDDVNKDQLDEELLAYLNTSTYLEVSS------WFWKKL
FYAMPDLSK

**>Lan-TLRβ5**

VFSHAPHLQELYMSDNHLDKIDDAALEKLFRNLTKLRKLIISQTRLTHLPPKLFETKPFL
RELQLGSNQLSSLDPVVFQSLFSLQMLYLENNLIRTIYESSLFVWKNLTKISLAENLFSC
TCDNFWFRTWMDTT--QTTIVALNSYRCYEPKELAKSPFLDWHPSKAQ-
CTPAWVIASAIGVSIMLFLALVTVVSHRYRWYIRYWCFTLRSRYKRLEPFENNGTFVFD
AFVSYNCHDRHWVIQRLLPKLEYDAGFKLCLHDRDFIVGHDIVDNIVDALEVSRKTILVL
SNNFAQSQWCQLEMTMAQHKLFDENKDILVLILLEDIKPENLSNRLTLLLRKQTYIEWP
REEEGQDLFWERVKAALQKPYG

**>Lan-TLRα1**

KLNGQPNMEIVEFSNYAY------VFITIFHGYPALHTLIAVRNNITVFPQATLLNFPKL
RYVDLRYNSIKELKI---PR-GSNRVFDLRHNDIQDLTIENVNAMRYAAHVDFRNNPIDC
GCNNSDAVKHLRSEVVKSTYRFLYDIPCHHGET--TTTIRSINLDDLNECF-IIIIPWIV
LGLLACLVVLTAILTVYFRREIQILLFRLKCRCR------S-PPKRFDAFVSYNSGDEHW
IVHTLAPKLENKPPFRLCLHYRDFIVGAAIAENIIESIEASRHTIMVLSENFLKSEWCLM
EFRAAYHQGLRERNKHLIAIVLEDILLDDIEADLRSHLRTTTYLKVSD------WFWDKL
IYCLSRNPH

**>Lan-TLRα6**

-ITTLPAMP-IYLQNNKLEIIT--------------------------------DYFSRV
HTLVASNNSIAKITSRIFWY---ISHVQLDGNILKSLPQDIESMKHNITSLSLSRNPWTC
SCENLWLKSWLLKKRK--VI-HMDSIICTNE--VKGKPISQVT--EEMLCHPSYIQVAVS
LGILLLLTLITIAVLYKYRFEVKVILHRFNWHPR-----QEMTEKLYDAFISYSSEDRFW
VHTTLAPTLENQLPYRLCMHCRDFLPGEAIDNNIIQAIQNSRCTILVLTKNFLKSNWCIF
EFQQAHYQMIHNAHFKVIVILKEDIPAEEMDDDLRAYLRTHTYLEAKD------WFWKKL
LYVMPTMNK

**>Lan-TLRβ7**

IFSAVPSLNTLSLANNSLGQT-
PAILKKMFKNLGQIWKLRLSGNGLVELPLGMFDDLVQM
TDLHLQVNQITTLPAGIFNKCKKLAHVNVENNKIISISEGVLWAIGSLRQLDLSGNKWTC
DCDIRWFVHWLRNT--RVLLSKGKQEHCNLPSDLRQLKLVDFCPAWIE-
CDNLHLTAGLTSS--VVAITLTSYLIFIFRWDIKYAWVIRKTRRNGYVEI---
PDERYAAFVSYCSKNTKWIKDELLKNVEDDMGLRLCIYERDFICGNPIVDNIEEYMNQT
TRVVFVVTGDSLQSRLCDHEFKVAQNKLFEKRITSIIFILHEDVDKKTIPDNMQTMMRHT
TCLCWPENGRQKTVFWKKIRLALLR---

**> Lan-TLRδ3**

DFKD-LKLKELYLNGNKIRTLP----PNIF-KLRELTHFDGSYNELQSIPD--IDQLQNL
KYIRLKQNRLRRLPE--LGNVKSLEVICVSENCLQDIPAEKLAKLPKL-RLCLHSNRLGQ
----A-VFQTLKKAKFHVRFDN-RLVD---P------KIKD----KIEGCH---------
------MTVPS-----------------------------GKMPIYDVFILYSEEDEKL
INDVFLPGLEEENELKVCVAFRDYIPGQYVSEEAISNMKKSRKIIALLTEHFDEQK--AV
EINQAVGADQGRQSCSVIPVVSGNI---KIPAQFEKIVPLRA------------VDWDKL
LTAIKA---

**> Lan-TLRδ4**
AIKKLTKLESLALNANEIRELN----IGIF-DLEHLIFLDASHNPIFAIPK--VQKLKKL
EYLRLKMCRLQALPE--LGDLPRLETICVSENMISKVPAEKFQKMRQLRTICLHSNRLSV
E---LKLRQ-LK----DVRLLNDQSLNC--------------------GCY---------
----------------------------------------K-KPTEKDVLIIYSGTDDRV
VDDEILPILEEELRFSAVVDFRDFIVGKPVFTQYAANRKSCRKILFVLTADFCSGK--RM
HLNEALQAVADDKRSRIIPLIWND-PNFQLPEELRSYVQLHK----NE------SRNEKL
WKALA----
**> Lan-TLRδ2**
DFKG-SQLKALYLNANYIRTLP----QNIL-KLKELIIFDGSYNELQFLPD--IDQLQNL
KYIRLKQNQLRRLPE--LGNMKSIEVICVSENRLQHIPAEKLAKLPKL-RLCLHSNRLGQ
----A-VVQILKTAKFEVRFDN-RSVD---P------KIPKI---KIEGCH---------
------MTVLS------------------------------RETPVFDVFILYSEEDEKL
ITDKFLPQLEDKAELKVCFASRDYIPGHFELKEALNNMKKSRKIIALLTEHFDEQK--AV
EINQAVDADLARQSCSVIPVVWGNV---KMPAQFKKIVPLRR------------VDWDRL
LTAIKE---
**>Lan-TLRβ1**
LFKGTSKLKILFLSDNELGYVFNDPNGMLFKNLYKLENLTLERNRISQLWPAQFQNLTS
VKNLSLSDNQVSFFTSQLFAPMTSLRALNLSQNMISLVNSSSIGGLGRLQTLDLSGSPF
ACTCDLVWFRRWINQT--NITLSQLDVYTCNTPAERRGMPLLQFDPDAID-
CVNPIYLASAVGG-TLALLVVVFISLYRWRWFLKLRYYRFKRLAKGYERV-
EGDDIVSDAFVSFCDADREWVAVELLARMDSAGRFNL-
VCDLNFLPDKSELESVVEAIECTRKAIVVLSDAYIGDPRCQF
ELEQIYESSVERQRYEMILVLKG-
LPNGKIPKVLRQRLERGEFLEWTEDANGQQLFWDQLGEKLEQRPH
**>Lan-TLRβ10**
DIRNLTRLKILHFCDNTISLTRPD--NNFFTGMVSLESLNLAGKELEGVDLTLFNPLINL
KVLNMTYTGLTKVTPGTFKPLQNLRTLDLSDNKLVEIDGDIFKYIPKLATFLFNNNRFSC
DCHLVRFVGWLKHT--SIQI---EDQPCFSPSKLSAVKVGDYSPGFLE-CK-QVLLYALG
--TLVL--LIFTAVITFYRWDIRFWWQKVRPKKQGYIPI----DGEFDAFVSYSSKDEDW
VVGTLVRNLEEEARFQLCLDNRDLIPGNFIIDNLIQGMEKSKCCLFVITRNFVKSEWCNF
ELNTAISKMLDERKNVVILIYLEHIPDKDLPKNLRRLKKHVTHLKWPNDERKIDIFWKKL
QLVLYHKKE
**>Lan-TLRδ5**
NIKKLTKLESLVLNANEIRELN----IGIF-DLEHLIFLDASHNPISAIPK--VQKLKKL
EYLRLKMCRLQALPE--LGDLPRLETICVSENMISKVPAEKFQKMGQLRTICLHSNRLSV
E---LKLRQ-LK----DVRLLNDQSLNC--------------------GCY---------
----------------------------------------K-KPMENDVLIIYSGTDDTV
VDDEILPILEKELRFSAVVDFRDFIVGKPVFTQYADNRKSCRKILFVLTADFCSGK--RM
HLNEALLAVADDRRSRIIPLIWND-PNFQLPEELRIYAKLNR----KD------YFWKKL
QKALA----
**>Lan-TLRβ4**
IFSNSPNLHDLSMHDNQLNDMNSTALETVFRNLTKLRKLYLQNSKLANLPAKMFVNNG
MLSTLQLQSNYLSTWDPIVFLPLLSLKHLFMDHNHIRILNETSFFIWTNLTEVNLAGNPF
SCTCENLWFRNWIQST--KAKVLQLHKYICYS----AKTPFLDWHPTKAQ-CTPAWVIASAI

GVSIMLFLALVIVVSHRYRWYIRYWCFTLRSRYKRLEPFEDNGAFVFDAFVSYNCHDR
HWVIQRLLPKLEYDAGFKLCLHDRDFIVGHDIVDNIVDALEVSRKAILVLSNNFAQSQW
CQLEMTMAQHKLFDENKDILVLILLEDIKPENLSNRLTLLLRKQTYIEWPREEEGQELFW
ERVKASLQIHSG
**>Lan-TLRα5**
-YTSVP-LG-LTLANNGITELKNNSINQSFTGLFQLRTLNLSFNLLEDLKEYSFSGMTML
ENLYLDHNLLTSIDPSTFASLSRLKILTLHSNRLEYLLPDVF---TSLVHLTLSHNRWPC
DCDVIYFKHWVVSYKA--IIFDVGNINCTFKRIVQGKRVLYF---DEDYCNRTHVAALVS
VSILFFLTVIVTSLLLYYRTEIKVWIFKFGCRPK-----PDDDEKIFDAFISYSSKDEHL
IVHELAPRLENHPSYKLCLHYRDFPVGASIAETIIDAVEASKRTILVLSQNFLDSEWCLY
EFQTAHHQALQDRTNRVIVILLEDIPLKNMDNELRAYMKTKTYLRWDD------WFWDKM
AYALPDVHK
**>Lan-TLRβ2**
LFSHTPNLHELNLNSNHLRHLNSTAMETMFRNLTQLRKMYIRNAGLSVLPPNMFVNKG
MLSELQLQSNSLSTWDPFVFQPLISLKKLYMNGNRISVLNETSFHIWNNVTEMDLSGN
PFSCTCGNLYFRNWMQTT--QVKLLEIHRYQCFEPKDLEKTLFLDWHPTIAQ-CT-
VWIIASAIGVSTMLFLALVIVVSHRYRWYLRYWCFSLRARYKRLEPFEDNGTYVFDAFV
SYNCHDRSWVIQRLLPKLEYDAGFKLCLHDRDFIVGHDIVDNIVDGID--------------------
---------------------------NLSNRLTLLLRKQTYIEWPSEEEGQELFWERV
KAALRRPPE
**>Lan-TLRδ6**
NIKKLTKLESLVLNANEICELN----IGIF-DLEHLIFLDASHNPISEIPK--VQKLKKL
EYLRLKMCRLQALPE--LGDLPRLETICVSENMISKVPAEKFQKMGQLRTICLHSNRLSV
E---LKLRQ-LK----DVRLLNDQSLNC---------------------GCY----------
--------------------------------------K-KPMENDVLVIYSGTDDRV
VRRRILPILEKKLGLSAVVDFRDFTIGKPVFTEYADKLKNCRKILFVLTADFCSGM--KL
HINEALQAVADDKRSRIIPLIWND-PNFQLPVELRSYAKLNR----KD------YFWKNL
KKALA----
**>Lan-TLRβ3**
IFSNSPNLHDRSMHNNQLNDMNSTALETMFRNLTKLRKLYIHNSKLANLPPKMFANNG
MLSTLQLQSNYLSTWDPIVFQPLLSLKKLFMDHNNIRILNETSFFIWTNLTEINLAGNPFS
CTCENLWLRNWIQST--KVKLLQLHYYQCYAPEKLAKTQFLDWHPTKAQ-
CTPAWVIASAIGVPTLLFLALVIVVSHRYRWYIRYWCFTLRSRYKRLEPFENNGTFVFD
AFVSYNCHDRHWVIQRLLPKLEYDAGFKLCLHDRDFIVGHDILDNIVDALDVSRKTILVL
SNNFAQSQWCQLEMTMAQHKLFDENKDILVLILLEDIKPENLSNRLTLLLRKQTYIKWP
CEEEGQELFWERVKAALQKPYG
**>LIon-TLRα2**
-LSAIPRMP-LHFENNRIEELS--------------------------------
GYLKFVVGLWMSRNDITAISREVVELLSKAKGIYFQYNKISRLSKSVMSLWKGVSKLDL
TYNLLVCDCHSEWLRHWIIEASY--LV-NGWKLRCASDETARGRAILTVE--SHEVCKT---
IIAIIFGTVFILLIVAFALVVRYRQEIKIWLYKYDWHPK--D--DSDPSLIYDAFICYSSLDYDW
AVHTLWNKLENTPPYKLLLHQRDFIPGQMTMDSIYEGVNSSKRMIMLVTQNFVRSDW
CMAEFRTAHHEVLSKNTNYLIAILGEDLDIECVPEDFKVFLKNTTYLKKDE------
NFWDRLFYALPQKGP
**>LIon-TLRβ8**
AFLGLEHIEELDLNDTDIGAV-REDLKYVFKPLKGLKRLNLADSLFKHYPVVIFQNQAEL

EELDWSSNAITAIGSVVFSTLRRLRRLDLRNNQLIYISGKVFTTLTNLQTLLMWENSFAC
NCKLRGFTYWLSRSKFKDAICESYSEPCRSPPKHVGHRLESFLPTWLD-
CENAIVALSTSLVLLFCLSVSLSVVAYRKRLSIRYWYVIRKLKRKGYIPL--
SRRHSYDVFIAYMPQEQRWVEHTFRPELEKDVAFRVATVDREFQVSGEVIDLVEGGF
RHSSHVIFIVTDEFLSWEKSDYMMTQAEVMYLEKGCPEPILVLKERITMDQAPLSYKRLI
RHVVRLHWPQGQGYTEDFWKNLRLVLLGERT

**>Llon-TLRβ4**

EMEHLTKLEKLFLSCAGTTVLP--------STLATLKQLEISEWSFREDSIQIISSLKNI
EILSLQQSNIHHFPYQELMMLDNLVTLDLSRNRISALDHKRIWKIPKLRKLNVASNNFEC
NCSMLPFSEWLRRPRPRIEIISLFNVKCASPSKYYNMALFNFDKD----CRSPIILPSTL
GP-IGLLIAIVIFVTVRYRGYIRYGLMLIRARWRGYGSI-EGCKFKWDAFVSFNGADYDW
VYNQLKPKLEDEAGYRICLHHRDFTIGEFITDNIVKCIDRSRKTLLILSDDFAKSQWCQL
ELSVAQHKLFYDDRDVLILVKLNDVSPENITGTMQVVMRTKTFITWSDALAEQDLFWK
QLILALKRPPG

**>Llon-TLRβ7**

AFTGLENLEQLELNDTDIGAV-IEDLKYMLRPLKSLKRLDLDKSRFSHIPEDTFLNQVNL
EELHLADNEIRVIGHSAFRTLVRLKYLDLRNNRIEFIHGEAMGHLTSLLTFLFTENNFGC
HCDLAGFTKWLKEHTFHENRCEWRSERCTVPLHLKDTPILDYQPGWTD-
CDNLILTVSSIFSVFLILSSSIAIHSYRRRLSIRYWYVLQKLRRAEGGTQ-
NSLDEPFDVFISFELNDRYWVEETLLPNLEDDIRFRVCTVDRDLDPGRPEVMNIARGIR
NSRNVIFVVTRELIQTAWCEYEICLAETQSLQEGNCRLIIIFLEKFTWEELPLCMKRLLSH
VNFLRWPETAHEQEDFWRRLRLVILGETT

**>Llon-TLRβ6**

AFTGLENLEQLELNDTDIGAV-IEDLKYMLRPLKSLKRLDLDKSRFSHIPEDTFLNQVNL
EELHLADNEIRVIGHSAFRTLVRLKYLDLRNNRIEFIHGEAMGHLTSLLTFLFTENNFGC
HCDLAGFTKWLKEHTFHENRCEWRSERCTVPLHLKDTPILDYQPGWTD-
CDNLILTVSSIFSVFLILSSSIAIHSYRRRLSIRYWYVLQKLRRAEGGTQ-
NSLDEPFDVFISFELNDRYWVEETLLPNLEDDIRFRVCTVDRDLDPGRPEVMNIARGIR
NSRNVIFVVTREFAQTSWCEYEISLAETQYLQEGNCRLVVLFLQKFTWEELPLCLKRLL
SHVNFLRWPATAHERNEFWQRLRLMLLGELT

**>Llon-TLRα1**

-LNKLPSIKLIYLNQNNF----------------SQISLDHSENLLTEILE--RSIKGHV
KVLDLSYSSIADIDNEFLEKLSHLTHLYLNGNKLTKLTEHTLSLQERLTELHLYNNTWDC
SCSAMLMIKLLNRLIARKTLVRPDEIVCVTPERNRGRMVYMV---DDELCE-TKLAFYIE
QHFVLNMTLALLVLKFFKRETIQLLTL------RAIVNDDD-TSMVFDAFVSYCEDDRVW
VEQELIPCLQQEPPYKICQHRLNFVPGFTVQQNEFNAIKHSRRTIIVMSNAYLGREHCQ
YEFKTAYNYWITEKEPRLVVVKYPDVEDRN-QETCHAYFRKFTYLEKDE------NTFDRL
LAFMPRR--

**>Llon-TLRβ5**

AFVGLEKLEQLELNDTDIGEI-TEDMKSVFHPLKGLKRLDLDKSRFSNIPDGMFMNQVN
LVELHLADNKLRAIDHSVFRTLLRLKQLDLRNNRLGFISGEVMARLPSLQTFFFTGNNF
GCHCDLAEFTRWLKKNDFRENQCELYNEKCVVPPSMKDTSILDYQPTWLG-
CDNLILTVSSAFSFFLILSTSVAIHAYRRRLSIRYWFVVQKMRRAEYEAL-
DNSDIPFDVFVCFEKNDRFWVEKTLLPKLEDDIRFRVCTVDRDLDPGRPEVMNVARGI
RNSRNVIFVVTRELIQTAWCEYEICLAETQSLQEGNCRLIIIFLEKFTWEELPLCMKRLLS
HVNFLRWPETAHEQEDFWRRLRLVILGETT

**>Llon-TLRβ2**
ALHGIPNLEVLDIHGNKFDDMTQD--
ANFLSMFRNLKRLSMGKMGLFFLEGWIFDNLTKLERLELSFNALGNITARWFKNLKYLK
KLEMRDCRIATVNVKSFAFLNQLNSLDLRDNPFSCDCSIQWFLNWSKHHGNQLYMFN
RKDYTCASPQWLHRMPLRKFTIP---SCYKTLLTGLSV
AGVAIIVCFFVLAFFSRYRWHIKYKLFKLKIWF-QYEEL-DGSKYEFDYHVHYDDQDVSW
VVNTLIPELEDKRGYRLYIKHRDSSLCQYIIENIRYSIEHSYKTVLCLSNQFTQNPGTQF
LLSFIINKLVNEKKNILVCILLEEIQGENLLETLEEVLTEKSYIRLPEDREAMEYFWSRV
DEALH-PRN
**>Llon-TLRβ1**
VFQNLKRLKILRLTNNDLGPQLKDTKGELFAGLENLEELYIEKNDIQELTGDVFRHLKGA
KMLELGENAISQWGTSTFSQNSTLKHLNLSRNRIATINEPSLADLKLLTTLTLTANPFSC
DCGLVWFRRWINST--NVTFPELELYTCNSPVRMAGIPLLKFDPDSLT-CKDPYILGGSL
GG-VVILILVISLVIYRYRWFIKLRAYRIAKAVAGYEPI-PGDDLRFDAYISNHRDDRRF
VLDELLQNYDNNGGFRLCFNERDFVPGEYDLTNVTENMSQSQRGLIVLSLQYIQDHFQ
DFELHLLLKEANLRA-FGLIVIELEEIPPNRIPNGLRRIFEEGDHLSWSEDPNQQQL
FRERLTNKLQRRPQ
**>Llon-TLRβ3**
PIAGLNKLYKFGTTGADF----IN--RGLFINKTELQSLHIGSGRITMNHLDILRNVTSL
KSLRLEKMDIREIPH--ILHLKHLKSLTLVSNKIEMIPQHFIPKLDGLKTVDLAFNPFAC
NCSLMPFSNWLRNASRLATVTNLDVTLCFSPASYKNTPLFNFEKD----CRNPIILPSVL
IP-LVLLILVVTAVSVQYRGYIRYACMLVRARWRGYDALNEGRSFKYDTFVSYNREDA
AWVLRVLRPKLEDEVGYQICLHDRDFTVGEDIVDNIIMSIDESRKTILVLSDNFAKSQWC
QLEMSLAQHKLFEDNRDVLILIRLGEVAEENMTRTMRMLMRTKTYITWPQNEEGIDLF
WRNLIFALRRPPG
**>Lrub-TLRβ2**
FFKGLTNLQNLSLSWTDFGEVDHSQGADIFDQANTLRSLNFLGGFITHISAGLINDLHNL
THLNFQSNSIGYLFSEWFENLTNLKVLNLNDNKITTVVGANGRFFRSLSQLTIGENAFD
CDCQLRFFSEWLRGLDDQIRVSDMDKAVCLTPIAYENKRIKDF---KSAVCS---GIAFTS
AAAALLFIILSVIVGCYCRHDIKYMMAIRRLHHR---TLR-GSKLIYDV----NDTDRQW
INTNIGNITDKD--FNITTNHPDIVPGEARSNPLSKQINQCYYTLLLISNHFKDDLWPEV
HANLTVQEI---HNIRFVIVLIDNLRLRDLPRELKALAKQRPCFKWPTETLRRRLFWRQL
ILALIKRRA
**>Lrub-TLRα1**
-LNKLPSIKLVYLNGNNF----------------SQINLDHSEKLLTEILD--KSIKGHV
KVLDLSHSNISDIDKQFLEKLPYLSHLYLNDNMLTKLTLNSLRIADRLTEIHLYNNTWHC
SCNMTQMTILLNHLIARKTLVRPDEILCATPERHRGRRVYML---DHELCG-TDSMSFVV
LTSIIAITLATLILKFFKPKTLQVALP------RAIEDDDD-TGMVFDAFVSYCEDDRVW
VEEELIPRLK--PSYKICEHKKHFVPGITVQENEYNAIKHSRRTIIVMSNAYLERGHCRY
EFTTAYTYWIIEMKPRLVVVKYPEVVDQNK-ETCHAYFRTFTYLAKDG------NTFDRL
LDFMPEK--
**>Lrub-TLRβ1**
ELHGIPNLDVLDIHGNKFDNMVQD--ANFLSMFPNLKTLQMGKMGLFFLKDCIFDNLTKL
EKLELSFNALGNIPARWFKNLKHLKTLEMRDCRIATVNEKSFEFLNQVTHLDLRNNPFS
CDCSMKWFLNWSKYHNNRLVNYNQKDYTCASPPTLHGLPIKNFTIP---
SCYKTLLTGLSAAGVAVLICFFSLALLGRYRWHIKYKLFKLKIWY-QYEEV-

DGSKYEYDYHVHYDDEDVTWVLNTLIPELETKRRYRLYIKHRDSPLCEYIIENIRYSIEH
SYKTVLCLSNKFTRNPGTQFLLSFIINKLVNEKKNILVCILLEEIEGENLLDTLEEVLTERN
YIRLPEDREAMEYFWTLVDQALH-PRN
**>Lrub-TLRα3**
-LKAIPHLTNFIHKDVQV----------------ESSILILSHNNINSLNP--YEHLKHF
TQIDLSHNNLSFIDPSWFNLFKGVSQLHLNDNNLTQINPKDIETFRNLEELHLYNNPWD
CGCNKIWFKSWLSYLADAGVVMKPHLITCASHHWNKEKIVHNL---YVSFCQ-TFIAMVLI
--VLVTIFFIFILM-----YKLRLYIFSFNVHLR-----EEGENMESDAFISYNGADYDW
VKNKFNIRL---MKYKLC-NLRQAVDGRDFLENIETALMTSKRTIVVLTKHYLEDEECTR
EFKIAREYWVDIKRHRLIVLKH-GVDIDEIDNEIRLFLKRYKLIEMDR------NLWRNL
SYAMPNP--
**>Lrub-TLRα4**
-LKAIPTLNGLIKPDNKI--------------------LLLTGNNITNLNA--VQYLDSF
TTIDLSHSNISAINEYTL--LGKVSKLFLNDNNLTHLLVQGLKIFTNLEELHLYNNPWNC
SCDQKWMKLWLLNYIDAGIILKPHMVTCGSQNWNEGKVIHTL---PEEFCFPTILYIFLV
---LLVCILVG---VKQFIYNNKYQLFEFNLHPC-----EHGENMNYDVFISYCENDRSK
VLKDLIQSRD--PPYKICDPDDSFPRGRPINESIADAVTKSKRTIIVLTKAYALKSYCCE
EFRTALDHWSMDERHRLIVI--KDIETNTIDDKLRQYLREYTYLELDV------NLWKRL
SYILPQP--
**>Lrub-TLRα2**
------------------------------------------------------------------------------------------
TNWPGHIQSFEDKVTEIHLYNNPWDCSCNKLWMKKWLNKLIV--ILVKPDKIICGTP--
NKGQMFYMV---EDSFC--TDKKIIIIL--VISIVLAVLILRH---VFHKLWPDPLPPNRI---
DDSNTSGMTFDAFVSYCEPDSEWVEQELIPNLQQNPPYKVCHHKQYFQPGMPSEWN
EFMAIIHSRRIIIVMSNSYLEREHCRFEFSTAYSYWIHTKNPRIVIVKYPDLEIVN-
REACSAYFKRFTYLAKDE------T-FKRLFQFMPLE--
**>Nge-TLRβ6**
VFLGLGHITELELNMTDIGKI-TEDLEFVFRPLKNLERLNLANSALKDFPRDLFLNQVNL
KILDLSYNRITIIGDVILSTLKKLRYLDLRRNEITEISGDALRKLTYLQIALIAKNNYAC
TCALRGFTKWLLEHKVTDDVCADYSAGCRSPPQFVGKRLDKFQPSWID-
CDNAIVTLSTLFTIIFCLFTILSIYGYRKRLSIRYWYVIRKIKRKGYLPL-
PALHDLHDAYVAYADNDHAWVEGTLLPNLEDDVTFKICTNERDFQVSDEIIDIVEKGIKC
SDRIIFLITNAYLQATKSDYVMAQAERLYLETGRPHIILIMKEKINFDRVPLSFKRLISHAA
RLHWPENQGQRNDFWKNLRLLLLGERT
**>Nge-TLRβ5**
VFLGLGHITELELNMTDIGKI-PDDLEVVFRPLKNLKRLNLANSAINDFPRDLFLNQVNL
EFLDLSYNRITIIGDVVFSTLKKLRYLDLRRNEITEISGDALRKLTHLQIALIGKNNYAC
TCALRGFTKWLLEHQVGGEVCADLSASCRSPPQFVGKRLDRFQPSWID-
CDNTIVTLSTLFIII----VCLSIYGYRKRLSIRYWYVIRKIKRKGYLPL-
PALHDLHDAYVAYADNDHAWVEGTLLPNLEDDVTFKICTNERDFQVSDEIIDIVEKGIKC
SDRIIFLITNAYLQETKSDYVMAQAERLYLETGRPHIILIIKEKINFDRVPLSFKRLISHAAR
LHWPENQGQRNDFWKNLRLLLLGERT
**>Nge-TLRα**
---------------------------------------------------------------------------------------------------------------
------------------------------------------------------------------------------------------------------------
NSLDYVWVVHTLWNKLEKRPAYRLLLHHRDFIPGGMIMDNIVEGVTKSKRMIMYVTDN

FIKSQWCMVEFRTAHHEALSKNMNYLIAIVDEELDIENVPEDFKVFLKNTTYLKRNE------
HFWDKLYYALPQRGP
**>Nge-TLRβ2**
VFKNLKHLKKLKLGKNELGGLIDDKKGELFAGLDHLEQLDLRMNSVKELTDGVLKPLKG
MKKLELGRNSISAWGPATFSQNRTLQHLNLSNNNIATISKSSMSTLTSLVTLTLTGNPF
SCDCGLVWFRRWIDHA--NVTFPGLKSYQCNSPPVREGLLLLKFDPNSLT-
CIDPYVLGGSIGG-AVILCLVMTLVMYRYRWFIKLRAYRFGQAMREYEPI-
PGDDLHFDAYISNRDSSREFVLGTLLPVFDNNGAYRLCFDERDFEPGEYVLTNITNNIA
QSQRGLIILTPEYIHDKFYELELHMLLEEANKRP-
FTIIVIELVEIPPNRVPNGLRRIFEARNQLTWSENPDEQALFKDRLTNKLERRPQ
**>Nge-TLRβ4**
------DCGYLNISYFKFDDQLELANPVPFRLTPNIAEFMVSAARCFVQPQYK-------
---------LVSLLRAILRDEYITWHKKMFLLRVVAVVLPLLADGRRASDVEYVKKGVNV
NCTEKGWKDVPKNLPKKIASIRRAFVGLTKLESLDLTANKIRXIPQ--RCHKRLLAGLCA
FGVLLIVTALGLGLFSRYRWKIKYKIFKLRLWFYQYEEL-
DGSKYEYHFLVHYDDSDFPWVRDMLIPELEHKRGYRLYIKDRDSRLCEYILENIQYSIE
NSYKTVLCISNQFTQNSWCQFLLRLLIQKLVNEKKNILVCILLEEIGGENLLDTLENVLTQ
KNYIRLPEDREAMAYFWTCVVEALH-PRN
**>Nge-TLRβ1**
VFKNLKHLKKLKVGKNELGGLIEDKKGELFAGLDHLEQLDLRMNNIQELTDGVLRPLKG
MKKLELGRNSISAWGPATFSQNRTLQHLNLSNNNVATISKSSMSTLTSLVTLTLTGNPF
LCDCGLVWFRRWIDHA--NVTFPGLKSYQCNSPPIREGLPLLKFDADSLP-
CIDPYILGGSIGG-AVIICLAMTLVMYRYRWFIKLRAYRFGQAVREYEPI-
PGDDLEFDAYISNHHSSSEFVLGTLLPNFDNNGAFRLYFDERDFEPGTHDLTNMGKKI
SQSQRGLIILTPEYIQDKFHELELHLLLEEAKKRP-
FTIIVIELVEIPPTRVPKGLRRVFEARDQLTWSENADEQVLFKERLTNKLERRPQ
**>Nge-TLRβ3**
---------------------------------PEFEDFEVVSRKVFEVERWSFDKAVHICGWVLRF--
VYNLRHPNLRHSGPLSHEEMFLLRVVAVVLPLLAHGRRASDVEYVKKGVNV
NCAEKGWKDVPKNLPKKISSIRRAFVGLTKLESLDLTANKI-------------------
---------------SRYRWKIKYKIFKLRLWFYQYEDL-DGSKYEYHFLVHYDDSDFPW
VRDMLIPELENKRGYRLYIKDRDSRLCEYILENIQYSIENSYKTVLCISNQFTQNSWCQF
LLRLLIQKLVNEKKNILVCILLEEIEGENLLDTLENVLTQKNYIRLPEDREAMAYFWTCV
VEALH-PRN
**>Pau-TLRγ14**
LFRAVGKLEHLFASRNMFGLFKPGDLVTTLSGLPNIKTIDLSINRLSYLPRGIFSECPNL
THLDLQRNGMKSVHL--FSSLPSLKYLDLSHNEIKGFSKEQTEDF----FLKLENNSFEC
SCDNIPFIEWIQSDGSNDTVLNKSKLICEFADG-RMTALVDVDLNKLYDCMKIIIAVATM
----AVVLLSTSLIMWRYQWYIKYWIYVLRLRHR------DDGTEPFASYISYADNDYDL
ANT-VCTKLEE-SNLPVFFRDRDTSLGTSIFDEYFRGIESSRKCILCLTDSHLNCAERYF
ELQMSMVR----GKGFLIPVVVGNLALEKLPKPLRRLLRDDVYFEWPKTDLEEEDFWKSL
IAAVLTRKG
**>Pau-TLRγ6**
---------------------------------------------------------------------------------------------------------------------
-NCRRSLLGF------------------------E-EH-QICIKRTWGIAKCHRS---LDAVFSYCLH-----------
-------------------HQHLILAAFTGKMVTCLSSEDRHWVHDVLRARLEENSDFGLCIHYRNFL

PGRNIEENVIDAIESSRHSMLVVSRNFLKSEWCIFEMHMARNIFRRQQKDVLLLILLEDI-
VQDAPLTLVNLLRSRTYLKWPADDVGQEAFWERLKETLKREPE
**>Pau-TLRγ5**
RYKAFPALKKLILAGNKLYIMLRDRKTKHFRYLNNLTHLDLAYNSINELYPETFNDLPAI
KEVLLRGNRLISVTSMNITGMPSLKNISFAKNNVRFVSENVLHLWHG-
KSVDFSQNPFNCSCLFLPFLRWFNNVSSTVTLLNSDHYRCGDEKKT---
YVRKISLDALTKCTSAWMIISIAMSICVALCITLGSVLYRHRWTISFWINFAARKSHSYSP
H-MRQRFQYDAFVIYSSEDRHWVHDVLVTRLEDESDIGLCIHYRNFLPGRDIEENVL
NAIENSRHSMLVVSRHFLKSEWCIFEMHVARNVLRQQRKDVLVLILLEDIPVQDAPLTL
VNMLRTRTYLKWPANDVGQEAFWEMLKETLTQEPE
**>Pau-TLRγ2**
IFQNLTSLKTLDLSHNLLADKLKDEYGTIFQNMTTVVKIDLSGNGIYKLHINTFHHLKKV
REVILRNNRLASLPAVHIEYLTALKLVDMTSNRVEYLSKASLDSLDK--TVLLADNPFNC
TCEMLVLFRWLHEYLEGLHIKDNHSVYCSHNTSLR---LSNFQFDDLQKCT-LWIWLSLS
TILLLTALIVGLGVAYRRRWTIRFWLIAAR---QGYHRL-PMPEYKYDAFLCHSGENTWW
A-KRIQDHLEDDIGMKLCIYYRDFPVGVPIVECVNDAIVDSRYIILLITKSFIKSQWCIY
EFFMAKSKVFCENRSRLIVLVMEKLTDDVLPRTLQNLMKDSVYLEWTDNALGQEQFW
KRLRERLRTEPP
**>Pau-TLRγ13**
QFKAFGNLEHLMASRNRFGLFRSEDLVTTFSNLPRIRTIDLALNSLTSLPEGMFSQCPL
LEYLNLERNGIKVVNI--FATLPSLKLLDLSHNEIKGFTKDQTDDL----VLQMNNNSFEC
SCGNIPFIEWIQSDASDGIVLNKNNLTCQYEDG-KITSLRAVQ--GLYQCIKIIITVSSL
----TSVALCTALIVWRYQWHVRYWIYILRLKRR------DEVSKCFDAFISYSDEDHNL
AAS-VFRKLEE-MGLQIFFRERDTVIGACVLDESFRGIESSREIVLCLTESYLNSDQCYF
ELRMSMLR----GKGFVIPVVVGDVALEKLSKPLRHLLREGVYFEWPTLESEE
KDFWKSMKKAVLTAKG
**>Pau-TLRα2**
-LTETS----ISLAKNNLSNIGHNYIVNTFKDANCLKELRLDNNKLTQLKRYYFESLNNI
VELWLQNNDITSIDKDSFIHLTHLQRLFLHRNHLHTLPESKL---DSIVEVTLAQNNWSC
ECSFASFRHWLLDHID--IMADITNITCTATSTSRGEELVSF---DINFCNQRLISGLIA
SILTLVTITLSLSLLLKYRDTIKVWLYRYGWRPS-----SADVRKKYDIYLSSTNTEA--
-CRELLAELEDLPRYVVFFPQRDLIPGGVTTNDITEAIKESWRTIVVLSPAYLQDSWRMF
EFLRAHYCSVHTKTNRIIVLLSEPMKADDMEKDIQAYLTSKSYIKLWE------RLYDKI
RYRLPDGRK
**>Pau-TLRα1**
-LSGIPE---VSLANNEITELNNNDIPNTFRELACVKVLRLDHNQIAHLAAYIFTGLRQL
RELDLQSNLISSIDNRTFHQIIHLEKLQLHDNRLVSLPEPNT---TSMKHITLSGNPWTC
GCDFAQFRRWLILHMD--IIPDILEVKCTLKQTSNNRKLIDLSLINIEYCYESIRNALIS
TIIIFICLIITTIVVYTNRMEIKVWIFRYGRRPY-----KDDFSKPYDAYISYSDKQLNF
VIHELLPKLEQSPHYKLHVRARDDLPGGVRANDIISTLENSCRTIAVLSENYVADEWCLF
EFQRAHYNALHSKMHSIIVVLLHDVKA-DVDKEIQLYIKTGSYMRRDD------KLWQKI
RYALPDTRK
**>Pau-TLRβ3**
YFKGLKKLDEILLGRNDFSEFDRKSPIEIFSDLRSLRKLNLNYVNLKYLHDGFFAALKNL
TKLILDGNGFSGWSPRAFQYLVSLQHLSIINCQVFTINSSMLQPVATLTRFEGYGNPFA
CECKLRWFISWLEMMNSSTHV---KPYKCLTPKKWHGHSIFEYNTTDDD-

CSDWLLITATASGTVVFVTAAISGIAYYHRWSIRYWMFLARSRRKKEISLRRREDFEYD
CFVTYSSLDTDFVVQEMLSHLEGENDLRVCIHERNFQVGGDITDNIVQSIETSRKIVVVL
TENYVKSEWCKLELNMAHAKLLDERRQALIIIMKEKVSVKLMTPILRHLVRNQTYILWNG
SDILQTAFWGKLVQAIMKP--
**>Pau-TLRβ2**
YFANLTSLEHLSLNFVDLGIDIDSKPECLFEGLVKLKVLDLSSTALNGLPERLFKDLTSL
ERLILRKNQLSGWNGAVLENLKNLRSLDVSMNQIRTINQSSLRPVSTLNHFQGFHNPYI
CDCNLAWYVDWVGQMHATLKISHKISYKCSN---LKNKSLLQYRPTFFE-
CHRLAILCGSGFG-LFTVIFTVIGLLYKHRWYIRYWIFLLRSRRSTHLEETDGLLYTYDCFI
TYSGEDSNLVTQQLLPKLENEFGYKMCVHERDFKLGREISENIAESIEKSRKVLVVLTQ
NFVQSEWCKFEVNLAHANALHNARQSLIIILVEDVSFEHMTPILRFLMRKKTFLEWTND
AQGQTLFWERLKNAIQQYGQ
**>Pau-TLRγ10**
----------------------GKGLSTIPRNLPNATVLILRRNSITSIPANIF------
--VLLQRNNIAVVD-ATFKKLPFIKLMDLGHNYIKKFSFEQVEDL----TLRLTGNLFEC
SCKTVDFITWIQAPGTSNVIENKKDLKCASPDG-THQRVVEVSLQGLGYCVQLIISSAVV
S----AMGICVAVVIWRHKWTIKYWMYLL----KRRR--GIDRIRPRHAYISATDDDLEK
ANMIFQQ-IEDKLENSVFWKHRDTVPGRSTFDEIFRGVEESRKVILCITQSYSTCTQ-NF
EVEMSF----ARGKGFIIPVLIGDVPLERLPRPLRRLLRDDIYLEWPNNVAEMPNFWVSL
HEAVMTQKG
**>Pau-TLRγ7**
LFSHFPSIKVIELQDNLLGLEMPDQFADVFENVRTLLEIDLSSNYLNKLAAECLAGNAKL
TRIHLANNKLDKLG-LRLENFPMLEFLNLSKNAILFLSSNETSALDSI-ILDLSGNILLC
SCATLNFLDWLRT---SVIFSGRNTYTCSY-KG-KPRNLEDVDLENFRECMSVTVLTSSL
VA--GTILVMLATVGWYRRGHIRYVIY------KFRQ--HPNDDLRYDAYLAYESRDCDV
AVEMAAI-LEGDHGLDIYIHDRNAPVPGDHYDSIFDGLGRSKKVILLITDHALRSESWSF
ETDLSL---SIKGKGKILCVVKGHLSIGRLNRKLRYLMADDTYLVWPEDNDVEKTFWRHV
AVAITSKNG
**>Pau-TLRγ15**
FLAYLPSIEVIELQDNLLGLEMPHQFARVFENVTTLTEIDLTRNYLHNLTAECLTGNENL
AKIHLANNKLDKIGL--VEKFPMLEFLNLSKNAILFLSSSETSALDSIAIVDLSGNIMLC
SCATLNFLEWLRTA--SVTFPKKSSYTCTYKGK--TRNLEEFDYEDFRECFSITVLTSSL
VA--GTVLVLSVVIGWYRRWHIRYIIY------KFRQPPEPNDGHRYDAYLAYESRDHDK
ALE-MTAVLERDHGLKIYIHDRNAPLPGDHHDSIFDGVGRSKKVILLITDHALRSQWWSF
ETDLS---LSIKGKSKILCVVKGHLSIGRLNRKLRYLMADDTYLLWPDNENAANTFWRNV
ALAITSKNG
**>Pau-TLRα6**
-LRSFPKLP-LHLEDNFLEHVE--------------------------------EYLKRV
TKLFASRNNISDVSEKVLKKMEKITVLYLDSNKLTTLPEYIKKMTRRLTHVNIKHNFFEC
DCNTLWIKYWLRENIA--KVIETQNILCSSG--TKGKSIIYVP--DNKVCEL---VAAIV
LAVTLTIFLVAVVSVYKHRQEVKVLLYKLQWHPK--E-LDEDETKIYDAFISYCQKDYRF
VCNDLRSSLEQNPPYKLCIHERDFMAGAPIYENIMNSVKLSKRMIMILSNDFLLSEWCM
LEFRTAHQKVLKEHSRYLIIIALGDIVSRNTDEDLQAYLKTNTYLTVDD------LFLERL
RYALPRPTS
**>Pau-TLRα4**

-LTSLPKLPRTNFSNNHLTEVT--------------------------------
AYFPNIIDLDISGNNIRNVSDAALIQLRNIKVLNVAKNKLTTMPRRLLESSANSTAISLSGN
SWNCTCSEVWFIKWVLSKSS--VVTDSHGLFCSHP--MRGKRFSDDV--
VTERCDADYTAVAVSVGVSSTVLLIVIVIVTVFSEDIKVILFKWNIDIR--N-
VDNCSDRNFDAFVSYSSLDGDWVRNHLLPLLENDPPFKTCFHERDFIPGLPITENIIQAI
QKSKRTVLVVSKNFIDSEWCQFEFLTAHKTFLETKENKLIVIVVESVNLRSLNPKLRAYF
NTKTFLKVTD------LFKEKLYYAMPRLME
**>Pau-TLRα3**
-LTSLPAAP-FHLSKNSINKIE--------------------------------DYLTRV
FNMDLSYNNISVIDEDAFQNMKQVQSVDLRGNGLTTLPLLLKQGTRNLQKIFLGENKYN
CSCENAWMKNWLKRNSN--ISAGLEDIVCDSP---KKFRAIKVI--EADNCREKFTQEVVV
SICLCLLVLVLSIIIIIYRDLFRVLMYHFNVRIH-----EEETDATYDAFIAYSSLDGEW
VRNKLMPLLENRKPFKVCIHEREFLPGLSVADNVHRCMDLSRRNVMVVSQNFINSEW
CRFEFQAAHAATMRNKSKRLLLIMLEDIRQDNLGDDIKSYLKTNTYLEAEE------
WFKPKLFYFMPSVKS
**>Pau-TLRγ4**
RWQATPSLKKLLLTGNRLYVMLRDKKTKHFKYLKNLEHLDLADNNLAEIYPDMFRELPV
IKEIVLRRNRLFSATNLDVSGMPLLKKVNFVNNKIRFVSEGALKLWHG-
KSIDFSGNPFNCSCHFLPFLEWFNNASPTVTLLNSDQYLCRS-----
GAYIKNVDIKRLNQCKKTWRVISITVAIGVALSITFGGVLYRYRWTIKFWIVFAARRSR---
EVDRCRKFRFDAFVIYSSEDRYWVHDVLRTKLEDGNDFGLCIHYRNFLPGPPIEENIIG
AIENSRHCILIVSRNFLQSEWCIFEMHMARNVFRQQQKDVLILILLEDVPVQDAPLTLVN
LLRTRTYLKWPADDVGQEAFWETLKDTLRQEPE
**>Pau-TLRα5**
-LTNLPKLP-LQLSDNRIEELK--------------------------------DYFHRL
LELDLSNNGLRTMSDIALVKLTNITTLKLNGNRLRTLPRSTETWSQSLRQLALHDNLWE
CTCDTMWFRDWLIQLGS--VVQEPDSIMCFKD--EEWKPIKKA-----ILC--DYIPLAIT
VSSVSAVLMLAAVLMYIYRMEMKLLIFRLNWHPR-----TEILNKKYDAFISYSEEDSMW
VRRLIQLLEVDDPPYITCFHHRDFIPGVSTAANIEMAVHDSQCTIIVLSPAFVQSEWCMF
EFQVAHACLMDNEIGKVIIIIKEDIEVKKLQPDLKSYLRTMTYVKASD------WFSEKL
YYALPQKDK
**>Pau-TLRγ1**
ILQNMTSLRTLILSDNSLADKLADEYGSLFQNMTSVVEIDLSYNGIYVLHSNTFYHLKNV
RKIILRNNRLASLPSEHTENLTVLSFVDMASNKIQYLSKAFLDSLNR--TVLLSGNPFNC
SCGMLVFLHWLADYDDEDKIQDYRKLHCHRKPRLH---LSDFRFADLEECT-LWIWLSII
TMCTLAVLTVCFGVTYHKRWVIRFWLVATR---KKYNRL-PTTQYTYDAFLCHSSEDVRC
V-ERMRERLEEGSRLKLCLYYRDFPLGVPIIECVNEAIADSRYILLLITKNFIKSQWCIY
EFFMAKTKVFCENLSRLIIVVLEELTNDVLPRTLQSVMRDNVYLEWTDDVQGQEHFWQ
RLEECLGTEPP
**>Pau-TLRγ12**
LFRAIGKLEHLFASRNRIGLLTSQDLSATFSSLPNIRTIDLSLNQLSSIPESMFSLCPHL
ERLNVQRNGMKSVNL--FANLQSLKFLDFSHNEISGFTKEQTIDL----ALQMNNNSFEC
SCNNVPFIEWIQHESSNDTVLYKENLTCLYKDG-REEAVISVDLGGLYECIKIIIIISTL
----ISVVLCTALVAWRFQWHIRYWVYILRMKRR------DDV----------SKKXYDLALS-
VLHKLEE-MGLLAFFRDRDTDLGVCVLDECFRGIESSRKSIVCLTENYLNSGQRYF
ELRMSMLR---GKGFVIPVVVGDVALEKLSKPLRHLLRDGVYFEWPTLESEEKDFWKSM

LKAVLTPKG
**>Pau-TLRγ11**
MFRAIGKLEHLFASRNRIGLVTSQDLTATFSSLPNIRTIDLSLNQLSFIPEGMFSLCPHL
ERLNVQRNGMKSVNL--FANLQSLKFLDFSHNEISGFTKEQTIDL----ALQMNNNSFEC
SCNNVPFIEWIQLESSNDTVLNKENITCLYKDG-REEAVISVDLGGLYECIKIIIIVSTL
----ISVVLCSALIAWRFQWHIRYWVYILRMKRR------DDVSKNFDAFISYADVDYDL
ALS-VLHKLEE-MGLLAFFRDRNTDLGACVLDECFRGIESSRKSIVCLTESYLDSDQRYF
ELRMSMLR----GKGFVIPVVVGDVALEKLSKPLRHLLSDGVYFEWPTLESEEKDFWKS
MLKAVLTPKG
**>Pau-TLRγ9**
FFRDFPSLQTLQLASNRFGLFTESYLEAIFSNLPSIRKIDLADNLLTTVPKAMFSNCTSL
VTLNLRYNPLVTFEF--FSLFPQISYVDVGDCKIQEFKAYQVKFFSSL-KVNVSGLDLDC
NCENKEFYEMIQKNKSLADLEGKQELACTRDGV--RVKLAHLGLSGLESCMYIFVILCVT
---VVSAILISVVIVLYCRWNIKYLVHLTKRKLK-----NQANALYYDVYLSFSEDDRDT
AFQ-LFTGLNN--GLEVFYWPRNSRPGTCQFEEIFEQMGLCKKIVILITASTENSAMQNF
EIRMSLPR----GKGFVIPIVKEDYVIGKLPGPIKNLLRQDLFFLWPEQEKDQEMFYRNV
KRAARSKDG
**>Pau-TLRγ8**
LFRAVPSLQTLLLAVNRIGLFTETELVAIFSNLQNIKEIDLTDNMLTTIPKVMFSNCTSL
VILNLQKNPLLTFEL--FSLFPGFTFIDLSNCEIQEFKYFQTAFFSSFRVVNVSGLSLSC
NCNNKEFYDMIRNNKSLVDLEGREELTCTHGTE--RIKLVDLDLSSLESCWYTFLIVCVT
---LTSVVIIGVVVVLYCRWNIKYLVYLTKNKLR-----NNANIFEYDLYISFSEDDRDI
AFQ-LFTGLHNK-GLDVFFWPRNSRPGSCVFDEIFEQLDGSKKVLVLVTSSTEN
SVTQNFEIRMSMAR----GKGFIIPVVTEDFVVCNLPQGIKHLLRHDLYFLWPEEDEEKE
EFWKNLERAITTKRG
**>Pau-TLRγ3**
SYLPCPNLRKLILAGNRLYVMLRDKGTKHFANLPNLEYLDLANNNLTELYPEMFSELPAI
QTIVLRGNRLFAVANMDISDMPSLKKIIFARNRIQFVSEGALQLWHG-KKLDLSQNPFNC
SCQSLSFLRWFKNVSSTVTFLSPHGYLCRDKFHQKSEYVGEVDLKRLEECRKDWIVISI
TVSTGVALLITLIGMIYRHRWTIKFWIVFAARRG-PINSLDRQRRFHYDAFLIYSSEDRHW
VHDVLREKLEEDNEFGLCIHYRNFLPGQPIEENIMYAIENSRHSILVLSRNFLKSEWCIF
EMHMARNIFRQQRRDILILILLEDIPVQDSPLTLINMLRTRTYLKWPADDVGQEAFWEML
KQTLKKEPG
**>Pau-TLRβ1**
----------------------------MFEGLVKLKVLDLSYTNLKGLPERLFKDLTGL
ERLILRQNQLSGWNDVVLRNLKNLRSLDVSMNQIRTINQSSLRPVSTLNHFQAFSNPYI
CDCNLAWYVDWVRQMHGTLKISYKIPYNCSN---LKHKSLLQYRPTFFE-CHRLVILCG
SGFG-LFVVILAVIGLLYKHRWYIRYWIFLLRSRRSNHLEETDRLLYTYDCFITYSGD
DSDLVTQQLLPKLENEFGYKMCIHERDFKLGREISENIAESI-------VVLTQNFVKSE
WCKFEVNLAHANTLHNARQSLIIILVEDVSFEHMTPILRFLMRKKTFLEWSNDTQGQRV
FWERLKDAIQQRGQ
**>Phe-TLRα**
-LNGIPE---VSMANNNLAVLNNNTIPDAFMNVDCILVLNLSQNKLTYLDASMFNGLKDL
RELHLQENNISTIMKDTFQRLQKLEILHLHKNSLTVLHEPSI---GSLKKLTLANNEWVC
DCEFAPFRQWLIEHMN--IIQDLINITCVIKEKSKNRKLIDLSLTNIEFCYESLRNALIS
VIIIFISIGVTSTLVYKYRNSIKVWLFRYGWRPY-----QDDFSKTYDAYVSYSDKQLNF

VLHELLPKLERAPQYKLYLRDRDLIPGGVQANDIIEAIEDSCRTLVVLSENYTTDEWCLF
EFQRAHYNALHNKNHNVVVIKLHDIQTEDIDKEIQLYIKTGSFFKRED------KLWEKV
RYALPDMRE
**>Phe-TLRβ**
----------------------------------------------------RFFKDLSQL
LTLELGQNKLQGWDPEIFKNTTKLQYFSVYSNNIGTLNKSSMHLLPSLKRFDAYSNPYV
CNCDLIWYCDWLRNMQRKVTIRSDRPYNCSN---LKKRTLLSYNPTFFD-CHQLKIILPSG
FG—FVFVVFIAGLAYRYRWYMRYWLFLLRSRRNKHLEEHERLCYEYDCFVTYSG
EDSEWVIQEMLPKLEQEFRLRACIHERDFELGHDIYENIAESIENSRKVIVILTKNFVKSE
WCKFELNLAHANTLHNACQKLIIVMKECVPMNIMTPLLRYLVRKRTFLEWSNDEQGRT
LFWNRLNIALTTAAG
**>Ese-TLRα**
----L--LNNLDLSSNRISYIP----PRLFYKLEYLATLNLRSNQLTELDI--YFYLPRI
QTIDLTFNRISRFTNEVINNLPSLKQADLRNNLITSFDDYVLRLYRSL-TMRLDNNPLNC
DCAKI-FTQLLRNSVDTTNI---FRALCQT---FNGKSIFNFSLN---ACSSLFQIAGYV
IGLLLLLLMILYCLILAICFNCIPFFYVCPCKS-GVK-----RDKEYDLFISYNRANEKW
VKEQLVPFIKENENYILHYNNEN-KLDEVFGPYVKDIMSKSSCILFILSDAFLLKEWNNK
DLRQHLRYLITKEKTRFICVQMHDICDEEVEEYFTDKLQIPRFVSLENDE----LFWKKL
AYYLPKPKS
**>Pcau-TLRα2**
-QSEIPNVS-LRLDGNNVTTIHNSVIPGSFRDLNNLIALYLDGNEIEEIGDDQFNGLADI
KELHLENNMLVNISTTWIDVTPMFSMLALHGNAFSKAPEAIY---IRSSEYTLRQNPWIC
DCTDEFFLDWLRSNVD--NISDIGEMMCTIPRAITVIEILDF---EMVYCTAGFIAGFVM
LGVLFLTTVLCIALTHHYQHEIKLWLFKYGVRVR---DPESDKAKKYDAFISYHNSDEDI
ILREFVPQLEHETPYKLCVHNRDFLAGEFIAENIVYAVENSRRTIVLLTASFIDSEWCRY
EFQAAHNQAISEKVNRIILVVFEDIPKGKLDKNLEAYIKTNTYLRYDD------MFWSKL
RYALPAVRA
**>Pcau-TLRα1**
CLEKIPGLTKLSIAHTTIRSVV--YMRDFFKYHPNLTYLDMTGTRFSSLTTEAISSLDHL
RVLRLRDTGISSIPD--FARL-QLKELDLSYNNLMRIPVALL---DSLKHLDLRENPLIC
ECSTIDFMHAAQRFGLVGYLDDPDALSCTFSDQ--SIALRKVHIQ---DCG-VINIFAIV
MASLIL--LAIVVVTYRRRRYIAYYFHVTAVRLKRYEPA---GEYEYDAFVGYST-ELNW
IINFLLPKMENENPYRLFLEERDMPAYGMQVSNIVAFMDKSHTVILVITQTFLTDVYCNF
MLKTAAM------RNNVIIIFLETIATEEFPAELRVLQLHSTCLHWSENRNSQERFWKAI
EYAMPQDPS
**>Pcau-TLRα3**
-ITEFADWAEIFLDGNFLDDFN------------------------------GLNLEKPL
TILSLSQNNIDEGGLSLKEILTHVTYLGLTENNLTQLPDKDFMLAASVQVISLNGNPFRC
DCETSYLKRWFSNNAQ--RINKPNETFCTSGPLYRRTAIADLP--DDVTCDSTFPYVYLS
LLVVALLAGVA---------GVGVYVC------RGDDGDESTG-KEYDAFIAFSSQDFEF
VARTLVPGLEGRPPYRLCVHDRDFHAGKLIMDSIIQAIEVSRSTVLLLSNHFIQSNWCKL
EFQASFIEVLANPRYKLIVIVCEPIEMDSLEPDLRFYIKTHTYLEIKD------KFWEKL
CAALPRPLA
**>Hsp-TLRα3**
-LTKVPYVP-LFMDGNDVNRLPNSVINGSFVGVSNLRILRLDGNLLQNLNGFEFLPLGNL
HELYLHDNLLEFVAKATFAALGHLKVLTLHNNRLHRIPSDLF---QHLTELTLSVNSWIC

DCENSTVQVWAESISD--IISDINLTYCLLGIT--GENLSEFNDS---LCMENFLYLILI
LVLIFIICVILAFLLYRFCYEFKVCLYRYGWRIN-----MEDYHKKYDAFVSYSTRDELF
VLEEFVHRLE--PQFKVALQYREF-PSSSVADGIMDGAHKSRRFVIFITENFLHYVWKEP
ESKSAHQQVLWDTRNQVIIVMLTERPDDKFEPDLRLYMKSKTCLRWND------MFWDKL
YYTMPDIKR
**>Hsp-TLRα4**
-VSDLKEFEFMFFDGNFLEAFF------------------------------NLNLTKPL
LVLSLMDNNLRADSLSLKDILLHVTHLSLSGNNMTELPDKKFMQKRDVENFIITGNPFR
CDCHTLYLKNWFQQNSD--VIVHANATFCKFGPYYQQTPIMDLP—DEVTCGATLPDV
QVNLATLLFAAPVT---------GVVVFLYNYRR—QSVKENE GVGIKEYDAFIAFNSNDFDL
VAYTLVPVLESNPPYRLCVHDRDFPAGKLIMDSIINAVEQSRATILVLSNNFIKSHWCKL
EFQASFIEVLGNPKYKLVVILCEDIPIDSLDADLRYYLKTHTYLELND------DFWPKL
IAALPPPMG
**>Hsp-TLRα1**
GLFNIPNLKVVDLCSNNLTRVP----QKLFHDVPTLESILLVNNSISFIHREDFKNLPAL
QLVNMSLNVIEGISPDGFIGVDNLNTLDFEANSLVSFAANNIGFLSKLREVDLQGNVMK
CGCFEISFSELLSSQ--RLTF---TDIHCVTPENLE--QV--F------HCPKSWLLYVCI
VG--VLFLFVILIIIVRFCSRVKIACHRYGIRIR-----QQPKGKTYDAYICYSRSNEHW
VSSTLVPVLESRPPYKLCVHNRDRPAGDSSSNSMVNAIKQSKVTVLVLSDDFMTSDW
CMVEFSPLHQSMSSY-TNNIIPIVLENIESRNVNTEMKRILRNKQALHVDD------YFWDKL
YYMLPDAEA
**>Hsp-TLRα2**
--TAVPEIS-LHLDSNDLSTIGNSEIAGSFRDLSKLTLLDLEGNSLTDVGSGVFTGLQSL
QRLHLSRNDISHVDERAFEGLSRLSALFLDGNALALPAAALY---ANASQFTLSGNPWIC
DCSMEFFFSWLKVNVD--RISDIGSTLCTV-----EMPIMDF---ETAYCSSAFIAGLAV
LAILFLLTVVGMAVVYRYQYEIRVWVFRYGIRRR---YPESDKNKLYDAFLSYHNGDEEM
ILKEFIPRLEYERKFLLCIHARDFVPGEFIAENITQAAENSRRTIVLLTKRYLESEWCRY
EFQAGHNQAICDQVNRIILVVFGDIPKDKLDSNLQAYINTNTYIRYDD------RFWDKL
LYAMPDPPI
**>Rva-TLRα**
-LTTIPMLP-VYLDGNRLPSLPNSRINHTFNGLSQLRNLHLHHNQITILRGGEFSQLVSL
EVLDLSWNDIHSIHEHTFLTLTKLRVLNLAGNQLDSLITLPL---PTVSQLFLANNVWEC
WCNEERLTEWLVRFTA--RIQDIHHMHCYDRSQ--AL-LRDMPRE---RCSASFVVVGIV
LGCVCFLVIVLVAFVLRYRYEIQVRLYRFRLRLS-----EEDYEKICDAFISYSDLDEHL
VLGELAPRLEFSPKYKLFLHYRDHPLGMRTPESIIQGVQLSKRTILVLSENYLKREWAKL
DFKTAHQQVFKDKKNKIIIVLLGDIQMKDLDVDLRIYLKQNPCLQWGE------LFWKKL
YYALPDPEP
**>Hex-TLRα**
-LTSIPMLP-VYLDGNVLPSMPNSRINHTFNGLSHLKVLHLHHNQITVLRGHEFDQLVNL
EVLDLSWNDIHSIHPATFSQLTKLRVLNIAGNQLDSLVALPL---PSLTQLFLANNLWEC
WCNDDRLTDWLVRHSG--KILDIHHLHCYDRSQ--AL-LRDMPRE---RCSASFIMVGVI
LGCVCFVVILAIALILRYRYEIQVRFYRFRMRLS---SQDEDYEKMFDAFISYSDQDEHL
VLGELAPRLEYVPKYKLFLHSRDYPLGTRTPDSVIQGVQMSKRTILFLSENYLKREWSK
LDFKTAHQQVFKDKRNKIIIVLLGDIQMKDLDVDLRIYLKQNPCLHWGE------LFWKKL
YYALPDPEP
**>Pcap-TLRβ**

MFSNF-SLKELYLGDNKLGPAFEGNLGHLFDNLTVLSLLDLSFNDIDIFSIDQFSSLSAL
KVLNLNHNKVSIFPPDVFDGLKSLERLNIKANKITVLEAGSFQLMKKLKEIDFSENPLQC
QCDVMDFFHWINFT--NLTITHWNDYFC--PQR--NTSLKEFLIMEAEECLHNIVIICAI
TISSLVIFVLLCVLAFRL-YRFIYVRASVEVNTQKSTTIKKNKVKCYDAFICYTSKDADW
IPALFKEHLGEARKLRLYFHDNHKHIERTTSWDVMNKVDSSYKVVFISTKNFVQTDWF
QWESMMLMF-----QDCAIIVGLEDIPTMNMSYTLQWLVRTKPFITWPVLDTDIGLFWDDL
AIYIKER--

**>Lloa-TLRα**

-LEPSPDIP-IYLEHMEIPVVRHSEIPLAFNTLPSLQLLDLSGNYLMRLTGDELYRTNKI
TTLLLHNNHLMSLGDRLNEVMPQLKTITLHNNKLQDLPLSIEQ--KQITDITLGSNLFRC
DCSPRFIQYWFSSNLD--MIHDVSDIFCVENISNFGDDIFKIP--MTQIATASFLIITAL
LAVALITIGLICLAVLFLRKTKSVIVQRYKVPPF--GTHTTGSSPLFDAFISYSKKDEKL
IIDTLYRQLES-EEYILCLLHRDSPNYSTISDELINQMECAQSLILVLTQHFLNNEWKTL
QIKTSHQIFAKNRHKKLIALLGDGIEPNQLDAELGQILRKNTCIRMND------LFWNLL
HSALPVRIA

**>Ovo-TLRα**

-LKPSPDIP-IYLENMEIPIVRYSEVPLAFNTLPSLQLLDLSGNYLMRLTGDELYRTNKI
TTLMLHNNHLMSLGDRLNEVMPQLKTITLHNNKLQDLPLSIEQ--KQITNITLGSNLFRC
DCSPRFIQFWFSRNLD--VIHDMSDIFCVENISNFGEDIFKIP--MAQIATASFLIITVL
FAVALITTGLILLAMLFLRKTKSVIVQRYKVPPF--GTHTTGSSPLFDAFISYSKKDEKL
IIDTLYRQLES-EEYILCLLHRDGPNYNTISDELINQMECAQSLILVLTQHFLDNEWKTL
QIKTSHQIFAKNRHKKLIALLGDGIEPNQLDAELGQILRKNTCIRMND------LFWNLL
HSALPVRIA

**>Ael-TLRα1**

-LEH------LLLTNTSIDRVP-------------------------------------LAQLSLGRNAITEIGVETFAN
ASRLSFINLSHNGARSFRKNVTDSLESSTEIK-ADSSLECECLG--EREWLGSSAE----
TSVTNLHCLAPQINGGEKRHRLQV-NQEIADAWTIVLIAIFSILLVFIIVAIIVILRFKVE
IQAFVFNFGVRIK---LPNDVGDKIYDAFLIFSADDEDWVVNTLLQKLETAPPYR
ICIHYRDFVPGNPIIQNVMDSVANSKSTLAVISDGFINSQWCKYEFVTAFQQTLKNAAHK
LCAILTQKIEPQLLKSNLQFYLKTNTYLEKSD------MFWEKLFFSLPDP--

**>Ael-TLRα2**

-LEH------LLLTNTSIDRVP-------------------------------------LAQLSLGRNAITEIGVETF
ANASRLSFINLSHNGARSFRKNVTDSLESSTEIK-ADSSLECECLG--ERE WLGSSAE----
TSVTNLHCLAPQINGGEKRHRLQV-NQEIADAWTIVLIAIFSILLVFIIVAIIVILRFKVEIQ
AFVFNFGVRIK---LPNDVGDKIYDAFLIFSADDEDWVVNTLLQKLETAPPYRICIH
YRDFVPGNPIIQNVMDSVANSKSTLAVISDGFINSQWCKYEFVTAFQQTLKNAAHKLCA
ILTQKIEPQLLKSNLQF---------------------------------

**>Isc-TLR3**

-LTQLPTLP-LDLSGNKLESLDSN----------------------TA---GLAKKAPFL
RLLNLSDNLLSSIDPSEIPQ--GTDELFLRGNRLSRFPIDLVSKF-NMSILELAGNPWSC
DCEDYAFRQWAEAYTD--VVEDAEEITCAKGPN--LKRFMDL---GQKLCPS--LSYGLP
LLVLLIISLAASTAYLRHKRAIKVWLYRGVCSS-CIKEDDLDEDKIFDVFLSFSSKDSMW
AYEQLIPGVEA-
HGFSVCTYDRNFKGGFLLQDIIHEAVSCSRRTLLLLTKNFVESEWCRW
EFRVAHHQALEDKINRLILVLVDELAPGLVDEELQLYMQATNYLRWGE------HFWDKL
IYSLPKKDA

**>Isc-TLR2**
-FTQLV-LRKLMLRNNRI------YIDGTFRNNGNLKYLDLAENRIEWLGKRAFSGLVNL
DLLSVSDNFLLHLNGSV-SHMPQLRILNFSHNA--------------IQ--TLYGNDFYN
DPELT-FYAYG-NNLS--TI---GAFQ-TSPKL----RM--F------ACA---------
--------LTASTAYLKYKREIKVWLYRGLCSR-CIKEDDLDDDKLFDVFLSFSSKDSNW
AYNELIPKIES-HGFSICTYDRNFKGGYLVQDIIHEAVACSRRILLLLTENFVESEWCRW
EFRVAHHRALEDNTNRLIVVLVDEVTSDAVDEELRRYMQVTNFLRWGE------HFWDKL
LYSLPKKDS
**>Isc-TLR1**
-LTDLPTSP-LYLQSNSISSLV-------------------------------APRWENL
TEVYLDENLLSNLDLTT---MRRLQILSLTNNRLRSLTPQLMGMLSSL-SLSLSGNPWIC
DCSTFSFKTWLRGHVY--MVKDYPDIACGD-----GVRINEI---PDSYCPVKQLAAVTA
ICVLAVLLVVVSVLYYRNRQTIIAYVYHFHNVFE-----DLDEDKTYDAFVSYSSADRDI
AMG-LLNSLESEEMFKLCIHERDWLPGYNISWNIVNSVQNSRRTILVVSKDFLE
SVWFQVEFHTAYYQMLEDRVDRLIVIVRGELPAETLDKELKFLLTTKTYLVWGE------
WFWEKLKYAMPHRRQ
**>Isc-TLR5**
-YGAIPRVP-LYLDGNDMSHLSNSTINRTFNGLVGLQVLHLDHNKVTALHGFEFENLTNL
RELHLSHNRLATVSNRTFVSLKSLTILYLDNNYIVEFQVWNF---PSLSDLRLGHNPWSC
GCRFMEFQDWVHMFGA--PLKDSVAIRCRQNQT--GP-LLEF---NATACT-NYMPLLIV
LPSVVVLLLFVLVLVLVYRKQMKVWVHKYGVRLR---QYAPEVDRLFDAFVSYCK
KDEAFVAQILAPELECHPPFRLCLRYRDLPMSGYVAEAITEAVECSHRTLVVLSEQFLK
SEWCRFELKTAHHELRCNSRHRLVVVLLDDVAVKEMDADARQCLRSAVLLRWGD-----
-RFWEKLRYALPDAAR
**>Isc-TLR4**
-HIAVPQLP-LYLDGNDIPALSSSTVNRTFSGLRTLRVLRLERNRLATLHGYEFDGLGEL
KELYLSYNHLTHVNNATFVPLKSLEVLHLDHNYILEMAIWNL---PRLNDVRLADNPWSC
DCHFAQFTDFLQNKGA-ELVRDLFSIQCVHNET--ALPLWEL---NTTSCT-DLVPLLVV
LAALFLLLVCIVVLAFVYRRHLSVWFYKYGVRMR-----PAEEEKLFDAFVSYSKKDEAF
VAQILAPELECQPPYRLCLHYRDLMAGGYLTDAITEAVESSRRTIVILSEHFLKSEWCRY
EFKSAHHEVLHSCTHRLVVIFLGRVSYKELDPDIRLWLKSSTFLRWGE------RFWDKL
RYAMPDTRH
**>Dpu-TLRβ**
TFYGLNSLEYLNMDRCKL----TD--EAIFAGAPRLRHLSMRDNQIVSFGSNPFADATSL
VSVDLFKNRIRGWDTQLFAGSPDLDVLNLAENQISTVSKAMMADIANLSEVDLLGNPID
CDCNLEPLRRYLEDTEDNLLI---KADHCSSPDKWRFQPITSFDPD---HCYYSFVIALYI
LIPTVCLSMLVGYAIYRSRWVIRYYMFRKRLSQ-SSSSMAEEGNFKYDAFVSYS
NVDHAFVAR-MVGMLENPPHYKLCVYERDFTAGNVLNDCIMQSIATSRKVVLVISE
NFIQSHWCLWELHLAQHSLLEDKRNGLVLVVVGKLKLNQCPPTLRFLMKTRIYLEWDL
DPSKQRVFWERLRDALAQKPD
**>Dpu-TLRα3**
-LNEIPRLPRMNLSSNSI------QIPNSS-----------------------DCYPDVTWLDLSHNGMDESS
MSDWQNLPKLNRLDLTHNNFNSIPNGVVDSWHNL-TYNLNGNPWKCDCTNLALL
NFIYGSWK--RLEDFNQMKCDN-----GQKISEL---SVELCP-SVKYYTIPLPILALLIVCVGIIV
YRNRRVIRAWLYRQLCLWK---EEEENDERIYDAFISFSHHDEIFVNEVLVPQLERPPHY

QLCIHYRDWLAGEWIADQIVRSVATSKRTIVVLTENFLDSLWGKLEFRTAYKQVLTDKR
MRLIIIVKGELPPDKMDQELQTYLSLNTYLKYDD------FFMDRLRYALPHNTS

**>Dpu-TLRα2**

PLNDLPGIP-LYLDGNNLTELSGSTLNRTFHGLGALQVLQLADNELEELRGSEFEPLDHL
RELYLQNNKLRFISDTAFVHLRSLQVLRLDGNRLLTFPLWRL---PHLNQLSLGLNPWSC
ECRFLAFQQWIAAHPQ--QLVDSDSLHCLMGDQ----QLIGF---EFNSCSADYLPVMAA
GICLFLLGLIAVVLVFVYRQTVRIWIFRYRIRLS-----EEDKDAMFDAFVSYSLKDEQF
VSQVLAAELEHESSFRLCLQHRDFPTSHPGGDPLTLGLAASRRIVLVISQSFIESEWTR
PEVRTALTGFLRLPRSRLVAVLLTPWTDDQSDPELSLLLRSSIIIRWGE------NFWSKI
RYYLPDPTP

**>Dpu-TLRα1**

-FDDLN-LKTLWLDSNKL------KIGKIFKNIPQLISLQLGSNVIKQLEIGAFSNLPNL
FQLNLQNNQLDILPSDVLQSLANLKYLDLSNNKLTIID-------RNL-TYSLSGNPWRC
DCSNLALLKFIYGSWK--RVEDFNQMRCDN-----GLFFFEL---SVELCP-SLKYLTIA
MPVLALLVFCICTIFYRSRRVIRAWLYHQFCLWQ---EEEENDDRIYDAFISFSHNDEKF
V-DELVAQLERPPNYQLCLHHRDWLAGEWIPDQIVRSVASSKRTVVILTENFLDSF
WGKLEFRTAYQQVLKDKRMRLIVIVKGELPPDKMDTELQTYLSLNTYLKYDD------
FFMERLRYALPHKKN

**>Dpu-TLRα4**

-HPNLPRIP-LYLDGAQLRALSSSIINRTFNGLRGLYVLHLEDNRIRTLEGFEFSDLESL
RELYLHNNAITSIQNRTFSALKHLQVLRLDGNRLVDFPVWNL---PELNALTLNDNPWSC
DCLFLALRTALHTAGP--KVSDASQLICGGSNR--NRSL---------LCV-DYLPLLVT
TLVAFIAVTLIILFVFIYRQPVRVWCHRYGLRLS---SAATPDSKLFDAFLSYSAKDDAF
VQQMLATNLEYSPTYKLCLQHRDCPSGGGLSETISQAVDSSRRTVMIISPNFIKAEWC
RFEYKSALHQLFGTSRKRLIVILIGDVTHKDLDADLKLYLKTNTYLQWGE------GFWDKL
RFALPDPVQ

**>Ppr-TLR1**

---------------------------------------------------------------------------------------------------KSHIDSL
SSLEYLDVRDNEFACSCDLRWFQTWMTQVP-KMIIPNKNSLHCRSPTDMTQDSVA
NYTTPWIK-CDNFIVVGGVSV--FVVLAIVITLLVGLKRWEIKYWWVFKKARM-GWRK
LRDS---EFDAYVCYHSEDEEWVTQTLQENIEGNVNFKLCIEERDFILGRQHLENF
TDLLNKSHKVLIVVSQNYLKSLWCRFEVGMAQLKLYEDNRDLLIFILLDNLKRKDMPRA
LKCLMLNSRVLPWPKSSQMKSVFWMKLKLALQE---

**>Ppr-TLR2**

ILTKLPNLSKLDLSSNELGS--SSILPYLFKNLSTLRELDISDNILRTMHEDMFNSLLHL
EKLVIRHNYFKTLPTNIFKRLVNLRSLVMSTNNLCGLYKDQISPLISL-RLDVRRNSFYC
SCSLRWFQNWLQTT--RVWVDDKYKMRCNSPSDQWSNTLINFTIPWYR-
CDNTIMASTSV--SVLFLIISIVILLIFFRWDIKYWWVFSKVSIRGWHHF--
SEEKEYDAFVCYEHSDEDWVMKELLENVEKNNTFKLCIHERDFIPGKRIVDNIERGINL
SHKVIIIVSSAYLSSQWCEFELDMAHIKLTKKKK----------V------------------------------------
---------

**>Ppr-TLR3**

----------LLLGMNNLTGFINSETPPLFDKNSYLKSLDLATNGLHVVHEKTMKSLPNL
QYLNLSDNFLSDIS---ISGMLQLIVLNISKNNWKDTPTELIQSLQKLNCLDISSNPLTT
ECEIQEFIQWSLYT--NVTLTKYNKFECILKNN----HV--FSETVIKTCNSGMFLAIGL
CTSFFILSVIVLTITYRNRWTIKWWLFLARKYLRLREELAEQRNYQYDAFVAFSADDITW

LKSDLIPELEMDRGLKLCIHHRDFQLGVPIEENIVNAIANSRKTILLITNKFVHSNWCMF
EVHMARQRLFDEGKNVIIAILLEEVNIGKLNRTLRNILTSNTYLEYPKNEDGQQLFWIKL
VDALRSNKD
**>Ppr-TLR4**
LFSCLKNLATLLLQGNNLGGVITDLHGDLFSSKSNLVDLHLDDNNVETLPVNLFKNATS
LKRLSLSKNTIRHWHERLFRKTTSLEYLNLAHNQISLMNKTSLPNLNILKTLNLTANPFA
CTCDLIWFHDWVQNT--KVNLPGVEDYTCDSPQIFQGVPLKEFDPNKLV-CWEKLIMY
ISFPS-LIAVILVSFLIVYKKRWSLRRYWFIMKMRARRKRLMEE—GLEFDAFISYCTA
DKDWVEQTILSKLDKKA-FKFYYDARDDIPGKGIYDNLQYGFEHSRKILIILSTEYFEDK
HADLELQLIPEIEVDAREDKVIFVFKEDVYVNKIPRSIRRKVDNDDFLTWTDDQAVQDLF
WGRLHEELSK-PQ
**>Cs_Toll4**
SLIDMDNLTTLLLQNNHLGDSLRDVIGQTFSSQKKLKVLNLSKNSIKALPYLIFKNQVSL
KNLSLARNAMTDVS-FSLKTMKMLQFLDLSDNQIEYVTSENMGYLDQIAH
LNLSGNILACMCDNQQFLSWIATT--KVHIIDRGQLKCLYRNKT-TLSLSRIIRSQLKDC
SLWIVMTSCVTGFGLLLILSLITLLYHRRWQLRYLWYIGRKKIDPFH--HDSRLPQIDVY
ISYEQHDVQTVTDYVYPFFERRG-YIVKIRE-EFEATDKLYRVIPDTVNKTRKVV
VFLTPSYCKDYWNTFEFNIAAYEGIYTKRNIIIPVLIGDFSDQNFTPEIRSFVNSKIVLRFP
SQAHRINTFNEQLEHWLQ----
**>Cs_Toll13**
TFKYLPKLIALEVSYFNMFDMPAQSINTLFSPLSNLKYLMCYNCQIRDDPKLFLSNKSQL
NRIKLDRNYIENISNDTFKSNPLLKTLSINMNKIGHLKASEIDFLNSLDSLDLSHNPFIC
DCDLEWFITWLKST--KTAKVEYQNYVCAYPANMAGTKLTDVHYTYRE-
CHPVWEWVGIVGGPIAVVLAIVSFVLYRKRWSIKHYIYLMRKRR-NYILV-
DGENFLYDAFVAYNQEDSDWVREHLLPVLEDEHQLKLCIHERDFRAGILINDNIVTCIEQ
SKKIIIILSNEFAKSGWCMFELRVAHSKHIEDE-MELVVILLERINGRNMNN
SMKTLLETTTYIEWTEDQHGQLLFWNQLKASMNK---
**>Lrug-TLR2**
-FLRIPELQK-----------------------------------------WRYDIPKLIYLDLSYNDIDRIDINGFPDDG-
LGRINLQFNNITTIRSEDIRA-LEA-IVDISNNPINCGCG-IKLYEF------LGNYEYIR
DLVCHSP--LKGRKIRNLT--QKEICPTH-MVLIVSLCAVVVILGIIIIILL
RYRREVVILVYRLHI---PCQPVDTYDSKNYDAFISYSSKDDDWVLRTLVQRLENEE
KFKLCVHHRDFEIGAAIADNVVQSVEDSRHTVMVLSRNYVDSEWCIY
EFRTALHQSLIERQKHLIVVILEDVPKSELDPDLRKCLETFTYIQVGD------LFWDKI
RYSLG--HR
**>Ppe-TLR1**
VFKNLSSLKTLNLSKNRLKYMVEDKYGGMFEGLDNLVTLNLENNDIEGLSPVIFLHLTN
VQNLLMSGNRMAHWDKELFTNTQHIKNLDLSRNKISVIDEHALNNYSNFKTLNLEDNPF
ACNCELVWFCKWANRT--KVTLVKFNNYTCSSPKSRQGVLLINFDWRSLV-CFNPYI
IAGSVVG-GVFLMTLIVVMLYRCRWRISLCCYRCGQRC-DYHYL-E-
EDKRFDAYISFDKKDNSFVQEVIMNQFDRNGKYQLCFEPRDFRLGSSIVGSMCVAVEN
SHRAIIVFTNAYLASGRFQMELDLLHNEHLDRSFGMIFVTTGPQLDFRLLPKWLHKSYE
DGKFLVWDENSSAQEEFRQRLDRKLRTPPP
**>Ppe-TLR3**
VFQRLNSLKHLDLSLNQLGRLTQLQFVRLFQPIRTLEYLHLGHNDLTFLNAPIFEDMLAL
KTINLVENSLKVVQSNLFESSPSLERVLLSDNQISFLDAGMFRRMTNLSFLSIEENEFEC

DCALRQFRDWGHGD--GVMILGLHGERCFASDKRLDSRVTEYETEWIE-
CDHEYIIAGALGL-FFAFATLLAGLVYRYRYDLLWWLLKRRRRR----PT--
TAGERYHAFVSYNSRDSRFTLS-MIRYLEDDIRFKISFDG--FDPASFISDCI
VQCIERSEKIIFVVSRTFLQSEWCSYELRMGELKCFEERRNIMILIFLEKIPVKELPRSLR
TLVRQINYLQWPVDERARDVFWKRLKIALSKDAK

**>Pps-TLR3**

YFANLTWLEHLNLEGVNLRDVNVDTPDCLFEGLDNLKVLDLTNTNLKGIPTRFFKDLRS
LQELILRQNRLSGWNDVVFDNLTALQFLDVAVNQIRVVNMSSLQSVKSLNRFQAFSNP
YICDCNLAWYADWLRRMHATLKVTYQLPYNCSN---LKRVSVLSYRPTFFE-CHRLIIL
SASGGG-LFILVMLTVTVLYQYRWYIRYWMFLLRSRRAKHVEEADRLIYKYDGF
VTYSGEDSEWVIRTLLPKLEKEYGFSMCIHERDFTLGRDISENIAESIEQSRKVLVVLTN
NFVRSYWCKFEVNLAHANTLHNSRQSLIIILAEDVDMDLMTPILRYLIRRKTYIEWTMND
QGQILFWKRLKEAMQKRGN

**> Pps-TLR2**

ILKNMTRLKTLILSDNMLSDSLVDESGRMFKDMNSVETLDLSGNRIFILHVNTFKHLINV
KTILLKDNRLASTPAVNVESLESLEQVDLSSNRIQYLSNEFLQSVTK--SVRLTGNPFNC
TCEILVLLQWLAGSEDAMKLVDNTTLACSGDNIIRGAKLVDFHYKSLQRCI-LWIWLSVS
TILTISILTFCIGLAYRKRWSLRFWIIAAR---RKYERL-PSTNYTYDAFVCHSSFDAKW
M-NTLQKELEQEPNFKLCLHYRDFPLGLPIVECINDAIVNSKYIILLITRNFIESQWCIY
EFYMAKTRVFCENRSRLIIVILEHLPEAVLPQTLQNVMKDHVYLEWTDDPVGQQAFWE
RLRENLAAEPP

**>Pps-TLR4**

-LTDFPTLPDLNVSNNDLTQLP-------------------------------IYLTQLVELDATGNNI
GDISGTALLQMSNIKALYIRNNNLRKLPKALLDSHGNASVLTLGENPWDCSCPNEW
FLKWMTSRGS--VVTDVGDVTCDIP--VRGQRFSDDV--IKQNCETDYMVMAST
VGSITALLIIALVLVVIFRHDIKVILFKWDIDLR--A-EEQSKDRPFDVFVSYSSLDGEW
VRQKLLPMLERNPPFRTCFHERDFLPGAPIAENIMRAIQASKRTLMVVSKNFIASDWCE
FEFLTAHKSFMETKQNKIIVVMLEDVDTKSMDPMLRAYFTTKTFIRAAD------LFKEKL
YYAMPR-TE

**>Pva-TLR1**

-------------------------------------------------------------------L-----------------DLSHNSLGYMESS
VFQNLSGLKFLNISKNKFKCDCDLRPFRDLLHEA--TFHF---GQNPCYYPTSLKKQL
VSNYSLSFIA-CDHEMIIVLAVAG-FIFLTIPIPALIAYYRLNLKYWWYFGRRRA-GYRPL-
DGGVHRYDAFVCYSKNELSWVVRELVEELENNERFQLCIHDRDFDLGGDIVDNIIRSID
CSRRVIFILSREFIRSYWGTFELNLALMEAIEKRINFIILIFFENIPKKEIPRHLQCFMRHVT
YASWPQQARAREMFWMKLKLALRNREE

**>Pva-TLR2**

YFANLTSLDHLILNGVYLRALNDDKPECLFQGLGKLKVLDLAFTHLKGLPERLFKDLTRL
ETLILRHNQLSGWNDVVLENLKNLRSLDVSGNQIHTINQSSLRPVSTLNHFQAFSNPYI
CDCNLAWYADWVRQMHATLKISYQVPYNCSN---LKNKSLLQYRPTFVE-
CHRLIILSASGFG-LFIVILAVIGLLYKYRWYIRYWIFLLRSRRARHIEENDRLL
YTYDCFITYSGEDSDLVTQQLLPKLENEFGYKMCIHERDFKLGRDISENIAESIEKSRKV
LVVLTQNFVQSEWCKFEVNLAHANTLHNARQSLIIILVEDVGFEHMTPILRYLIKKKTFLE
WSNEEQGQRVFWERLKDAIQQRGQ

**> Pva-TLR3**

YFAGLLTLISLKMSDLDMGKLS--DKVCLFEGLTELKYLDLHKVMLKNLPANIFVDLKSL

VYLRISGNKLLAINPVVFSSMLSLERLDVSSNLIQNINQSSLQPFKGLNYFEAYNNPYAC
TCDLQWYTDWLRQMKRKIIRKNMQYKCASPKKFKKKNLLTYNPTFID-CYEPFIIGTTV
GS-FAVLATIVVAVGYHYRWYIRYWLFLFRSRFAKNLREDERLVYRYDCFVTYCE-
DDGWVLETLRPKLEDEFGFRVCLQDRDFELGKSKVDNIDEAIQNSRKVLIFLTANFAMN
SWCNFELSLAHANCLENDQQHLIIIMMEDVSPKYMTPILRYLVRKRTYIEWTGDEVGQN
LFWQKLPDAIRSPNQ

**> Pva-TLR4**

YFSGRPSLVSLKMPGVNLHKQT—KSSCLFRGLFRLKHLDLHDVQLKRLPSDMF
QDLQSLVYLRLSGNMLHEINPVVFSML-SLARLDVSSNQIQNINQSSLQPFKGLNQF
EAHYNPYACTCDLQWYTEWLRTMIRKVTIRKYMQYKCATPKQIERKNLLTYDPTFLD-
CYEPFIIGTSVGS-FAVLVIIVVTVGYHYRWYIRYWLFLFRSRFAKNLREDERLVY
RYDCFVTYCE-DDGWVLETLRPKLEDEFGFRVCLQDRDFELGKSKVDNIDEAIQ
NSRKVLIFLTANFAMNSWCNFELSLAHANCLENDQQHLIIIMMEDVSPKYMTPILRYLVR
KRTYIEWTGDEVGQNLFWQKLPDAIRSPNQ

**>Pva-TLR5**

RYKIAPSLKKLILSGNKLYIMLRDQKKKHFKHLNNLTHIDLSHNSINELYPEVFSELSNV
KEIILRRNRLLAVTSMNITEMPSLKNVSLVANNIRVVPETSLRAWRG-KSFDFSRNPFNC
SCQFLPFLRWFNNVSSTVTLLHAENYRCGDEQKI---FVREVRLDELVKCTSAWIVISIA
LSICVALCITLGSILYRHRWTIRYYIVSAARRTRGHPPQ-MLRRFQYDAFVVYSSEDRHW
VHDAMRTRLEDGSDFGLCIHYRNFLPGRHIEESVIDAIENSRHSILVVSRNFLRSEWCIF
EMHMARNIFRQQRKDVLIVVLLEDIPVQEAPLTLVNLLRTRTYLKWPADDVGQEAFWE
MLKETLKKEPE

**> Pva-TLR6**

LLKGLGNLKWLFLNNNQLGKNLLQEYSLLFGDTLSLKELHLDRNDISSLPGNLFHSMKN
LEILSLRDNKISHWSPKLFAPLKSMEALDLSNNLIALINQTSVHNING--AFNLTGNPFAC
TCDLMWFRQWVNIT—NISFPCIGQYACNSPHSLQNTKFLDWYPDPRD-CINPFYVG
GSVCG-TVLLMLVISGATYRRRWFIRLSWYKLTHRRRGYRSLNNADVPSFDAFVSFCE
EDRQWVFDTLMKTFDEDNNFNICHDERDFPPNLSTAGCIFGCIENSRKFIVVVSEDYD
YCGRLEIELHYALQEIMEDAEFEIIVLLKDNPHPSRIPKHVAHLVSDPEFVEWPSDNDGQ
QLCLRRLQTMLERD--

**>Ce_Toll1**

-MVPVVELP-IILSGVTLPQLRGTSIPKAFHTLPALKTLDLSDNSLISLSGEEFLKCGEV
SQLFLNGNRFSTLSRGIFEKLPNLKYLTLHNNSLEDIPQVL----TALSKISLSSNPLRC
DCSGEHAAEWFSLHRH--LVVDFPKVECWENVTNMGNDVFVMP--IEELRDYSILFVIIT
ISIAVLLCVLVILAISFIRKSHDAINQRYKA--S--NCSTSGSSPLYHAFVSYSKKDEKM
VIDQLCRPLED-EDYQLCLLHRDGPTYCAISDELIAQMDSSQCLILVLTKHFLENEWKTL
QIKTSHQLFAKNRAKRVIAVLGDGVDANLLDDELGQILRKHTRIEMRS------LFWTLL
HSSLPSRLP

**>Dm_Toll1**

-LTHVP-LPNLHLENNTL----------------------------LRLPSANTPGYESVTSLHLAGN
NLTSIDVDQ---LTNLTHLDISWNHLQMLNATVLGFLMKWRSVKLSGNPWMC
DCTAKPLLLFTQDNFE--RIGDRNEMMCVNAEM--PTRMVEL---STNICPAVFIALAVV
IALTGLLAGFTAALYYKFQTEIKIWLYHNLLLW----EEDLDKDKKFDAFISYSHKDQSF
IEDYLVPQLEHPQKFQLCVHERDWLVGGHIPENIMRSVADSRRTIIVLSQNFIKSEWAR
LEFRAAHRSALNEGRSRIIVIIYSDIDVEKLDEELKAYLKMNTYLKWGD------WFWDKL
RFALPHRRP

**>Dm_Toll2**
-LAALPRIP-LYLDGNNMPELEASTLNGSLAQLVNLRVLHLENNKLTALEGTEFRSLGLL
RELYLHNNMLTHISNATFEPLVSLEVLRLDNNRLSSLP-------HSLQGLTLGRNAWSC
RCQQLRLAQFVSDNAM--VVRDAHDIYCLDAGI----K-RELELIANGDCS-YRLPLLAA
VL-VLIFLVVVLIIVFVFRESVRMWLFHYGVRVP-----FEDAGKLYDAIILHSEKDYEF
VCRNIAAELEHRPPFRLCIQQRDLPPQA-SHLQLVEGARASRKIILVLTRNLLATEWNRI
EFRNAFHESLRGLAQKLVIIEETSVSAEAEDAELSPYLKSVPSLLTCD------YFWEKL
RYAIPIESP
**>Dm_Toll3**
-LLQMPSLSS----------------------------------------------------s-----------------------RVTYVDLRNNNL
TALSQKNRSSINRL-KLHLLDNPWSCSCNDIEKINFMKSVSS--SIVDFTEIKC-SN----
GEKLVSIN--QHI-CP-SDLFYYLALAISLVATIIALNFLIWFRQPVLVWFYHGVCLSA----
RELDKDKRFDAFLAFTHKDEALL-EEFVDRLERRPRFQLCFYLRDWLAGESIPDCIG
QSIKDSRRIIVLMTENFMNSTWGRLEFRLALHATSRDRCKRLIVVLYPNVKNDSLDSEL
RTYMAFNTYLERSH------NFWNKLIYSMP----
**>Dm_Toll4**
-LSEIPQLPTLVFERNSLKKWP---------------------------------PGYSSV
TRFYLAHNRLSDIDQ------DKLEYLDISNNNFSALDDRVRGFLKRL-QLSLFGNPWTC
RCEDKDFLVFVKEQAK--NIANASAIQCIDT----GRSLIEVE--ETD-CP-SVLIYYTS
LAVSLLIIALSINVFICFRQPIMIWFYHEICLSA----RELDEDKKYDAFLSFTHKDEDL
I-EEFVDRLENRHKFRLCFYLRDWLVGESIPDCINQSVKGSRRIIILMTKNFLKSTWGRL
EFRLALHATSRDRCKRLIVVLYPDVEHDDLDSELRAYMVLNTYLDRNN------NFWNKL
MYSMPHASH
**>Dm_Toll5**
-LEELP-LPRLKVGNNSL----------------------------TSLPTSEHSGYANV
SGLFLSDNNLTSLGS---DQLPNLTHLDVRGNQIQSLSDEFLLFLNNTMTLSLSGNPITC
GCESLSLLFFVRTNPQ--RVRDIADIVCTKQKK----SFQQM---EAFLCP-SYLLISCV
VGGLVIVICLLTVFYLMFQQELKIWLYNNLCLW----EEELDKDKTYDAFISYSHKDEEL
I-SKLLPKLESPHPFRLCLHDRDWLVGDCIPEQIVRTVDDSKRVIIVLSQHFIDSVWARM
EFRIAYQATLQDKRKRIIIILYRELEHNGIDSELRAYLKLNTYLKWGD------LFWSKL
YYAMPHNRR
**>Dm_Toll6**
-YSEMPRVP-LYIDGNNFVELANSHINTTFSGLKRLLILHLEDNHIISLEGNEFHNLENL
RELYLQSNKIASIANGSFQMLRKLEVLRLDGNRLMHFEVWQL---PYLVEISLADNQWSC
ECGYLAFRNYLGQSSE--KIIDASRVSCIYNNA--SV-LRE----KNGKCT-GLLPLLLV
ATCAFVAFFGLIFGLFCYRHELKIWAHTNCLMNK---VDQLDKERPNDAYFAYSLQDEHF
VNQILAQTLENDIGYRLCLHYRDVNINAYITDALIEAAESAKQFVLVLSKNFLYNEWSRF
EYKSALHELVKR-RKRVVFILYGDLPQRDIDMDMRHYLRTSTCIEWDD------KFWQKL
RLALPLPNG
**>Dm_Toll7**
-TTELPRVP-VYLDGNNFPVLKGSAINRTFASLASLQLLHLADNKLRTLHGYEFEQLSAL
RELYLQNNQLTTIENATLAPLAALELIRIDGNRLVTLPIWQMHATTRLKSISLGRNQWSC
RCQFLQLTSYVADNAL--IVQDAQDIYCMAASSSGSLK-RELDFNATGACT-SYIPLLAA
AL-ALLFLLVVIAMVFAFRESLRIWLFHYGVRVP-----CEESEKLYDAVLLHSAKDSEF

VCQHLAAQLETRPPLRVCLQHRDLAHDA-THYQLLEATRVSRRVVILLTRNFLQ
TEWARCELRRSVHDALRGRPQKLVIIEEPEVAFEESDIELLPYLKTSAVIRRSD------
HFWEKLRYALPVDYP
**>Dm_Toll8**
-YEQLPHIP-LYLDGNNFRELQHSVLNRTFYGLLELEVLQLQSNQLKALNGN
EFQGLDNLQELYLQHNAIATIDTLTFTHLYHLKILRLDHNAITSFAVWNF---
SYLNELRLASNPWTCSCEFIDLRDYI-NRHE--YVVDKLKMKCISGNSPASLPV--V------
QCSNDYIPILVAILTAFIFVMICISLVFIFRQEMRVWCHRFGVRLN---VDKNEREKLFDA
FVSYSSKDELFVNEELAPMLEMEHRYKLCLHQRDFPVGGYLPETIVQAIDSSRRTIMVV
SENFIKSEWCRFEFKSAHQSVLRDRRRRLIVIVLGEVPQKELDPDLRLYLKTNTYLQWG
D------LFWQKLRFALPDVSS
**>Dm_Toll9**
AFDGIATLKYLYFERSNIKDLE--------KSLKNLQVLGLAGNNINALTPAMFQSLESL
EILDLSSNHVGNWYRSAFHN-SALRVLNLRSNTINMLSNEMLKDFERLDYLSLG
DNDFICDCHLLWYIPWLQRSYSKLRFEDYMVAKCSAPYHLDGDTLLDFQLQVDENCQ
SELHVTNTVIAVMLVGACILGFIIYLKRWHIHYYYSSLKSAAKKFTNIQRDPSAVYDIFISY
CQNDRTWVLNELLPNVEETGDVSICLHERDFQIGVTILDNIISCMDRSYSLMLIISSKFLL
SHWCQFEMYLAQHRIFEVSKEHLILVFLEDIPRRKRPKTLQYLMDVKTYIKWPTAKEDR
KLFWKRLKRSLERE--
**>Ci_TLR1**
AFKHV-NLTC--IKFNQV----EQ-NGIMFSGL-MVKQLYFIRSNIRSISSSAFTGSVHL
RLLDVSYNKITGLEKDIFTNL--LEELNLRGNQIRVLDPSTFSSLVNLRSLDIENNRFLC
NCDIIPLQQWIIDKLYRILL---RNVTCSLHSSRSYVDIIEW---DSELCWK--KIVGIV
LGC-LLLSTACAVFGFSVRFQALFWYEMIKSKV-SYHPRNRSDVYEYQAYISC
DSVDEAWVVRQLLCAIENETPMKLCFPSRDFKPGCPKMVSAANNLRLSKHALVILSKD
YVANSWTRFELSMVSEMWRNSERESLIVVYLKEV--ERL---PVLGVRRNAWLV
WPTDVADRPSFWMKLRRSLAK---
**>Ci_TLR2**
RFNVLPTIPRLDLSNLQL----NE--KISLTQLTRLTTLNLTGNKLTSIPL--QGLPRSI
ENINLSRNKISTLPATTLTCLPNLKQLDLRNNSFSTIQTQEVSIFLAVTSVLLKGNPLEC
NCKLRPLITWIQTNEKDLSTHDLKDLICFTPKRFEGRFIINLSES----CP-NLLIGGLV
TALLIIIIIIVNIYLYKKRKKQERRDI-------GFKDLEED-TYEYDAFVSYSSDDVEF
VYK-MLEEMEEKRERKMCIHERDFTPGRGIADNIVECISTSRRMVLVVSRKYASSA
WCQYEVQIALTELHAKRRRLLVPILLEDVTREQYAGSVTTILSAITAIQAPKAQRTWANF
WNKLDKTLT----
**>Od_TLR**
QLKNL-NLRGVDFSMNKITHFC----IDDFVNLEDLEFFNASLNQITDIPNNTFSFAREL
RVLDLHANSIQELN---FANLPELRMLDVSENQIRTSVDPYL---GALEQLDASYNPFQC
DCQLKKFVQFVQEPGRIVGIQAQSRYKCQIPRLLGNLNLLRL---QDKVCENEFYLSILA
IS-IVVILVVVAVVSKNRRQRMKMKELSGRNRVR---AAKN-NIVKNDAAILCHINSQKW
VTDVMLPTLKQKPQEKLYI---DFIKSQVKNEKLRRCVEQNKRVIIIITTEFASSDACLF
CLQAIYDLTRRNRKDGIVLVVLEPIPWNSMPHALKILMAEKTFIQYPVEDVGRQYFWDA
LRASIYQERT
**>Sp_TLR020**
VFRQLSVLQELNLEYCQIGNL-----PLVFSGLESLQKLSLKGNNIQHIHDDVLSGLGQV
NIIDFEGNQIIYLDELIFSNNRNLTNLSLADNKLTRFNQKTFKPISSISSLDLSMNPIDC

NCDLKWLIYWINKP---IHLIDRDKTICSSLEPFREKPLLDVDPNEL--CILGLLFL-IP
LASIGL--VVISVLLYHFRWQLRYKLFLLKLAA-GYKEMRDHNDYEFDVNIIFGEDDEEW
IREQLRPALGERLQ-RNVFGDEDLVLGMHYLDSVHYVVSHSYKTIIVLSRAAVQDRWFIL
KFRTAMDHVSDTLTEFVVVVFLEDIPDDEMPFLARLYLNDGRYIHWTEDARGQECFWD
ELTKNLT----
**>Sp_TLR007**
VFQNLSQLVYLDMTNSRIHTLR----SGLFSPLSSLRYLYIGENNLGEVPGDIFNGLFRL
NVLTFQNNILSSLDPKTFAQTLRLTDLYLPGNQISTIKPGTV---NTS-RFDISKNPFSC
TCSLAWFRQWLDSAD--IDFKHADQTLCSGLKGLSKQPILSFHPD---HCGVIFLIAGIS
FTGIFL--FFITLLAYNRRWWLNHKLFLLKLAV-GYKEMAEADNYEFHLNLMFLEEEEEW
VDRVMKPALEERFHQNIIYGDKDLHLGMFYINAINDALDNSFKTVLLISNQSIRDAWCMT
KLRMALEHLNETGLDKIILIFLEDIEDENLPYLVRLFMSRNKYMLWTDDEDGQELFWAQ
FEKSMRAN--
**>Sp_TLR053**
IFTPLRNLVELDLTSCCIKQVA----SRTFANLTTLLQLSLQDNDLTSIPKDAFQGLQNL
QVLRLQNNLIKFIHQGLFMGTNELEQLYLQNNHISTVASNTF---SSL-RFNIAYNPLTC
DCQLAWFRQWLNEVEGKIDLAPKNQTRCSSLKVLVNQIIWSFHPD---YCGITMIIVSAC
FAPILV--LTLGILVYLNRWWINYKLYLLKLAI-GYHEITEPEDYEFQLNLMFHDDDEWW
VNDCMKPFLEQRMHERVIFGDADLHPGSFYLNAIYDVIENSHKTILLLSNQSVDDTWYM
TKLRMTVEHMNDTKLEKVILIFLEDIDDDHLPYLVRLLLSRNKYLLWTEDEEGQEVFWA
KVQKSMRQN--
**>Sp_TLR039**
ILTDLLLLQELDLSDCQLTEI-----VNAFEGLQSLQILHLEGNQLLDLPHGVLWNMAHL
RNVYLEGNKLKYLDRDLFFNSSRLRNLTLARNQLTGLNHSTFKPIKTLLSIDISENEITC
TCNLKWLPIWLSGS---ITLLNEIDTRCSSLEELELKPLMSFKPAEL--CGPIALYCSLP
IVTTWI--IIVLVFAYRHRWFLKYKLFLLKMAV-GYREIRDFDDYEFHLNVMFAEEDEGW
VRYRLRPVLEELLE-RNVYGDNDLPLGMHYYDAVHYVVEKSYKTIVLVSRAAIQDNWFII
QFRTAADQVNDTQIENMVVIFLEDIPDVELPFLVRLYLSDRKYLSWKEDERFQEYFWQ
KLIKMLKRN--
**>Sp_TLR056**
IFQNLSNLQILRLDKCSLSVL-----IGIFIDLKSLVSLHLENNHLKVISTGLFDKLYDL
QYLLLNGNELTYLDSNLFKYLSSLRCLDASENRISGLNHSTIEPL-RLTTLGLSLNPLVC
NCNLKWLPGWLKGT---IELIDSMGTTCNTLEPFRGKQLITFDPRYE--CGPITLYSCLA
MIGFVL--IFAVGLIYYQRWWVRYQLFLLKLCF-GYEEVHDRGEFQYDIAIMLDEIDNEW
VNQHLRPALMERGD-RIVCGDEELMLGMFYLDAVHYATEKSFKTIFVISHAA
LQDQWFMMKFRTVLDHVNDVGTEKMILVFVEDVEDDELPFLIRLFLSDHRYLVWPDDE
RGQEYFWEELIRDLTRH--
**>Sp_TLR044**
TFQGLQNLQNLEMDNSDITSLN----EDIFLNLTSLQHLSIDVNHIAELTSRHLADLRSL
VGVSIKSNEIKGLASDVFTNNPHLSYLYISHNHLTTVKEGTV-----LRTLDVSNNPFSC
NCEFTWFLNWINKA--EVSIIHPDQTNCSSLAPFKNQPILAFDPT---VCGPVWVYIITI
--FVIVTCIMICVVAYQRRWLINYKLFHLKILL-GRRDDHDR-DYEYDINLAFDDDDEQW
VRGILKPGLEERLDDRIVCGDDDLPLGMYYIEAITEVFEQSYKSILIVSNRAVDNHSFIS
KLRLAVDQMNEVELEKVILIFKEDIPDGRLPYLVRLFLSKNKYFRWSEDKYGQKIMWEK
LVRELGKD--
**>Sp_TLR016**

VFNNLSALQVLNMSDCQISTI-----SGAFASMTSLTILSLQNNDLQILPLHIFDNLIHL
SIFSIGNNVLVYIDEALFAKMQMITSIDLARNQLSTFNQTTFSQITTLSSIDLSQNPIEC
SCKSKWLIKLLRGA---IDVQNGKDTTCSFMKPFGGEALESIQPNDL--CTAFPVYFSAV
FFAVIF--VIFIIFVYHFRWQLRYKHFLLRLAI-GYREILDREDYDFDVYVISTDDDENW
IHDQLKPSFQRFLYSRNVFTEDDLPLGMHRTEAVDHVLTRSFKILVLVNKAACADDWFL
TCFRMAMDQVADTQTENIIVVFLENIEEDEMPLNVRLYMGGQGYVEWVEDDEGQKYF
WKRLEKCLSKH--
**>Sp_TLR100**
TFSSLGNLGTMYLQHNNLYNMWETIHVPFLKSLRRLKYLNLCYNGFQNIPNNSLSNLP
ELKALFLCHNKISHLQDNIINDL-PLTTLDLGHNQINLINQTLLEPLGTLKALTVSGNPFSC
GCDLQWFREWLDVT--QVHVDDNSHMICSSPPDMRGKLVIDFHPETLN-
CDHTWVLVGVG----SCMVFVTVALAVKFRFHINYCFNLVNARRRKYQRIKEDLPFLY
DAFVFFSHKDEEWVYNELVRHLEDDSGLRLCVHNRDFTLGRKILDNTIEAVDSSRFTLC
ILSADYLDSHWCKMEQEFAMANLIDR—DVLIIIALGEIPENKITKKLHKVMMKRTYLKW
PMEPVQRNDFWMKLKTVLREPNN
**>Hs_TLR1**
----TKSLLSLNMSSNIL--------DTIFRCLPRIKVLDLHSNKIKSIPK--VVKLEAL
QELNVAFNSLTDLPG—CGSFSSLSVLIIDHNSVSHPSADFFQSCQKMRSIKAGD
NPFQCTCELGEFVKNIDQV--SSEVEGWDSYKCDYPESYRGTLLKDFHMSELS-CN-
TLLIVTIVATMLVLVTVTSLCSYLDLPWYLRMVCQWTQTRRRARNPLEEQRNLQFHAFI
SYSGHDSFWVKNELLPNLEKE-GMQICLHERNFVPGKSIVENIITCIEKSYKSIFVLSP
NFVQSEWCHYELYFAHHNLFHEGSNSLILILLEPIPQYSIPSKLKSLMARRTYLEWPKEK
SKRGLFWANLRAAINKK--
**>Hs_TLR2**
----WPKMKYLNLSSTRIHSVT--------GCIPTLEILDVSNNNL-----NLSLNLPQL
KELYISRNKLMTLPDASL--LPMLLVLKISRNAITTFSKEQLDSFHTLKTLEAGGNNFIC
SCEFLSFTQEQQALA-KVLIDWPANYLCDSPSHVRGQQVQDVRLSVSE-
CHRTALVSGMCCA-LFLLILLTGVLCHRFHWYMKMMWAWLQAKRKPRK--
APSRNICYDAFVSYSERDAYWVENLMVQELENNPPFKLCLHKRDFIPGKWIIDNIIDSIE
KSHKTVFVLSENFVKSEWCKYELDFSHFRLFDENNDAILILLEPIEKKAIPQKLRKIMNT
KTYLEWPMDEAQREGFWVNLRAAIKS---
**>Hs_TLR3**
MLEGLEKLEILDLQHNNLARLWHANPGPIYKGLSHLHILNLESNGFDEIPVEVFKDLFEL
KIIDLGLNNLNTLPASVFNNQVSLKSLNLQKNLITSVEKKVFPAFRNLTELDMRFNPFDC
TCESIWFVNWINET--HTNIPELSHYLCNTPPHYHGFPVRLFDTS---SCKDPFELFFMI
TS-ILLIFIFIVLLIHFEGWRISFYWNVSVHRV-GFKEIDQTEQFEYAAYIIHAYKDKDW
VWEHFSSMEKEDQSLKFCLEERDFEAGVFELEAIVNSIKRSRKIIFVITHHLLKDPLCRF
KVHHAVQQAIEQNLDSIILVFLEEIPDYKLNHARRGMFKSHCILNWPVQKERIGAFRHKL
QVALGSKNS
**>Hs_TLR4**
IFNGLSSLEVLKMAGNSF----ENFLPDIFTELRNLTFLDLSQCQLEQLSPTAFNSLSSL
QVLNMSHNNFFSLDTFPYKCLNSLQVLDYSLNHIMTSKKQELQHFSSLAFLNLTQNDF
ACTCEHQSFLQWIKDQ--RQLLVEVERMECATPSDKQGMPVLSLNIT----
CQMKTIIGVSVLS--VLVVSVVAVLVYKFYFHLMLLAG-------KYG----
RGENIYDAFVIYSSQDEDWVRNELVKNLEEGPPFQLCLHYRDFIPGVAIAANIHEGFHK

SRKVIVVVSQHFIQSRWCIFEYEIAQTWQFLSSRAGIIFIVLQKVEKTLLRQELYRLLSRN
TYLEWEDSVLGRHIFWRRLRKALLDEQE
**>Hs_TLR5**
-----PSLEQLFLGENMLQLAWTELCWDVFEGLSHLQVLYLNHNYLNSLPPGVFSHLTAL
RGLSLNSNRLTVLSH---NDLPNLEILDISRNQLLAPNPDVF---VSLSVLDITHNKFIC
ECELSTFINWLNHT--NVTIAGPADIYCVYPDSFSGVSL--FSLSTEG-CDEKFFIVCTV
TLT---LFLMTILTVTKFRGFCFICYKAQRLVFKDHPQGTEPDMYKYDAYLCFSSKDFTW
VQNALLKHLDDQNRFNLCFEERDFVPGENRIANIQDAIWNSRKIVCLVSRHFLRDGWC
LEAFSYAQGRCLSDLNSALIMVVVGSLSQYQLHQSIRGFVQKQQYLRWPEDLQDVGW
FLHKLSQQILKKQT
**>Hs_TLR6**
----VESIVVLNLSSNML--------DSVFRCLPRIKVLDLHSNKIKSVPK--VVKLEAL
QELNVAFNSLTDLPG--CGSFSSLSVLIIDHNSVSHPSADFFQSCQKMRSIKAGD
NPFQCTCELREFVKNIDQV--SSEVEGWDSYKCDYPESYRGSPLKDFHMSELS-CN-
TLLIVTIGATMLVLVTVTSLCIYLDLPWYLRMVCQWTQTRRRARNPLEEQRNLQFHAFIS
YSEHDSAWVKSELVPYLEKE-DIQICLHERNFVPGKSIVENIINCIEKSYKSIFV
LSPNFVQSEWCHYELYFAHHNLFHEGSNNLILILLEPIPQNSIPNKLKALMTQRTYLQW
PKEKSKRGLFWANIRAAFNKEKS
**>Hs_TLR7**
VFDGMPNLKNLSLAKNGLKSF-----WKKLQCLKNLETLDLSHNQLTTVPERLSNCSRSL
KNLILKNNQIRSLTKYFLQDAFQLRYLDLSSNKIQMIQKTSFNVLNNLKMLLLHHNRFLC
TCDAVWFVWWVNHT--EVTIPYLTDVTCVGPGAHKGQSVISLDLYT---CELLILFSLSI
SVSLFL--MVMMTASHLYFWDVWYIYHFCKAKIKGYQRLISPDCC-YDAFIVYDTKDPEW
VLAELVAKLEREKHFNLCLEERDWLPGQPVLENLSQSIQLSKKTVFVMTDKYAKTENF
KIAFYLSHQRLMDEKVDVIILIFLEKPFQKSKFLQLRKRLCGSSVLEWPTNPQAHPYFW
QCLKNALATDNH
**>Hs_TLR8**
AFLNLPSLTELHINDNMLKFF-----WTLLQQFPRLELLDLRGNKLLFLTDSLSDFTSSL
RTLLLSHNRISHLPSGFLSEVSSLKHLDLSSNLLKTINKSALKTTTKLSMLELHGNPFEC
TCDIGDFRRWMDHL--NVKIPRLVDVICASPGDQRGKSIVSLELTT---CVSVILFFFTF
FITTMV--MLAALAHHLFYWDVWFIYNVCLAKVKGYRSLSTSQTF-YDAYISYDTKDADW
VINELRYHLERDKNVLLCLEERDWDPGLAIIDNLMQSINQSKKTVFVLTKKYAKSWNFK
TAFYLALQRLMDENMDVIIFILLEPVLQHSQYLRLRQRICKSSILQWPDNPKAEGLFWQT
LRNVVLTEND
**>Hs_TLR9**
TLRNLPSLQVLRLRDNYLAFF-----WWSLHFLPKLEVLDLAGNQLKALTNGSLPAGTRL
RRLDVSCNSISFVAPGFFSKAKELRELNLSANALKTVDHSWFPLASALQILDVSANPLH
CACGAA-FMDFLLEV--QAAVPGLSRVKCGSPGQLQGLSIFAQDLRL---CLDWDC
FALSLLAVALG—LGVPMLHHLCGWDLWYCFHLCLAWLRGRQSGRDEDALPYDA
FVVFDKTQSDWVYNELRGQLERGRALRLCLEERDWLPGKTLFENLWASVYGSRKTLF
VLAHTDRVSGLLRASFLLAQQRLLEDRKDVVVLVILSPDGRRSRYVRLRQRLCRQSVL
LWPHQPSGQRSFWAQLGMALTRDNH
**>Hs_TLR10**
-----PTVVNMNLSYNKL--------DSVFRCLPSIQILDLNNNQIQTVPK--TIHLMAL
RELNIAFNFLTDLPG--CSHFSRLSVLNIEMNFILSPSLDFVQSCQEVKTLNAGRNPFRC
TCELKNFIQLETYS--EVMMVGWDSYTCEYPLNLRGTRLKDVHLHELS-CNTALLIVTIV

VIMLVLLAVAFCCLHFDLPWYLRMLGQCTQTWHRVRKTQEQKRNVRFHAFISYSEHD
SLWVKNELIPNLEKEGSILICLYESYFDPGKSISENIVSFIEKSYKSIFVLSPNFVQNEWC
HYEFYFAHHNLFHENSDHIILILLEPIPFYCIPTKLKALLEKKAYLEWPKDRRKCGLFWAN
LRAAINNEQT
**>Nv_TLR**
-LHRLPKMP-VNLRGNAIRELP--------------------------------HYLGNI
TVLELSNNEIKELNMTFVDSLARVVNLAINDNKLKYLPRGVTNLTEGFRSLSISHNFFVC
DCYASWMRDWLANNTD--KIEDTSSILCASG--LEGLPIISVP--LSDNCSALSLLLAIV
LAVLLVLSVVAFVMTYCFRWEMKILMYHFNWHPR--D--DTDVSKIYDTFISYSSQDASW
VRETLQRTLESVPPYRLCIHDRDFEIGASIHDNILNSVRLSKRMIMVLSNHFIASEWCRL
EFRAAHQKVLEDRTNYLIIILFDDVDPSTLDDETKLYLRTNTYLSVSN------WFWQKL
FYALPKPLA
**>Ad_TLR1**
-LVAMPSVP-MFLQSNNIREIP--------------------------------GYLENV
TSLYLSHNQIQRLDEKTIDRLKRIETLFIDSNKLTTLPRNIENVS--FTKISLQHNFFRC
DCETKWMKQWLLREEA--HVDNIENILCHSD--VQGKAISRLP--DEELCLEAFKITAYT
LGGLLFVSLVAFAVGYKFRSEAKVFMYHFNWHPR--N--
DLDPNKPYDAFISFSGNDYEW
ICNTLCVRLENDPPYKLCLHHRDFLVGAPIQQNIFDGIERSKRMIMTLSKHFVRSEWCLL
EFRAAHQKVLEDRINYLIIILFDDVDMAEVDDEIKLYMRTNTYLSVKN------WFWEKL
FYALPQNTN
**>Ad_TLR2**
----MPSVP-LFLQSNKIEEIP--------------------------------SYLENV
TALYLSHNNIERLNEKTIDRLKRIEILFIDSNKLTTLPRNIENVS--FIKISLQHNFFRC
DCKTKWMKHWLLRQEA--HIDNIENILCHSD--VKGKAISRLP--DEEVCPAAFKITAYT
LGGLLLVFLVAFAVGYKFRGEVKVFMYHFNWHPR--N--DLDPNKIYDAFISFSGIDYEW
ISNTLCARLENDPPYKLCLHHRDFLVGAPIQQNIFNGIEKSKRMIMILSKNFVKSEWCLL
EFRAAHQKVLEDRINYLIIILFDDVDMAEVDDEIKLYMRTNTYLSIKN------WFWEKL
FYALPQNSK
**>Ad_TLR3**
-LVAMPSVP-LLLQSNNIREIP--------------------------------GYLENV
TQLYLSHNNIERLDEKTIDRLKRIEKLFIDSNKLTTLPRNIENVS--FTKIALQHNLFRC
DCKTKWIKHWLSRQED--HIEHIDNILCHSD--VKDKVISNLP--DEEVCLEAFKITACT
LGGLLFMFLLAFAVGYKFRSEGKVFMYHFNWHPR--N--
DSNPNKTYDAFISFSGNDYAW
ISNTLCARLENDPPYNLCLHHRDFLVGAPIQQNIFNAIEKSKRMIMILSKNFVKSEWCLL
EFRAAHQKVLEDRINYLIIILFDDVDMAEVDDEIKLYMRTNTYLSVKN------WFWEKL
FYALPQNSN
**>Ad_TLR4**
-LVTMPSVP-LFLQSNDIQEIP--------------------------------RYLENV
TSLYLSHNQIERLDEKTIDRLKRIQVLFIDSNKLTTLPRNIENVS--FTKISMQHNFFRC
DCKTKWMKQWLLREEA--HVDNIENILCHSN--VQGKAISRLP--DEKVCLEAFKITAYT
LGCLFLVFLVAFAVGYKFRSEAKVFMYHFNWHPR--K--
DSDPNKIYDAFVSFSGNDYEW
ISNTLCVRLENDPPYKLCLHHRDFLVGAPIQQNIFNGIEKSKRMIMILSKNFVRSEWCLL
EFRAAHQKVLEDRINYLIIILFDDVDMAEVDDEIKLYMRTNTYLSVKN------WFWEKL

FYALPQNSN
**>Am_TLR**
-FVAMPSLP-LFLQSNNIREIP-------------------------------GYLENV
TSLYLSHNQIERLDEKTIDRLKRIQVLFIDSNKLTTLPRNIENVS--FTKISLQHNFFRC
DCKTKWMKQWLLREEA--HVDNIENILCHSD--VQGKAISRLP--DEEVCLDAFKITAYT
LGGLLLVFLVAFAVGYKFRGEVKVFMYHFNWHPR--N--DLDPNKPYDAFISFSGIDYEW
ISNTLCVRLENDPPYKLCLHHRDFLVGAPIQQNIFDGIERSKRMIMILSKHFVKSEWCLL
EFRAAHQKVLEDRINYLIIILFDDVDMAEVDDEIKLYMRTNTYLSVKN------WFWEKL
FYALPQNSN
**>Of_TLR**
-LTAIPKVP-LKLEDNNIREIP-------------------------------PYMENV
TALYLTHNKIQVLNKSTVRRFTRIKVLFIDSNKLTYLPKNIENLN--FTSLALHHNFFKC
DCTTLWIKHWLQRKQS--KILHIKNVLCNSE--TQGKAIYTLP--NEEVCKKTFKIIALT
LGGALVLTFIAFIVAYKYRGEMKVLMYHFNWHPR--D--DSDPRKIYDAFVSYSGSDHQW
VVNTLQERLEHDPPYKLCIHHRDFVVGAPIQENILNSVDQSKRMLMVLSRNFLKSEWC
LLEFRAAHRKVLEDRMNYLIIILFDGINMDELDDEMKLYMRTNTYLSVSY------WFWEKL
YYAMPQSTD
>Mm_MyD88
----NPTV----------------------ADWTLLAEE--------------FEY-LEI
RELETRPDPTSLLDAASVGRL-ELALLDRE-DILKE-----------LK--RIEEDC---
-------QKYLKQQNQ-----------------SE--KLQV-RVE-SVQ-----------
LG-G----ITT--------------------------DD-LGTPELFDAFICYCPNDIEF
V-
QEMIRQLEQDYRLKLCVSDRDVLPGTCVWSIASELIKRCRRMVVVVSDDYLQSKECDF
QTKFALSLSPGVQQKRLIPIKYKAMKK-----DFPSILRFITICDYTN--CTKSWFWTRL
AKALSLP--
>Dm_MyD88
------KLRS----------------EEGYRDWRGISEL-------QKVDE---------
-----ANNPMLVLISQTVGHLEHLGIIDIQENLAKDTQ----RFI--MKAL------EAC
-C----FNNY--SSSNNITV---QSVQ-----------I--L---DEDRCV---------
MG--------------------------------------Q-PLPRYNACVLYAEADIDH
A-
TEIMNNLESRYNLRLFLRHRDMLMGVPFEHVQSHFMTRCNHLIVVLTEEFLRSPENTY
LVNFTQKIQIENHTRKIIPILYKDM-------HIPQTLGIYTHIKYAGDS----NFWDKL
ARSLHRQPS
>Bg_MyD88
----DIDVMH----------------N-GYPNYGGLAEL--------------FSG-LEI
MEFERAKSPTALLLEPKIGKL-ELFQLDRI-DVLTD-----------CH--AINDEA---
-------KLFIKESKA-----------------LL--KIQD-TVTPGSTPPQ--------
L---------------------------------------TGKKTYYDAYICYNPKDLEF
V-RELISRFESKYRFTLFVPFRDDLPGMNEHAINAKIIERCRHMIVVLSKNFLQSEACEF
QSTFAQSLSPGARNKRIVPIKIEDC-------VIPNILRIMACCDFTK--DLWDWSWDRL
ARSITAQQS
>La_MyD88
----NPSVLT----------------DSGFKDWKGLAEL--------------FMN-DDI
SNFESMRDSTLILKDSMIATL-DIQQLERY-DILEDPT---------IK--QVENNV---

-------KSYLRQSKE-----------M-----IE--KLQD-EVSMEYD--E--------
YK-Y----MTT-------------------------DV-TGKAMYYDAFVSYAEEDIGF
V-KKLISELEKEYGLKLCVQARDLIPGASTNTVCAKLIERCTRMVIILSPKFLNSAQCDF
QIKFAQSLSPGSRGKKLVPIMYKCC-------DIPSILRHIAICDYTK--DLQEWFWNRL
AMSLKAP--
>Aq_MyD88
-----QNL---------------------SDWRTLVEL-------------GFTY-EMV
MLLRAKNSPTSLLEEATLESL--IVMMELENV----------SALQQVK--SIH-D----
RPPP--RK---PNSCS----SDPSTMVA--P-AVPSLKYVDVNLQTNSDCEQ--------
LSS-----LSL---------------------------DHGDGSDFDIFLSFAPADAEF
A-DEMRLRLIN-AGISVYIASEGLMPGQSFIDEVADKIRGCRKTIIILSPDYNQCSWCNY
EARLAHHKNPDPKRHTLIPIVYRKC-------EVPDFMSHLFYLDFSRDHQCEKYFWDRL
YKSVRHQ--
>Lp_MyD88
----NPQLTT---------------LTGLRDFRGVAEL--------------FDY-WDI
QNFQMTSDPFKLLIQNTVGKL-DLEEIERT-DVIDD-----------VK--YAEIDA---
-------HNYLRKQKW-----------C-----KE--SLQD-NVS---SCTA--------
--------LTV-------------------------DVNEGEVSLYDAYVCYTDEDIEC
V-HILSRQLESE-GIRLFIRDRDLLLGQMEYEAFARLIERCNRVLIVLSPEFLKSVECEF
QTRYATSLAVEQQQRKLIPIIYRSC-------DVPHLLRYLSKIDFTK--HIHDWVWHRL
IHSIKGERE
>Spi_MyD88
----RPCL----------------------NDYRLLAAK--------------YTN-EEI
KYLESLREPVELMTRRTIAEL-SLQQIDRP-DVVQD-----------LQ--YI--DS---
-------TPFEKEREQ-----------------------------------------
--------RTN-------------------------NVQRTVRKSYHAFVCFAEEDKAF
V-DNLVKKMENNRNLHLCLPVRDFLPVGSHLETTALAIQRCKKFIVILSKNYDSSQGAIY
QAQIATSLAPGAKEKRIIPVLIDEC-------SIPRTLSHITYLDYLR--DEKH-FWNLL
CDTLTRN--
>Spu_MyD88
----RPRG----------------------CDWRDMAEE--------------FSYQLHI
QNFALESDPVKVLSASNVGKL-DIEKIERH-DVLHE-----------LP--FLEEDC---
-------KRW-KRTQA-----------------AR--DIQV-EVT-NFS--S--------
LR-G----ITL--------------------------DS-SGPPEMFDAYVCFAMADLEF
V-QQLRSQLESPHNYKLCIDQRDLLPGGSHALVTAEIINRCNKMLVILSPEFLQSPSCDF
QTKFAVSLEPGAMKRRIIPILVKPC-------DLPLIIRHITLCDFTK--DLRPWFWGRL
RKAMSIR--
>Cg_MyD88
----DPGVTG----------------D--YNDYQGLAEV--------------FTF-QDI
TNFQRQSKPTEMLYQPTVDNL-KLQPIGRSDDVITE-----------CA--LIKKDV---
-------DRYCKDITG-----------------SK--DIQD-SVSPNRSCDS--------
LG-L----VTI-------------------------DVEKGDTLYYDAFVIYNPKDLEF
V-
KELAGKMEAPYNLKFCIPWRDDLPGGSRYEVSAHMITRCRRTLVILSSDFLKSAAADF
QLKFAHCLSPGARSKKVVPVFSAPC-------KMPGILRAVSFVDFTN--GLRDWNWPRL
NAVLRCPRD
