## Supplementary Figure 2 for "The evolution of the metazoan Toll receptor family and its expression during protostome development"

**Supplementary Figure 2 - Second phylogenetic analysis, excluding the 150-200 aminoacid region.** Parameters applied for the construction of this phylogenetic tree are the same than the ones applied for the main phylogenetic analysis (Figure 4A). Bootstrap values are indicated next to the main nodes and all nodes with bootstrap values >60 are marked with full black dots.

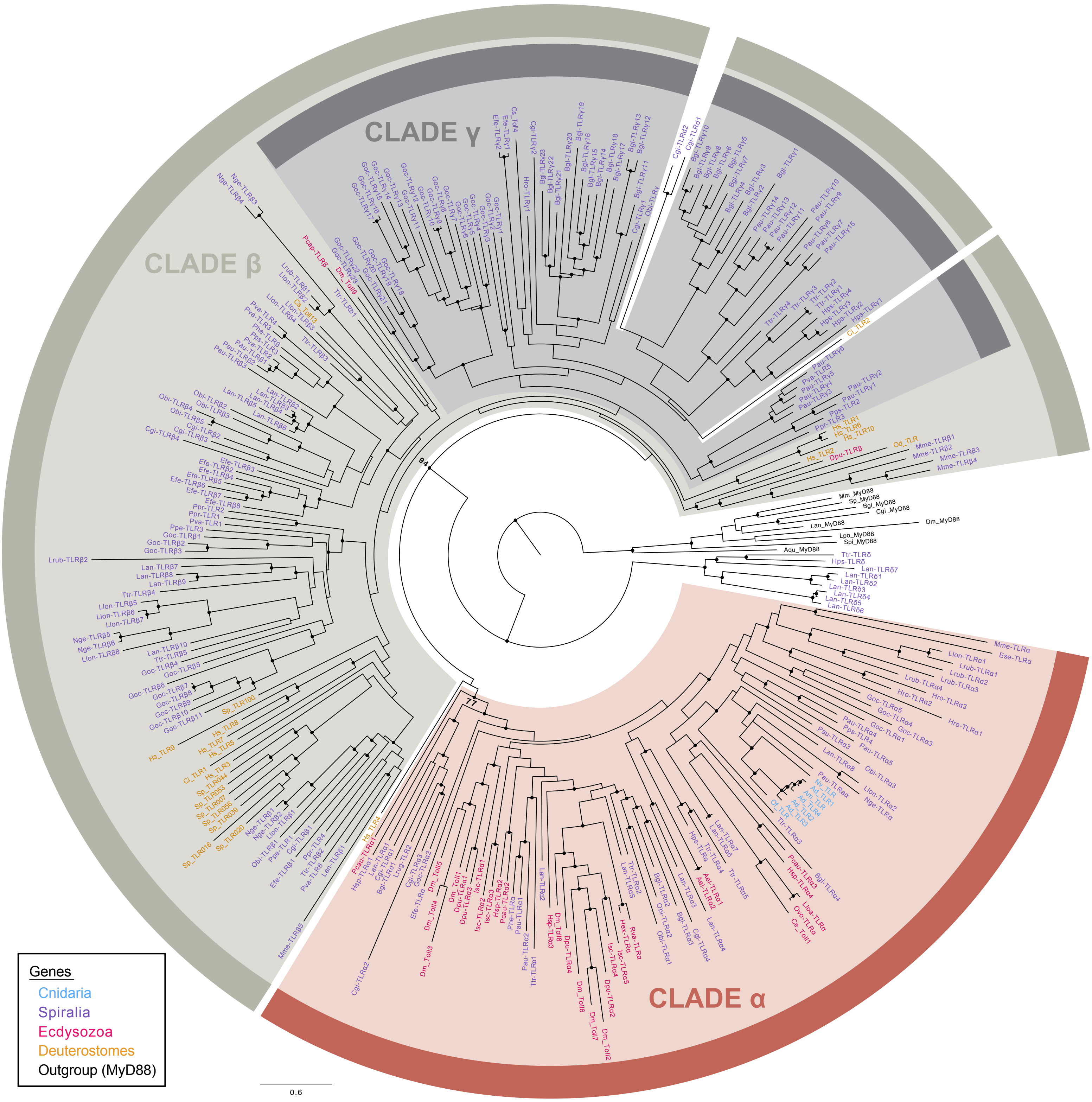
