## Supplementary Figure 3 for "The evolution of the metazoan Toll receptor family and its expression during protostome development"

**Supplementary Figure 3 - Third phylogenetic analysis, excluding the 349-354 aminoacid region.** Parameters applied for the construction of this phylogenetic tree are the same than the ones applied for the main phylogenetic analysis (Figure 4A). Recovered clades are named α, β and γ. Comparison with the main phylogenetic analysis is represented with blue and magenta dots. Bootstrap values are indicated next to the main nodes and all nodes with bootstrap values >60 are marked with full black dots.

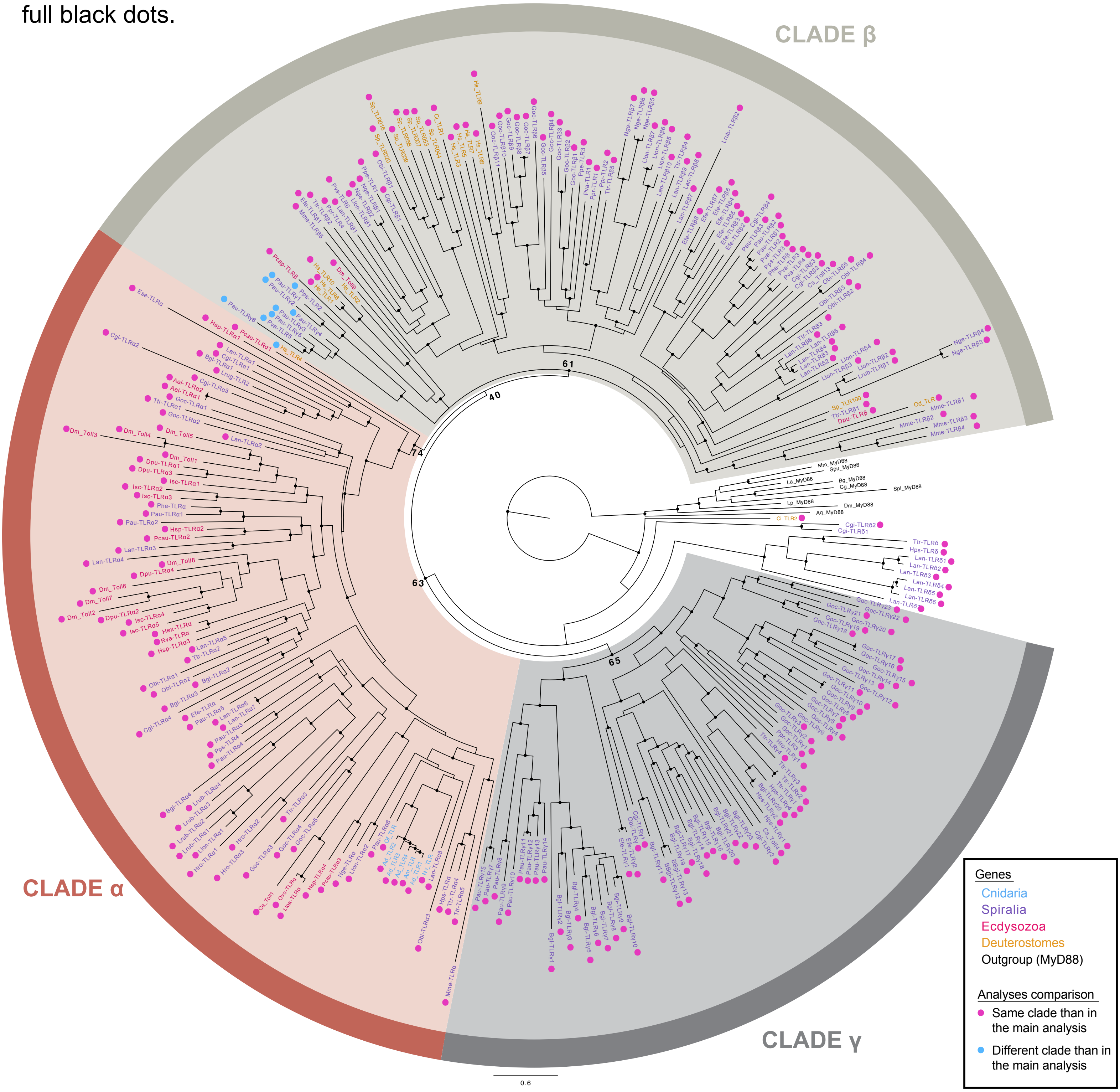
