## Supplementary Table 1 for "The evolution of the metazoan Toll receptor family and its expression during protostome development"

**Supplementary Table 1. Species included in our study.** See supplementary Table 2 for the accession number of individual sequences.

| | Species | TLR Source | Publication | Genome/transcriptome NCBI Accession number | Transcriptome BUSCO values |
|---|---|---|---|---|---|
| **Cnidaria** | *N. vectensis* | Literature | [1] | - | - |
| | *A. digitifera* | Literature | [2] | - | - |
| | *A. millepora* | Literature | [2] | - | - |
| | *O. faveolata* | Literature | [3] | - | - |
| **Xenacoelomorpha** | *X. profunda* | This study: Genome | Unpublished | | - |
| | *H. miamia* | This study: Genome | - | GCA004352715 | - |
| | *P. naikaiensis* | This study: Genome | - | PRJDB7329 | - |
| | *I. pulchra* | This study: Genome | Unpublished | | - |
| | *M. stichopi* | This study: Genome | Unpublished | | - |
| | *C. macropyga* | This study: Transcriptome | [4] | SRX1343815 | 89.2% |
| **Bryozoa** | *M. membranacea* | This study: Transcriptome | - | SRX1121923 | 96.9% |
| | *B. neritina* | This study: Transcriptome | [5] | | 96.6% |
| **Cycliophora** | *S. pandora* | This study: Transcriptome | [6] | SRX1531719 | 87.4 |
| **Annelida** | *G. oculata* | This study: Transcriptome. | Unpublished | | 99% |
| | *E. fetida* | This study: Transcriptome | - | SRX3108745 | 96.2% |
| | *H. robusta* | This study: Genome | [7] | AMQM00000000.1 | - |
| | *P. prolifica* | Literature | [8] | - | - |
| **Mollusca** | *C. gigas* | This study: Genome | [9] | AFTI00000000 | - |
| | *O. bimaculoides* | This study: Genome | [10] | PRJNA270931 | - |
| | *C. sinensis* | Literature | [11] | - | - |
| | *L. rugatus* | Literature | [8] | - | - |
| | *B. glabrata* | This study: Genome | [12] | APKA00000000.1 | - |

| | | | | | |
|---|---|---|---|---|---|
| | *B. glabrata* | Individual sequence(s) downloaded from NCBI | - | - | - |
| **Brachiopoda** | *T. transversa* | This study: Transcriptome. | [4] | SRX1307070 | 95.7% |
| | *H. psittacea* | This study: Transcriptome. | [8] | SRX731469 | 94.5% |
| | *L. anatina* | This study: Genome | [13] | LFEI00000000 | - |
| **Micrognathozoa** | *L. maerski* | This study: Transcriptome. | | SRX1121929 | 93.8% |
| **Gastrotricha** | *L. squamata* | This study: Transcriptome. | [14] | SRX1000997 | 89.6%. |
| | *Macrodasys sp* | This study: Transcriptome. | [15] | SRX534826 | 75.9% |
| | *Megadasys sp* | This study: Transcriptome. | [15] | SRX534835 | 70% |
| | *D. aspetos* | This study: Transcriptome. | | SRX1121926 | 90% |
| | *M. laticaudatus* | This study: Transcriptome. | | SRX872416 | 82.5% |
| **Nemertea** | *Lineus longissimus* | This study: Transcriptome. | [4] | SRX1343823 | 95.2% |
| | *Lineus ruber* | This study: Transcriptome. | Unpublished | | 95% |
| | *N. geniculatus* | This study: Genome | [16] | NMRB00000000 | - |
| | *P. peregrina* | Literature | [8] | - | - |
| **Phoronida** | *P. harmeri* | This study: Transcriptome. | | SRX1121914 | 90.4% |
| | *P. australis* | This study: Genome | [16] | NMRA00000000 | - |
| | *P. psammophila* | Literature | [8] | - | - |
| | *P. vancouverensis* | Literature | [8] | - | - |
| **Platyhelminthes** | *M. lignano* | This study: Genome | [17] | SRP059553 | - |
| | *E. multilocularis* | This study: Genome | [18] | PRJEB122 | - |
| | *H. microstoma* | This study: Genome | [18] | PRJEB124 | - |
| | *S. mansoni* | Literature | [19] | - | - |
| | *S. mediterranea* | Literature | [20] | - | - |
| **Rotifera** | *E. senta* | This study: Transcriptome. | Unpublished | | 95.2% |

| | | | | | |
|---|---|---|---|---|---|
| | *R. tardigrada* | This study: Transcriptome. | [21] | SRX1253177 | 91.1% |
| | *E. gadi* | This study: Transcriptome. | | SRX1121912 | 74.9% |
| | *M. hirudinaceus* | This study: Transcriptome. | [15] | PRJEB5803 | 84.5%. |
| | *A. vaga* | Literature | [22] | - | - |
| **Priapulida** | *P. caudatus* | This study: Transcriptome. | [4] | SRX507009 | 93.9% |
| | *H. spinulosus* | This study: Transcriptome | [4] | SRX1343820 | 96.4% |
| **Tardigrada** | *H. exemplaris* | This study: Genome | [23] | SRX2495681 | - |
| | *R. varieornatus* | This study: Genome | [24] | DRX012456 | - |
| **Onychophora** | *P. capensis* | This study: Transcriptome. | [25] | SRX451023 | 62% |
| **Nematoda** | *L. loa* | This study: Genome | [26] | ADBU00000000.2 | - |
| | *O. volvulus* | This study. Genome | [27] | CBVM000000000 | - |
| | *C. elegans* | Individual sequence(s) downloaded from NCBI | - | - | - |
| **Loricifera** | *A. elegans* | This study: Transcriptome. | | SRX1120677 | 36.2% |
| **Arthropoda** | *D. pulex* | This study: Genome | [28] | ACJG00000000 | - |
| | *D. melanogaster* | Individual sequence(s) downloaded from NCBI | - | - | - |
| | *I. scapularis* | Literature | [29] | - | - |
| **Tunicata** | *C. intestinalis* | Literature | [30] | - | - |
| | *O. dioika* | Literature | [31] | - | - |
| **Echinodermata** | *S. purpuratus* | Literature | [32] | - | - |
| **Craniata** | *H. sapiens* | Individual sequence(s) downloaded from NCBI | - | - | - |
