## Supplementary Table 3 for "The evolution of the metazoan Toll receptor family and its expression during protostome development"

**Supplementary Table 3** - *Hypsibius exemplaris* stage specific transcriptome analyses (RSEM and Kallisto methods)

| *HYPSIBIUS EXEMPLARIS* | | | | | | | | | | | | | | | | | | | |
|---|---|---|---|---|---|---|---|---|---|---|---|---|---|---|---|---|---|---|---|
| Values inicate Transcripts per Milion (TEM) | | | | | | | | | | | | | | | | | | | |
| RSEM | Zigot | Morula1 | Morula2 | Morula3 | Early gastrula | Gastrula 1 | Gastrula 2 | Elongation | Segmentation 1 | Segmentation 2 | Limb bud formation | Differentiation 1 | Differentiation 2 | Differentiation 3 | Differentiation 4 | Differentiation 5 | Differentiation 6 | Differentiation 7 | Differentiation 8 |
| Hexe-TLRα | 3,165 | 0,000 | 0,000 | 8,589 | 8,071 | 8,335 | 1,421 | 9,685 | 25,103 | 0,000 | 0,000 | 0,000 | 27,095 | 0,000 | 0,000 | 0,000 | 0,000 | 0,000 | 0,000 |
| kallisto | Zigot | Morula1 | Morula2 | Morula3 | Early gastrula | Gastrula 1 | Gastrula 2 | Elongation | Segmentation 1 | Segmentation 2 | Limb bud formation | Differentiation 1 | Differentiation 2 | Differentiation 3 | Differentiation 4 | Differentiation 5 | Differentiation 6 | Differentiation 7 | Differentiation 8 |
| Hexe-TLRα | 0,000 | 0,000 | 0,000 | 27,214 | 0,000 | 9,697 | 0,000 | 0,000 | 33,087 | 0,000 | 0,000 | 0,000 | 77,060 | 0,000 | 0,000 | 0,000 | 0,000 | 0,000 | 0,000 |

TEM ≥0,150

TEM <0,150

**Supplementary Table 4** - Priapulus caudatus stage specific transcriptome analyses. Analyses for the different methods (RSEM and Kallisto) and replicates (Rep 1_and Rep_2). For each method, average and standard error (SE) of the two replicates is provided.

***PRIAPULUS CAUDATUS***

**Values inicate Transcripts per Milion (TEM)**

| **Rep_1_RSEM** | **0d** | **1d** | **3d** | **5d** | **9d** |
|---|---|---|---|---|---|
| Pcau-TLRα1 | 2,551 | 2,294 | 1,509 | 1,016 | 1,512 |
| Pcau-TLRα2 | 1,471 | 2,427 | 1,822 | 1,983 | 2,988 |
| Pcau-TLRα3 | 0,000 | 0,000 | 0,822 | 2,371 | 5,496 |

| **Rep_1_kallisto** | **0d** | **1d** | **3d** | **5d** | **9d** |
|---|---|---|---|---|---|
| Pcau-TLRα1 | 2,433 | 2,059 | 1,487 | 0,940 | 1,589 |
| Pcau-TLRα2 | 1,440 | 2,093 | 1,767 | 1,890 | 3,089 |
| Pcau-TLRα3 | 0,000 | 0,000 | 0,825 | 2,156 | 4,627 |

| **Rep_2_RSEM** | **0d** | **1d** | **3d** | **5d** | **9d** |
|---|---|---|---|---|---|
| Pcau-TLRα1 | 7,824 | 6,555 | 3,418 | 2,230 | 3,617 |
| Pcau-TLRα2 | 0,680 | 2,285 | 1,477 | 2,630 | 5,675 |
| Pcau-TLRα3 | 0,000 | 0,000 | 0,167 | 1,159 | 3,396 |

| **Rep_2_kallisto** | **0d** | **1d** | **3d** | **5d** | **9d** |
|---|---|---|---|---|---|
| Pcau-TLRα1 | 7,609 | 6,107 | 3,311 | 1,985 | 3,445 |
| Pcau-TLRα2 | 0,724 | 2,432 | 1,475 | 2,469 | 6,272 |
| Pcau-TLRα3 | 0,000 | 0,000 | 0,189 | 0,667 | 3,275 |

| **RSEM_average** | **0d** | **1d** | **3d** | **5d** | **9d** |
|---|---|---|---|---|---|
| Pcau-TLRα1 | 5,188 | 4,425 | 2,464 | 1,623 | 2,565 |
| Pcau-TLRα2 | 1,076 | 2,356 | 1,650 | 2,307 | 4,332 |
| Pcau-TLRα3 | 0,000 | 0,000 | 0,495 | 1,765 | 4,446 |

| **Kallisto_average** | **0d** | **1d** | **3d** | **5d** | **9d** |
|---|---|---|---|---|---|
| Pcau-TLRα1 | 5,021 | 4,083 | 2,399 | 1,463 | 2,517 |
| Pcau-TLRα2 | 1,082 | 2,263 | 1,621 | 2,180 | 4,681 |
| Pcau-TLRα3 | 0,000 | 0,000 | 0,507 | 1,412 | 3,951 |

**Values indicate Standard Error (SE)**

| **RSEM_SE** | **0d** | **1d** | **3d** | **5d** | **9d** |
|---|---|---|---|---|---|
| Pcau-TLRα1 | 2,637 | 2,131 | 0,955 | 0,607 | 1,053 |
| Pcau-TLRα2 | 0,396 | 0,071 | 0,172 | 0,324 | 1,344 |
| Pcau-TLRα3 | 0,000 | 0,000 | 0,328 | 0,606 | 1,050 |

| **Kallisto_SE** | **0d** | **1d** | **3d** | **5d** | **9d** |
|---|---|---|---|---|---|
| Pcau-TLRα1 | 2,588 | 2,024 | 0,912 | 0,523 | 0,928 |
| Pcau-TLRα2 | 0,358 | 0,170 | 0,146 | 0,289 | 1,592 |
| Pcau-TLRα3 | 0,000 | 0,000 | 0,318 | 0,745 | 0,676 |

TEM ≥0,150

TEM <0,150

**Supplementary Table 5** - *Crassostrea gigas* stage specific transcriptome analyses (RSEM and Kallisto methods)

*CRASSOSTREA GIGAS*

**Values inicate Transcripts per Milion (TEM)**

| RSEM | Early morula | Morula | Blastula | Rotary movement | Free swimming | Early gastrula | Gastrula | Trochophore 1 | Trochophore 2 | Trochophore 3 | Trochophore 4 | Trochophore 5 | early Dshaped larva 1 | early Dshaped larva 2 | Dshaped larva 1 | Dshaped larva 2 | Dshaped larva 3 | Dshaped larva 4 | Dshaped larva 5 |
|---|---|---|---|---|---|---|---|---|---|---|---|---|---|---|---|---|---|---|---|
| Cgi-TLRα1 | 0,000 | 0,000 | 0,163 | 0,507 | 0,446 | 0,136 | 0,279 | 0,132 | 0,758 | 0,331 | 0,249 | 0,646 | 0,586 | 0,666 | 0,371 | 0,315 | 0,265 | 0,279 | 0,287 |
| Cgi-TLRα2 | 0,048 | 0,121 | 0,154 | 0,119 | 0,085 | 0,027 | 0,058 | 0,042 | 0,000 | 0,052 | 0,065 | 0,055 | 0,053 | 0,047 | 0,000 | 0,000 | 0,000 | 0,052 | 0,000 |
| Cgi-TLRα3 | 0,000 | 0,028 | 0,029 | 0,000 | 0,000 | 0,000 | 0,058 | 0,000 | 0,124 | 0,114 | 0,195 | 0,109 | 0,120 | 0,257 | 0,379 | 0,420 | 0,054 | 0,279 | 0,231 |
| Cgi-TLRα4 | 0,000 | 0,000 | 0,000 | 0,075 | 0,276 | 0,345 | 0,221 | 0,118 | 0,310 | 0,362 | 0,498 | 0,777 | 0,918 | 0,947 | 1,190 | 0,954 | 1,183 | 1,311 | 2,648 |
| Cgi-TLRβ1 | 0,000 | 0,000 | 0,000 | 0,000 | 0,032 | 0,045 | 0,000 | 0,000 | 0,108 | 0,052 | 0,000 | 0,044 | 0,000 | 0,047 | 0,098 | 0,097 | 0,048 | 0,052 | 0,311 |
| Cgi-TLRβ2 | 0,019 | 0,028 | 0,000 | 0,000 | 0,021 | 0,000 | 0,000 | 0,000 | 0,000 | 0,052 | 0,000 | 0,000 | 0,000 | 0,000 | 0,091 | 0,000 | 0,048 | 0,000 | 0,191 |
| Cgi-TLRβ3 | 0,000 | 0,121 | 0,029 | 0,104 | 0,000 | 0,000 | 0,000 | 0,000 | 0,000 | 0,000 | 0,065 | 0,000 | 0,000 | 0,000 | 0,053 | 0,000 | 0,054 | 0,000 | 0,072 |
| Cgi-TLRβ4 | 0,381 | 1,021 | 0,807 | 0,343 | 0,266 | 0,209 | 0,581 | 0,439 | 0,511 | 0,671 | 0,217 | 0,361 | 0,373 | 0,175 | 0,280 | 0,291 | 0,088 | 0,093 | 0,255 |
| Cgi-TLRγ1 | 0,000 | 0,000 | 0,000 | 0,000 | 0,000 | 0,000 | 0,000 | 0,000 | 0,000 | 0,300 | 0,054 | 0,000 | 0,093 | 0,117 | 0,083 | 0,040 | 0,041 | 0,083 | 0,000 |
| Cgi-TLRγ2 | 0,000 | 0,000 | 0,000 | 0,000 | 0,000 | 0,000 | 0,000 | 0,063 | 0,093 | 0,000 | 0,000 | 0,077 | 0,000 | 0,082 | 0,159 | 0,154 | 0,000 | 0,083 | 0,000 |
| Cgi-TLRδ1 | 0,029 | 0,000 | 0,000 | 0,000 | 0,000 | 0,027 | 0,163 | 0,042 | 0,402 | 0,331 | 0,260 | 0,537 | 0,399 | 0,666 | 0,636 | 0,259 | 0,734 | 0,382 | 0,686 |
| Cgi-TLRδ2 | 0,877 | 0,798 | 0,499 | 0,418 | 0,223 | 0,254 | 0,221 | 0,272 | 0,170 | 0,186 | 0,249 | 0,088 | 0,333 | 0,257 | 0,152 | 0,267 | 0,326 | 0,217 | 0,295 |

| kallisto | Early morula | Morula | Blastula | Rotary movement | Free swimming | Early gastrula | Gastrula | Trochophore 1 | Trochophore 2 | Trochophore 3 | Trochophore 4 | Trochophore 5 | early Dshaped larva 1 | early Dshaped larva 2 | Dshaped larva 1 | Dshaped larva 2 | Dshaped larva 3 | Dshaped larva 4 | Dshaped larva 5 |
|---|---|---|---|---|---|---|---|---|---|---|---|---|---|---|---|---|---|---|---|
| Cgi-TLRα1 | 0,000 | 0,000 | 0,141 | 1,011 | 0,890 | 0,187 | 0,671 | 0,100 | 1,331 | 0,772 | 0,302 | 1,031 | 0,821 | 1,193 | 0,820 | 0,457 | 0,472 | 0,512 | 0,384 |
| Cgi-TLRα2 | 0,058 | 0,137 | 0,279 | 0,143 | 0,068 | 0,062 | 0,133 | 0,099 | 0,000 | 0,127 | 0,149 | 0,127 | 0,135 | 0,000 | 0,000 | 0,000 | 0,000 | 0,127 | |
| Cgi-TLRα3 | 0,000 | 0,000 | 0,000 | 0,000 | 0,000 | 0,000 | 0,000 | 0,000 | 0,136 | 0,262 | 0,461 | 0,131 | 0,139 | 0,608 | 0,596 | 0,816 | 0,120 | 0,653 | 0,392 |
| Cgi-TLRα4 | 0,000 | 0,000 | 0,000 | 0,190 | 0,541 | 0,737 | 0,353 | 0,262 | 0,701 | 0,508 | 0,794 | 1,017 | 1,441 | 1,413 | 1,694 | 0,752 | 1,709 | 2,865 | 4,888 |
| Cgi-TLRβ1 | 0,000 | 0,000 | 0,000 | 0,000 | 0,000 | 0,000 | 0,000 | 0,000 | 0,119 | 0,115 | 0,000 | 0,000 | 0,000 | 0,107 | 0,000 | 0,205 | 0,000 | 0,115 | 0,344 |
| Cgi-TLRβ2 | 0,049 | 0,000 | 0,000 | 0,000 | 0,058 | 0,000 | 0,000 | 0,000 | 0,000 | 0,109 | 0,000 | 0,000 | 0,000 | 0,000 | 0,199 | 0,000 | 0,100 | 0,000 | 0,326 |
| Cgi-TLRβ3 | 0,000 | 0,209 | 0,068 | 0,270 | 0,000 | 0,000 | 0,000 | 0,000 | 0,000 | 0,000 | 0,147 | 0,000 | 0,000 | 0,000 | 0,114 | 0,000 | 0,115 | 0,000 | 0,124 |
| Cgi-TLRβ4 | 2,735 | 2,132 | 2,365 | 2,237 | 1,301 | 1,388 | 2,383 | 1,783 | 1,924 | 2,013 | 1,517 | 1,440 | 0,865 | 1,734 | 0,615 | 1,150 | 0,924 | 1,002 | 1,002 |
| Cgi-TLRγ1 | 0,000 | 0,000 | 0,000 | 0,000 | 0,000 | 0,000 | 0,000 | 0,000 | 0,000 | 0,502 | 0,118 | 0,000 | 0,214 | 0,186 | 0,091 | 0,089 | 0,092 | 0,100 | 0,000 |
| Cgi-TLRγ2 | 0,000 | 0,000 | 0,000 | 0,000 | 0,000 | 0,000 | 0,000 | 0,150 | 0,200 | 0,000 | 0,000 | 0,193 | 0,000 | 0,179 | 0,352 | 0,343 | 0,000 | 0,192 | 0,000 |
| Cgi-TLRδ1 | 0,000 | 0,000 | 0,000 | 0,000 | 0,000 | 0,000 | 0,267 | 0,000 | 0,794 | 0,768 | 0,300 | 1,025 | 0,545 | 1,424 | 1,048 | 0,455 | 1,527 | 0,765 | 1,402 |
| Cgi-TLRδ2 | 1,543 | 1,722 | 0,993 | 0,977 | 0,618 | 0,491 | 0,227 | 0,506 | 0,300 | 0,290 | 0,595 | 0,363 | 0,540 | 0,538 | 0,264 | 0,451 | 0,532 | 0,433 | 0,650 |

TEM ≥0,150

TEM <0,150

**Supplementary Table 6** - *Terebratalia transversa* stage specific transcriptome analyses. Analyses for the different methods (RSEM and Kallisto) and replicates (Rep 1_and Rep_2). For each method, average and standard error (SE) of the two replicates is provided.

*TEREBRATALIA TRANSVERSA*

**Values inicate Transcripts per Milion (TEM)**

| Rep_1_RSEM | oocyte | 8 hr mid blastula | 19 hr late blastula | 24 hr moving blastula | 26 hr early gastrula | 37 hr mid gastrula | 51 hr late gastrula | 59 hr bilobed gastrula | 68 hr trilobed gastrula | 82 hr early larva | 98 hr late larva | competent larva (131h) | 1 day juvenile | 2 day juvenile |
|---|---|---|---|---|---|---|---|---|---|---|---|---|---|---|
| **Ttr-TLRα1** | 0,000 | 0,000 | 0,000 | 0,000 | 0,000 | 0,000 | 0,000 | 0,000 | 0,000 | 0,000 | 0,000 | 0,000 | 0,000 | 0,000 |
| **Ttr-TLRα2** | 0,290 | 0,216 | 0,189 | 0,104 | 0,058 | 0,028 | 0,028 | 0,064 | 0,116 | 0,139 | 0,062 | 1,570 | 0,359 | 0,600 |
| **Ttr-TLRα3** | 0,000 | 0,000 | 0,000 | 0,088 | 0,099 | 0,000 | 0,000 | 0,000 | 0,000 | 0,000 | 0,000 | 2,302 | 1,983 | 2,589 |
| **Ttr-TLRα4** | 0,807 | 0,059 | 0,074 | 0,249 | 0,214 | 0,047 | 0,000 | 0,000 | 0,412 | 0,492 | 1,123 | 2,691 | 4,488 | 2,578 |
| **Ttr-TLRα5** | 0,105 | 0,137 | 0,041 | 0,064 | 0,074 | 0,056 | 0,120 | 0,100 | 0,245 | 0,177 | 0,072 | 0,071 | 0,098 | 0,235 |
| **Ttr-TLRβ1** | 0,129 | 0,000 | 0,000 | 0,096 | 0,222 | 0,150 | 0,000 | 0,000 | 0,039 | 0,253 | 0,577 | 0,708 | 0,153 | 0,000 |
| **Ttr-TLRβ2** | 0,524 | 0,634 | 0,287 | 0,329 | 0,330 | 0,122 | 0,240 | 0,237 | 0,155 | 0,164 | 0,546 | 0,342 | 0,065 | 0,071 |
| **Ttr-TLRβ3** | 0,331 | 0,471 | 0,771 | 0,553 | 0,577 | 0,207 | 0,065 | 0,428 | 0,142 | 0,076 | 0,330 | 0,224 | 0,458 | 0,271 |
| **Ttr-TLRβ4** | 0,000 | 0,000 | 0,090 | 0,048 | 0,346 | 0,197 | 0,037 | 0,391 | 0,567 | 0,076 | 0,824 | 0,602 | 0,381 | 0,294 |
| **Ttr-TLRβ5** | 0,000 | 0,039 | 0,066 | 0,160 | 0,058 | 0,066 | 0,000 | 0,000 | 0,000 | 0,000 | 0,000 | 0,059 | 0,000 | 0,000 |
| **Ttr-TLRγ1** | 0,000 | 0,000 | 0,000 | 0,000 | 0,000 | 0,000 | 0,092 | 0,064 | 0,000 | 0,000 | 0,000 | 0,000 | 0,501 | 0,094 |
| **Ttr-TLRγ2** | 0,000 | 0,000 | 0,000 | 0,000 | 0,000 | 0,000 | 0,000 | 0,000 | 0,000 | 0,000 | 0,000 | 0,106 | 0,000 | 0,282 |
| **Ttr-TLRγ3** | 0,000 | 0,000 | 0,000 | 0,000 | 0,000 | 0,000 | 0,000 | 0,000 | 0,000 | 0,000 | 0,000 | 0,000 | 0,000 | 0,000 |
| **Ttr-TLRγ4** | 0,226 | 0,275 | 0,082 | 0,192 | 0,568 | 0,442 | 0,406 | 0,309 | 0,142 | 0,088 | 0,227 | 0,236 | 0,458 | 0,294 |
| **Ttr-TLRδ** | 0,024 | 0,020 | 0,057 | 0,152 | 0,148 | 0,160 | 0,000 | 0,064 | 0,052 | 0,126 | 0,165 | 0,224 | 0,033 | 0,000 |

| Rep_2_RSEM | oocyte | 8 hr mid blastula | 19 hr late blastula | 24 hr moving blastula | 26 hr early gastrula | 37 hr mid gastrula | 51 hr late gastrula | 59 hr bilobed gastrula | 68 hr trilobed gastrula | 82 hr early larva | 98 hr late larva | competent larva (131h) | 1 day juvenile | 2 day juvenile |
|---|---|---|---|---|---|---|---|---|---|---|---|---|---|---|
| **Ttr-TLRα1** | 0,000 | 0,000 | 0,000 | 0,000 | 0,000 | 0,000 | 0,000 | 0,000 | 0,000 | 0,000 | 0,000 | 0,000 | 0,000 | 0,000 |
| **Ttr-TLRα2** | 0,121 | 0,161 | 0,207 | 0,222 | 0,080 | 0,116 | 0,034 | 0,035 | 0,157 | 0,343 | 0,229 | 0,668 | 0,639 | 0,403 |
| **Ttr-TLRα3** | 0,000 | 0,000 | 0,057 | 0,000 | 0,000 | 0,045 | 0,000 | 0,000 | 0,000 | 0,000 | 0,000 | 0,196 | 1,943 | 1,327 |
| **Ttr-TLRα4** | 0,664 | 0,484 | 0,607 | 0,625 | 1,045 | 0,695 | 0,051 | 0,129 | 0,868 | 0,520 | 0,774 | 0,701 | 3,791 | 5,293 |
| **Ttr-TLRα5** | 0,121 | 0,101 | 0,136 | 0,277 | 0,195 | 0,196 | 0,488 | 0,493 | 1,129 | 0,213 | 0,229 | 0,163 | 0,326 | 0,522 |
| **Ttr-TLRβ1** | 0,113 | 0,000 | 0,064 | 0,000 | 0,000 | 0,062 | 0,000 | 0,000 | 0,220 | 0,106 | 0,264 | 0,342 | 0,163 | 0,030 |
| **Ttr-TLRβ2** | 0,089 | 0,040 | 0,236 | 0,087 | 0,062 | 0,018 | 0,069 | 0,082 | 0,366 | 0,095 | 0,035 | 0,065 | 0,122 | 0,000 |
| **Ttr-TLRβ3** | 0,810 | 0,471 | 0,743 | 0,688 | 0,789 | 0,740 | 0,950 | 0,258 | 1,098 | 0,118 | 0,211 | 0,261 | 0,353 | 0,268 |
| **Ttr-TLRβ4** | 0,000 | 0,000 | 0,043 | 0,111 | 0,000 | 0,000 | 0,000 | 0,000 | 0,063 | 0,000 | 0,000 | 0,000 | 0,000 | 0,000 |
| **Ttr-TLRβ5** | 0,000 | 0,000 | 0,036 | 0,222 | 0,000 | 0,000 | 0,077 | 0,000 | 0,000 | 0,000 | 0,000 | 0,000 | 0,000 | 0,119 |
| **Ttr-TLRγ1** | 0,000 | 0,034 | 0,000 | 0,000 | 0,000 | 0,053 | 0,103 | 0,000 | 0,000 | 0,000 | 0,000 | 0,000 | 0,000 | 0,000 |
| **Ttr-TLRγ2** | 0,000 | 0,000 | 0,036 | 0,000 | 0,000 | 0,000 | 0,000 | 0,000 | 0,209 | 0,000 | 0,000 | 0,000 | 0,000 | 0,000 |
| **Ttr-TLRγ3** | 0,000 | 0,000 | 0,000 | 0,000 | 0,000 | 0,027 | 0,000 | 0,000 | 0,000 | 0,000 | 0,000 | 0,000 | 0,000 | 0,000 |
| **Ttr-TLRγ4** | 0,121 | 0,195 | 0,129 | 0,182 | 0,106 | 0,134 | 0,325 | 0,317 | 0,324 | 0,154 | 0,457 | 0,326 | 0,476 | 0,656 |
| **Ttr-TLRδ** | 0,000 | 0,040 | 0,279 | 0,649 | 0,230 | 0,160 | 0,394 | 0,047 | 0,178 | 0,035 | 0,070 | 0,065 | 0,190 | 0,313 |

| RSEM_average | oocyte | 8 hr mid blastula | 19 hr late blastula | 24 hr moving blastula | 26 hr early gastrula | 37 hr mid gastrula | 51 hr late gastrula | 59 hr bilobed gastrula | 68 hr trilobed gastrula | 82 hr early larva | 98 hr late larva | competent larva (131h) | 1 day juvenile | 2 day juvenile |
|---|---|---|---|---|---|---|---|---|---|---|---|---|---|---|
| **Ttr-TLRα1** | 0,000 | 0,000 | 0,000 | 0,000 | 0,000 | 0,000 | 0,000 | 0,000 | 0,000 | 0,000 | 0,000 | 0,000 | 0,000 | 0,000 |
| **Ttr-TLRα2** | 0,206 | 0,189 | 0,198 | 0,163 | 0,069 | 0,072 | 0,031 | 0,050 | 0,137 | 0,241 | 0,146 | 1,119 | 0,499 | 0,502 |
| **Ttr-TLRα3** | 0,000 | 0,000 | 0,029 | 0,044 | 0,050 | 0,023 | 0,000 | 0,000 | 0,000 | 0,000 | 0,000 | 1,249 | 1,963 | 1,958 |
| **Ttr-TLRα4** | 0,736 | 0,272 | 0,341 | 0,437 | 0,630 | 0,371 | 0,026 | 0,065 | 0,640 | 0,506 | 0,949 | 1,696 | 4,140 | 3,936 |
| **Ttr-TLRα5** | 0,113 | 0,119 | 0,089 | 0,171 | 0,135 | 0,126 | 0,304 | 0,297 | 0,687 | 0,195 | 0,151 | 0,117 | 0,212 | 0,379 |
| **Ttr-TLRβ1** | 0,121 | 0,000 | 0,032 | 0,048 | 0,111 | 0,106 | 0,000 | 0,000 | 0,130 | 0,180 | 0,421 | 0,525 | 0,158 | 0,015 |
| **Ttr-TLRβ2** | 0,307 | 0,337 | 0,262 | 0,208 | 0,196 | 0,070 | 0,155 | 0,160 | 0,261 | 0,130 | 0,291 | 0,204 | 0,094 | 0,036 |
| **Ttr-TLRβ3** | 0,571 | 0,471 | 0,757 | 0,621 | 0,683 | 0,474 | 0,508 | 0,343 | 0,620 | 0,097 | 0,271 | 0,243 | 0,406 | 0,270 |
| **Ttr-TLRβ4** | 0,000 | 0,000 | 0,067 | 0,080 | 0,173 | 0,099 | 0,019 | 0,196 | 0,315 | 0,038 | 0,412 | 0,301 | 0,191 | 0,147 |
| **Ttr-TLRβ5** | 0,000 | 0,020 | 0,051 | 0,191 | 0,029 | 0,033 | 0,039 | 0,000 | 0,000 | 0,000 | 0,000 | 0,030 | 0,000 | 0,060 |
| **Ttr-TLRγ1** | 0,000 | 0,017 | 0,000 | 0,000 | 0,000 | 0,027 | 0,098 | 0,032 | 0,000 | 0,000 | 0,000 | 0,000 | 0,251 | 0,047 |
| **Ttr-TLRγ2** | 0,000 | 0,000 | 0,018 | 0,000 | 0,000 | 0,000 | 0,000 | 0,000 | 0,105 | 0,000 | 0,000 | 0,053 | 0,000 | 0,141 |
| **Ttr-TLRγ3** | 0,000 | 0,000 | 0,000 | 0,000 | 0,000 | 0,014 | 0,000 | 0,000 | 0,000 | 0,000 | 0,000 | 0,000 | 0,000 | 0,000 |
| **Ttr-TLRγ4** | 0,174 | 0,235 | 0,106 | 0,187 | 0,337 | 0,288 | 0,366 | 0,313 | 0,233 | 0,121 | 0,342 | 0,281 | 0,467 | 0,475 |
| **Ttr-TLRδ** | 0,012 | 0,030 | 0,168 | 0,401 | 0,189 | 0,160 | 0,197 | 0,056 | 0,115 | 0,081 | 0,118 | 0,145 | 0,112 | 0,157 |

TEM ≥0,150

TEM <0,150

**Values indicate Standard Error (SE)**

| RSEM_SE | oocyte | 8 hr mid blastula | 19 hr late blastula | 24 hr moving blastula | 26 hr early gastrula | 37 hr mid gastrula | 51 hr late gastrula | 59 hr bilobed gastrula | 68 hr trilobed gastrula | 82 hr early larva | 98 hr late larva | competent larva (131h) | 1 day juvenile | 2 day juvenile |
|---|---|---|---|---|---|---|---|---|---|---|---|---|---|---|
| **Ttr-TLRα1** | 0,000 | 0,000 | 0,000 | 0,000 | 0,000 | 0,000 | 0,000 | 0,000 | 0,000 | 0,000 | 0,000 | 0,000 | 0,000 | 0,000 |
| **Ttr-TLRα2** | 0,085 | 0,028 | 0,009 | 0,059 | 0,011 | 0,044 | 0,003 | 0,015 | 0,020 | 0,102 | 0,084 | 0,451 | 0,140 | 0,098 |
| **Ttr-TLRα3** | 0,000 | 0,000 | 0,029 | 0,044 | 0,050 | 0,023 | 0,000 | 0,000 | 0,000 | 0,000 | 0,000 | 1,053 | 0,020 | 0,631 |
| **Ttr-TLRα4** | 0,072 | 0,213 | 0,267 | 0,188 | 0,416 | 0,324 | 0,026 | 0,065 | 0,228 | 0,014 | 0,175 | 0,995 | 0,349 | 1,358 |
| **Ttr-TLRα5** | 0,008 | 0,018 | 0,048 | 0,107 | 0,061 | 0,070 | 0,184 | 0,197 | 0,442 | 0,018 | 0,079 | 0,046 | 0,114 | 0,144 |
| **Ttr-TLRβ1** | 0,008 | 0,000 | 0,032 | 0,048 | 0,111 | 0,044 | 0,000 | 0,000 | 0,091 | 0,074 | 0,157 | 0,183 | 0,005 | 0,015 |
| **Ttr-TLRβ2** | 0,218 | 0,297 | 0,026 | 0,121 | 0,134 | 0,052 | 0,086 | 0,078 | 0,106 | 0,035 | 0,256 | 0,139 | 0,029 | 0,036 |
| **Ttr-TLRβ3** | 0,240 | 0,000 | 0,014 | 0,067 | 0,106 | 0,267 | 0,443 | 0,085 | 0,478 | 0,021 | 0,059 | 0,019 | 0,053 | 0,002 |

| | | | | | | | | | | | | | | |
|---|---|---|---|---|---|---|---|---|---|---|---|---|---|---|
| **Ttr-TLRβ4** | 0,000 | 0,000 | 0,024 | 0,032 | 0,173 | 0,099 | 0,019 | 0,196 | 0,252 | 0,038 | 0,412 | 0,301 | 0,191 | 0,147 |
| **Ttr-TLRβ5** | 0,000 | 0,020 | 0,015 | 0,031 | 0,029 | 0,033 | 0,039 | 0,000 | 0,000 | 0,000 | 0,000 | 0,030 | 0,000 | 0,060 |
| **Ttr-TLRγ1** | 0,000 | 0,017 | 0,000 | 0,000 | 0,000 | 0,027 | 0,006 | 0,032 | 0,000 | 0,000 | 0,000 | 0,000 | 0,251 | 0,047 |
| **Ttr-TLRγ2** | 0,000 | 0,000 | 0,018 | 0,000 | 0,000 | 0,000 | 0,000 | 0,000 | 0,105 | 0,000 | 0,000 | 0,053 | 0,000 | 0,141 |
| **Ttr-TLRγ3** | 0,000 | 0,000 | 0,000 | 0,000 | 0,000 | 0,014 | 0,000 | 0,000 | 0,000 | 0,000 | 0,000 | 0,000 | 0,000 | 0,000 |
| **Ttr-TLRγ4** | 0,053 | 0,040 | 0,023 | 0,005 | 0,231 | 0,154 | 0,040 | 0,004 | 0,091 | 0,033 | 0,115 | 0,045 | 0,009 | 0,181 |
| **Ttr-TLRδ** | 0,012 | 0,010 | 0,111 | 0,249 | 0,041 | 0,000 | 0,197 | 0,008 | 0,063 | 0,046 | 0,048 | 0,080 | 0,079 | 0,157 |

| **Rep_1_kallisto** | **oocyte** | **8 hr mid blastula** | **19 hr late blastula** | **24 hr moving blastula** | **26 hr early gastrula** | **37 hr mid gastrula** | **51 hr late gastrula** | **59 hr bilobed gastrula** | **68 hr trilobed gastrula** | **82 hr early larva** | **98 hr late larva** | **competent larva (131h)** | **1 day juvenile** | **2 day juvenile** |
|---|---|---|---|---|---|---|---|---|---|---|---|---|---|---|
| **Ttr-TLRα1** | 0,000 | 0,000 | 0,000 | 0,000 | 0,000 | 0,000 | 0,000 | 0,000 | 0,000 | 0,000 | 0,006 | 0,000 | 0,000 | 0,000 |
| **Ttr-TLRα2** | 0,300 | 0,139 | 0,162 | 0,084 | 0,059 | 0,034 | 0,032 | 0,063 | 0,127 | 0,114 | 0,031 | 1,694 | 0,342 | 0,638 |
| **Ttr-TLRα3** | 0,000 | 0,000 | 0,000 | 0,000 | 0,068 | 0,000 | 0,000 | 0,000 | 0,000 | 0,000 | 0,042 | 2,292 | 1,974 | 2,754 |
| **Ttr-TLRα4** | 0,726 | 0,074 | 0,125 | 0,249 | 0,224 | 0,043 | 0,000 | 0,000 | 0,335 | 0,481 | 0,913 | 2,408 | 3,993 | 2,758 |
| **Ttr-TLRα5** | 0,076 | 0,109 | 0,020 | 0,064 | 0,054 | 0,061 | 0,119 | 0,096 | 0,194 | 0,186 | 0,056 | 0,107 | 0,057 | 0,225 |
| **Ttr-TLRβ1** | 0,099 | 0,101 | 0,001 | 0,095 | 0,004 | 0,000 | 0,000 | 0,000 | 0,000 | 0,237 | 0,000 | 0,628 | 0,004 | 0,008 |
| **Ttr-TLRβ2** | 0,490 | 0,517 | 0,228 | 0,301 | 0,248 | 0,071 | 0,137 | 0,172 | 0,089 | 0,133 | 0,560 | 0,312 | 0,065 | 0,064 |
| **Ttr-TLRβ3** | 0,313 | 0,398 | 0,737 | 0,463 | 0,360 | 0,185 | 0,089 | 0,408 | 0,145 | 0,000 | 0,252 | 0,252 | 0,517 | 0,225 |
| **Ttr-TLRβ4** | 0,000 | 0,038 | 0,000 | 0,062 | 0,371 | 0,460 | 0,063 | 0,624 | 0,630 | 0,408 | 0,767 | 0,568 | 0,285 | 0,309 |
| **Ttr-TLRβ5** | 0,000 | 0,045 | 0,000 | 0,140 | 0,066 | 0,075 | 0,000 | 0,000 | 0,000 | 0,000 | 0,000 | 0,065 | 0,000 | 0,000 |
| **Ttr-TLRγ1** | 0,000 | 0,000 | 0,127 | 0,041 | 0,249 | 0,144 | 0,316 | 0,061 | 0,000 | 0,000 | 0,000 | 0,057 | 0,071 | 0,136 |
| **Ttr-TLRγ2** | 0,000 | 0,000 | 0,000 | 0,000 | 0,000 | 0,000 | 0,000 | 0,000 | 0,000 | 0,000 | 0,000 | 0,000 | 0,000 | 0,188 |
| **Ttr-TLRγ3** | 0,000 | 0,000 | 0,000 | 0,000 | 0,000 | 0,000 | 0,000 | 0,000 | 0,000 | 0,000 | 0,000 | 0,000 | 0,000 | 0,000 |
| **Ttr-TLRγ4** | 0,211 | 0,228 | 0,039 | 0,176 | 0,551 | 0,404 | 0,371 | 0,283 | 0,115 | 0,092 | 0,204 | 0,176 | 0,393 | 0,304 |
| **Ttr-TLRδ** | 0,000 | 0,020 | 0,038 | 0,135 | 0,200 | 0,192 | 0,000 | 0,042 | 0,079 | 0,209 | 0,183 | 0,281 | 0,000 | 0,000 |

| **Rep_2_kallisto** | **oocyte** | **8 hr mid blastula** | **19 hr late blastula** | **24 hr moving blastula** | **26 hr early gastrula** | **37 hr mid gastrula** | **51 hr late gastrula** | **59 hr bilobed gastrula** | **68 hr trilobed gastrula** | **82 hr early larva** | **98 hr late larva** | **competent larva (131h)** | **1 day juvenile** | **2 day juvenile** |
|---|---|---|---|---|---|---|---|---|---|---|---|---|---|---|
| **Ttr-TLRα1** | 0,000 | 0,000 | 0,000 | 0,019 | 0,000 | 0,000 | 0,000 | 0,000 | 0,000 | 0,000 | 0,021 | 0,027 | 0,000 | 0,000 |
| **Ttr-TLRα2** | 0,143 | 0,165 | 0,184 | 0,150 | 0,083 | 0,120 | 0,037 | 0,037 | 0,157 | 0,364 | 0,194 | 0,578 | 0,690 | 0,443 |
| **Ttr-TLRα3** | 0,000 | 0,000 | 0,058 | 0,000 | 0,000 | 0,000 | 0,105 | 0,000 | 0,000 | 0,000 | 0,000 | 0,167 | 2,062 | 1,534 |
| **Ttr-TLRα4** | 0,466 | 0,403 | 0,583 | 0,849 | 0,905 | 0,667 | 0,129 | 0,214 | 0,865 | 0,347 | 0,730 | 0,630 | 3,468 | 5,164 |
| **Ttr-TLRα5** | 0,116 | 0,090 | 0,133 | 0,252 | 0,186 | 0,159 | 0,430 | 0,401 | 1,017 | 0,222 | 0,207 | 0,147 | 0,312 | 0,618 |
| **Ttr-TLRβ1** | 0,094 | 0,001 | 0,047 | 0,003 | 0,001 | 0,045 | 0,000 | 0,000 | 0,226 | 0,001 | 0,192 | 0,354 | 0,130 | 0,000 |
| **Ttr-TLRβ2** | 0,082 | 0,025 | 0,235 | 0,079 | 0,058 | 0,014 | 0,065 | 0,076 | 0,314 | 0,116 | 0,034 | 0,069 | 0,140 | 0,086 |
| **Ttr-TLRβ3** | 0,626 | 0,416 | 0,633 | 0,515 | 0,671 | 0,562 | 0,778 | 0,221 | 0,927 | 0,086 | 0,076 | 0,183 | 0,199 | 0,000 |
| **Ttr-TLRβ4** | 0,000 | 0,000 | 0,060 | 0,000 | 0,000 | 0,119 | 0,000 | 0,000 | 0,172 | 0,071 | 0,000 | 0,170 | 0,080 | 0,000 |
| **Ttr-TLRβ5** | 0,000 | 0,000 | 0,038 | 0,253 | 0,000 | 0,000 | 0,042 | 0,000 | 0,000 | 0,000 | 0,000 | 0,000 | 0,000 | 0,000 |
| **Ttr-TLRγ1** | 0,000 | 0,000 | 0,033 | 0,000 | 0,000 | 0,088 | 0,145 | 0,000 | 0,000 | 0,000 | 0,000 | 0,000 | 0,000 | 0,123 |
| **Ttr-TLRγ2** | 0,000 | 0,000 | 0,000 | 0,000 | 0,000 | 0,000 | 0,000 | 0,000 | 0,174 | 0,000 | 0,000 | 0,000 | 0,000 | 0,000 |
| **Ttr-TLRγ3** | 0,000 | 0,000 | 0,000 | 0,000 | 0,000 | 0,000 | 0,000 | 0,000 | 0,000 | 0,000 | 0,000 | 0,000 | 0,000 | 0,000 |
| **Ttr-TLRγ4** | 0,100 | 0,166 | 0,131 | 0,113 | 0,100 | 0,109 | 0,279 | 0,264 | 0,256 | 0,159 | 0,321 | 0,348 | 0,477 | 0,572 |
| **Ttr-TLRδ** | 0,000 | 0,026 | 0,255 | 0,657 | 0,327 | 0,120 | 0,512 | 0,091 | 0,460 | 0,000 | 0,054 | 0,216 | 0,201 | 0,287 |

| **Kallisto_average** | **oocyte** | **8 hr mid blastula** | **19 hr late blastula** | **24 hr moving blastula** | **26 hr early gastrula** | **37 hr mid gastrula** | **51 hr late gastrula** | **59 hr bilobed gastrula** | **68 hr trilobed gastrula** | **82 hr early larva** | **98 hr late larva** | **competent larva (131h)** | **1 day juvenile** | **2 day juvenile** |
|---|---|---|---|---|---|---|---|---|---|---|---|---|---|---|
| **Ttr-TLRα1** | 0,000 | 0,000 | 0,000 | 0,010 | 0,000 | 0,000 | 0,000 | 0,000 | 0,000 | 0,000 | 0,014 | 0,014 | 0,000 | 0,000 |
| **Ttr-TLRα2** | 0,222 | 0,152 | 0,173 | 0,117 | 0,071 | 0,077 | 0,035 | 0,050 | 0,142 | 0,239 | 0,113 | 1,136 | 0,516 | 0,541 |
| **Ttr-TLRα3** | 0,000 | 0,000 | 0,029 | 0,000 | 0,034 | 0,000 | 0,053 | 0,000 | 0,000 | 0,000 | 0,021 | 1,230 | 2,018 | 2,144 |
| **Ttr-TLRα4** | 0,596 | 0,239 | 0,354 | 0,549 | 0,565 | 0,355 | 0,065 | 0,107 | 0,600 | 0,414 | 0,822 | 1,519 | 3,731 | 3,961 |
| **Ttr-TLRα5** | 0,096 | 0,100 | 0,077 | 0,158 | 0,120 | 0,110 | 0,275 | 0,249 | 0,606 | 0,204 | 0,132 | 0,127 | 0,185 | 0,422 |
| **Ttr-TLRβ1** | 0,097 | 0,051 | 0,024 | 0,049 | 0,003 | 0,023 | 0,000 | 0,000 | 0,113 | 0,119 | 0,096 | 0,491 | 0,067 | 0,004 |
| **Ttr-TLRβ2** | 0,286 | 0,271 | 0,232 | 0,190 | 0,153 | 0,043 | 0,101 | 0,124 | 0,202 | 0,125 | 0,297 | 0,191 | 0,103 | 0,075 |
| **Ttr-TLRβ3** | 0,470 | 0,407 | 0,685 | 0,489 | 0,516 | 0,374 | 0,434 | 0,315 | 0,536 | 0,043 | 0,164 | 0,218 | 0,358 | 0,113 |
| **Ttr-TLRβ4** | 0,000 | 0,019 | 0,030 | 0,031 | 0,186 | 0,290 | 0,032 | 0,312 | 0,401 | 0,240 | 0,384 | 0,369 | 0,183 | 0,155 |
| **Ttr-TLRβ5** | 0,000 | 0,023 | 0,019 | 0,197 | 0,033 | 0,038 | 0,021 | 0,000 | 0,000 | 0,000 | 0,000 | 0,033 | 0,000 | 0,000 |
| **Ttr-TLRγ1** | 0,000 | 0,000 | 0,080 | 0,021 | 0,125 | 0,116 | 0,231 | 0,031 | 0,000 | 0,000 | 0,000 | 0,029 | 0,036 | 0,130 |
| **Ttr-TLRγ2** | 0,000 | 0,000 | 0,000 | 0,000 | 0,000 | 0,000 | 0,000 | 0,000 | 0,087 | 0,000 | 0,000 | 0,000 | 0,000 | 0,094 |
| **Ttr-TLRγ3** | 0,000 | 0,000 | 0,000 | 0,000 | 0,000 | 0,000 | 0,000 | 0,000 | 0,000 | 0,000 | 0,000 | 0,000 | 0,000 | 0,000 |
| **Ttr-TLRγ4** | 0,156 | 0,197 | 0,085 | 0,145 | 0,326 | 0,257 | 0,325 | 0,274 | 0,186 | 0,126 | 0,263 | 0,262 | 0,435 | 0,438 |
| **Ttr-TLRδ** | 0,000 | 0,023 | 0,147 | 0,396 | 0,264 | 0,156 | 0,256 | 0,067 | 0,270 | 0,105 | 0,119 | 0,249 | 0,101 | 0,144 |

TEM ≥0,150
TEM <0,150

| **RSEM_Standard error** | **oocyte** | **8 hr mid blastula** | **19 hr late blastula** | **24 hr moving blastula** | **26 hr early gastrula** | **37 hr mid gastrula** | **51 hr late gastrula** | **59 hr bilobed gastrula** | **68 hr trilobed gastrula** | **82 hr early larva** | **98 hr late larva** | **competent larva (131h)** | **1 day juvenile** | **2 day juvenile** |
|---|---|---|---|---|---|---|---|---|---|---|---|---|---|---|
| **Ttr-TLRα1** | 0,000 | 0,000 | 0,000 | 0,010 | 0,000 | 0,000 | 0,000 | 0,000 | 0,000 | 0,000 | 0,008 | 0,014 | 0,000 | 0,000 |
| **Ttr-TLRα2** | 0,079 | 0,013 | 0,011 | 0,033 | 0,012 | 0,043 | 0,003 | 0,013 | 0,015 | 0,125 | 0,082 | 0,558 | 0,174 | 0,098 |
| **Ttr-TLRα3** | 0,000 | 0,000 | 0,029 | 0,000 | 0,034 | 0,000 | 0,053 | 0,000 | 0,000 | 0,000 | 0,021 | 1,063 | 0,044 | 0,610 |
| **Ttr-TLRα4** | 0,130 | 0,165 | 0,229 | 0,300 | 0,341 | 0,312 | 0,065 | 0,107 | 0,265 | 0,067 | 0,091 | 0,889 | 0,263 | 1,203 |
| **Ttr-TLRα5** | 0,020 | 0,010 | 0,057 | 0,094 | 0,066 | 0,049 | 0,156 | 0,153 | 0,412 | 0,018 | 0,076 | 0,020 | 0,128 | 0,197 |
| **Ttr-TLRβ1** | 0,003 | 0,050 | 0,023 | 0,046 | 0,002 | 0,023 | 0,000 | 0,000 | 0,113 | 0,118 | 0,096 | 0,137 | 0,063 | 0,004 |
| **Ttr-TLRβ2** | 0,204 | 0,246 | 0,003 | 0,111 | 0,095 | 0,029 | 0,036 | 0,048 | 0,113 | 0,009 | 0,263 | 0,122 | 0,038 | 0,011 |
| **Ttr-TLRβ3** | 0,157 | 0,009 | 0,052 | 0,026 | 0,156 | 0,189 | 0,345 | 0,093 | 0,391 | 0,043 | 0,088 | 0,035 | 0,159 | 0,113 |
| **Ttr-TLRβ4** | 0,000 | 0,019 | 0,030 | 0,031 | 0,186 | 0,171 | 0,032 | 0,312 | 0,229 | 0,169 | 0,384 | 0,199 | 0,103 | 0,155 |
| **Ttr-TLRβ5** | 0,000 | 0,023 | 0,019 | 0,057 | 0,033 | 0,038 | 0,021 | 0,000 | 0,000 | 0,000 | 0,000 | 0,033 | 0,000 | 0,000 |
| **Ttr-TLRγ1** | 0,000 | 0,000 | 0,047 | 0,021 | 0,125 | 0,028 | 0,086 | 0,031 | 0,000 | 0,000 | 0,000 | 0,029 | 0,036 | 0,007 |

| | | | | | | | | | | | | | | |
|---|---|---|---|---|---|---|---|---|---|---|---|---|---|---|
| **Ttr-TLRγ2** | 0,000 | 0,000 | 0,000 | 0,000 | 0,000 | 0,000 | 0,000 | 0,000 | 0,087 | 0,000 | 0,000 | 0,000 | 0,000 | 0,094 |
| **Ttr-TLRγ3** | 0,000 | 0,000 | 0,000 | 0,000 | 0,000 | 0,000 | 0,000 | 0,000 | 0,000 | 0,000 | 0,000 | 0,000 | 0,000 | 0,000 |
| **Ttr-TLRγ4** | 0,056 | 0,031 | 0,046 | 0,032 | 0,226 | 0,148 | 0,046 | 0,009 | 0,071 | 0,034 | 0,059 | 0,086 | 0,042 | 0,134 |
| **Ttr-TLRδ** | 0,000 | 0,003 | 0,109 | 0,261 | 0,064 | 0,036 | 0,256 | 0,025 | 0,191 | 0,105 | 0,065 | 0,033 | 0,101 | 0,144 |
